# Long-Term Effects of a Novel Continuous Remote Care Intervention Including Nutritional Ketosis for the Management of Type 2 Diabetes: A 2-year Non-randomized Clinical Trial

**DOI:** 10.1101/476275

**Authors:** Shaminie J. Athinarayanan, Rebecca N. Adams, Sarah J. Hallberg, Amy L. McKenzie, Nasir H. Bhanpuri, Wayne W. Campbell, Jeff S. Volek, Stephen D. Phinney, James P. McCarter

## Abstract

**OBJECTIVE:** Studies on long-term sustainability of low-carbohydrate approaches to treat diabetes are limited. We aim to assess the effects of a continuous care intervention (CCI) on retention, glycemic control, weight, body composition, cardiovascular, liver, kidney, thyroid, inflammatory markers, diabetes medication usage and disease outcomes at 2 years in adults with type 2 diabetes (T2D).

**RESEARCH DESIGN AND METHODS:** An open label, non-randomized, controlled study with 262 and 87 participants with T2D were enrolled in the CCI and usual care (UC) groups, respectively.

**RESULTS:** Significant changes from baseline to 2 years in the CCI group included: HbA1c (−12% from 7.7±0.1%); fasting glucose (−18% from 163.67±3.90 mg/dL); fasting insulin (−42% from 27.73±1.26 pmol L^-1^); weight (−10% from 114.56±0.60 kg); systolic blood pressure (−4% from 131.7±0.9 mmHg); diastolic blood pressure (−4% from 81.8±0.5 mmHg); triglycerides (−22% from 197.2±9.1 mg/dL); HDL-C (+19% from 41.8±0.9 mg/dL), and liver alanine transaminase (−21% from 29.16±0.97 U/L). Spine bone mineral density in the CCI group was unchanged. Glycemic control medication use (excluding metformin) among CCI participants declined (from 56.9% to 26.8%, P=1.3×10^-11^) including prescribed insulin (−62%) and sulfonylureas (−100%). The UC group had no significant changes in these parameters (except uric acid and anion gap) or diabetes medication use. There was also significant resolution of diabetes (reversal, 53.5%; remission, 17.6%) in the CCI group but not in UC. All the reported improvements had p-values <0.00012.

**CONCLUSIONS:** The CCI sustained long-term beneficial effects on multiple clinical markers of diabetes and cardiometabolic health at 2 years while utilizing less medication. The intervention was also effective in the resolution of diabetes and visceral obesity, with no adverse effect on bone health.

**TRIAL REGISTRATION:** Clinicaltrials.gov NCT02519309

## Introduction

Type 2 diabetes (T2D), obesity, and metabolic disease impact over one billion people and present a challenge to public health and economic growth(1,S34). In the United States, over 30 million people have diabetes and it is a leading cause of morbidity and mortality, especially through increased cardiovascular disease (CVD)(2). The remission rate under usual care is 0.5 – 2%(3) while intensive lifestyle intervention resulted in remission rates (both partial and complete) of 11.5% and 9.2% at 1 and 2 years(4). When lifestyle intervention is insufficient, medications are indicated to manage the disease and slow progression.

When T2D care directed at disease reversal is successful, this includes achievement of restored metabolic health, glycemic control with reduced dependence on medication, and in some cases disease remission. Three non-pharmaceutical approaches have demonstrated high rates of at least temporary T2D diabetes reversal or remission: bariatric surgery, very low calorie diets (VLCD), and nutritional ketosis achieved through carbohydrate restriction(5,6,7). In controlled clinical trials, each approach has demonstrated improved glycemic control and CVD risk factors, reduced pharmaceutical dependence, and weight loss. The three approaches show a similar time-course with glycemic control preceding weight loss by weeks or months, suggesting potential overlap of mechanisms(8,S35,S36).

With bariatric surgery, up to 60% of patients demonstrate T2D remission at 1 year(9). Outcomes at two years and beyond indicate ~50% of patients can achieve ongoing diabetes remission(10,S37). The second Diabetes Surgery Summit recommended using bariatric surgery to treat T2D with support from worldwide medical and scientific societies(10), but both complications and cost limit its widespread use(11,S38). VLCDs providing <900 kcal/day allow rapid discontinuation of most medications, improved glycemic control, and weight loss. This approach is necessarily temporary, however, with weight regain and impaired glucose control typically occurring within 3-6 months of reintroduction of substantial proportions of dietary carbohydrate (6,12,S39,S40).

A third approach to diabetes reversal is sustained dietary carbohydrate restriction. Low-carbohydrate diets have consistently elicited improvements in T2D, metabolic disease, and obesity up to one year(13,S41); however, longer-term studies and studies including patients prescribed insulin are limited. A low carbohydrate Mediterranean diet caused remission in 14.7% of newly diagnosed diabetes patients at 1 year versus 4.1% with a low-fat diet (14), and a small randomized trial utilizing a ketogenic diet demonstrated improved weight and diabetes control at one year (15). Systematic reviews also corroborate the effectiveness of a low-carbohydrate diet for T2D(16,S42) and it has recently become a consensus recommended dietary option(17). Nonetheless, sustained adherence is considered challenging(17), and an LDL-C increase is sometimes observed(18,S43,S44) with carbohydrate restriction. Given that total LDL-P, small LDL-P, and ApoB tend to improve or remain unchanged, the impact of an isolated increase in LDL-C on CVD risk in the context of this dietary pattern is unknown.

We have previously reported 1 year outcomes of an open-label, non-randomized, controlled, longitudinal study with 262 continuous care intervention (CCI) and 87 usual care (UC) participants with T2D(7). The CCI included telemedicine, health coaching, and guidance in nutritional ketosis using an individualized whole foods diet. Eighty-three percent of CCI participants remained enrolled 1 year and 60% of completers achieved an HbA1c <6.5% while prescribed metformin or no diabetes medication. Weight was reduced and most CVD risk factors improved(19). Here we report the results of this study at 2 years. The primary aims were to investigate the effect of the CCI on retention, glycemic control, and weight. Secondary aims included: (1) investigating the effect of the CCI on bone mineral density, visceral fat composition, cardiovascular risk factors, liver, kidney, thyroid and inflammatory markers; diabetes medication use, and disease outcomes (e.g. diabetes remission, metabolic syndrome); and (2) comparing 2-year outcomes between the CCI and UC groups.

## Materials and methods

### Study design and participants

The comprehensive study design has been published previously (7,25), and the results presented here are the follow-up 2-year results (*Clinical trials.gov identifier: NCT02519309*). This is an open-label, non-randomized, outpatient study and results presented here include data collected between August, 2015 and May, 2018. Participants aged 21 to 65 years with a confirmed diagnosis of T2D and a body mass index (BMI) > 25 kg/m^2^. Participants in the CCI accessed a remote care team consisting of a health coach and medical provider and reported routine biomarkers (weight, blood glucose and beta-hydroxybutyrate [BHB]) through a web-based application (app). Participants self-selected between two different CCI educational modes: on-site (n=136, CCI-onsite) or web-based (n=126, CCI-virtual). We also recruited another cohort of participants with T2D (n=87) who were categorized as usual care (UC). Exclusion criteria have been published previously (7,25). A brief description of the study participants and interventions (CCI and UC) are listed in the **supplementary data (Methods section)**. All study participants provided written informed consent and the study was approved by the Franciscan Health Lafayette Institutional Review Board.

### Outcomes

#### Primary Outcomes

The primary outcomes were retention, HbA1c, HOMA-IR-insulin and c-peptide derived (scores, equations in supplemental material A), fasting glucose, fasting insulin, c-peptide and weight.

#### Secondary Outcomes

Long-term body composition changes assessed in CCI participants included bone mineral density (BMD), abdominal fat content (CAF and A/G ratio), and lower extremities lean mss (LELM). Body composition was not assessed in UC participants. Cardiovascular-, liver-, kidney-, thyroid-related and inflammatory markers were analyzed (Table 1 and Supplementary Table 1). Changes in overall diabetes medication use, use by class, and insulin dose were tracked over the two years of the trial.

**Table 1.**
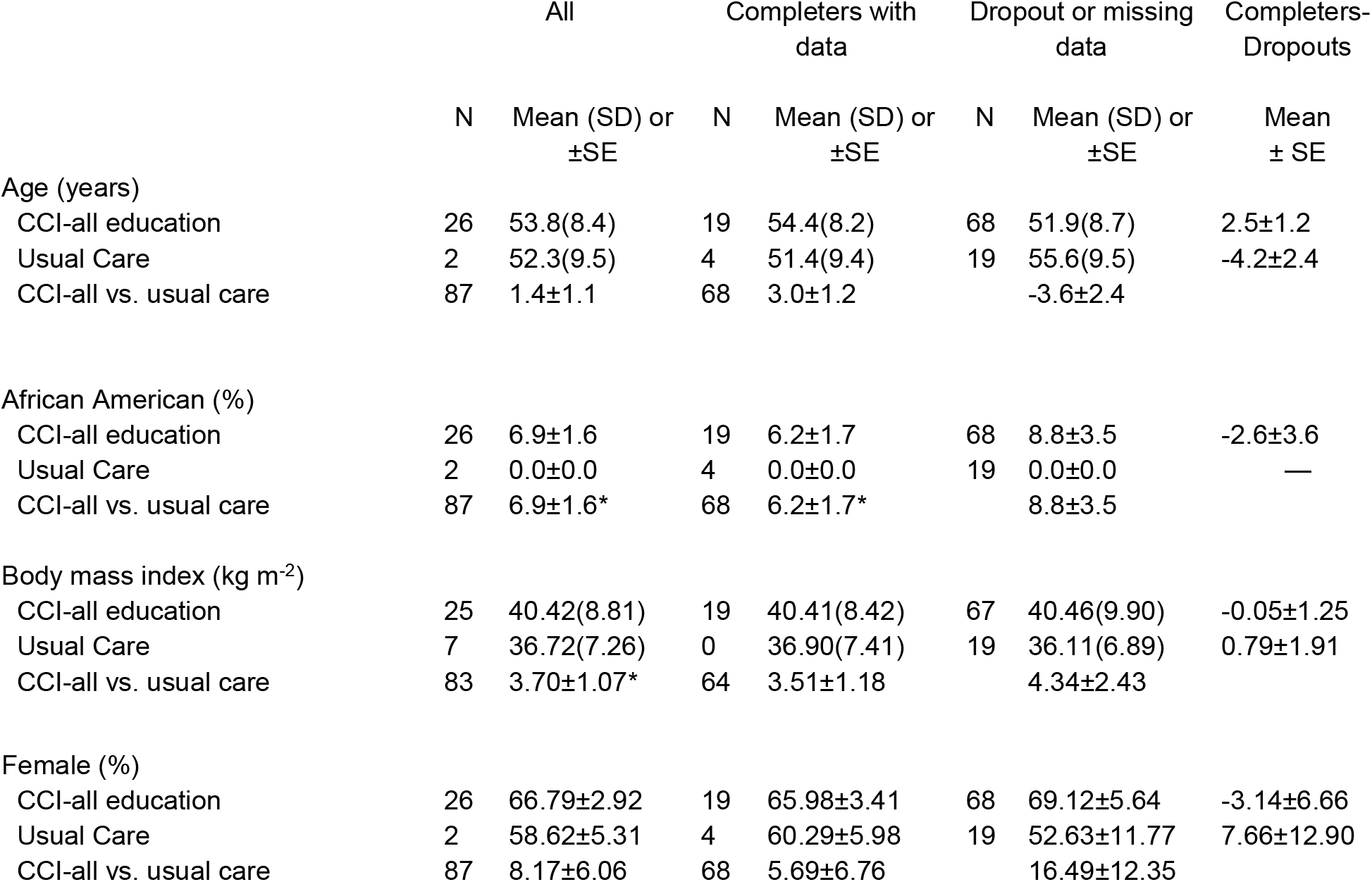
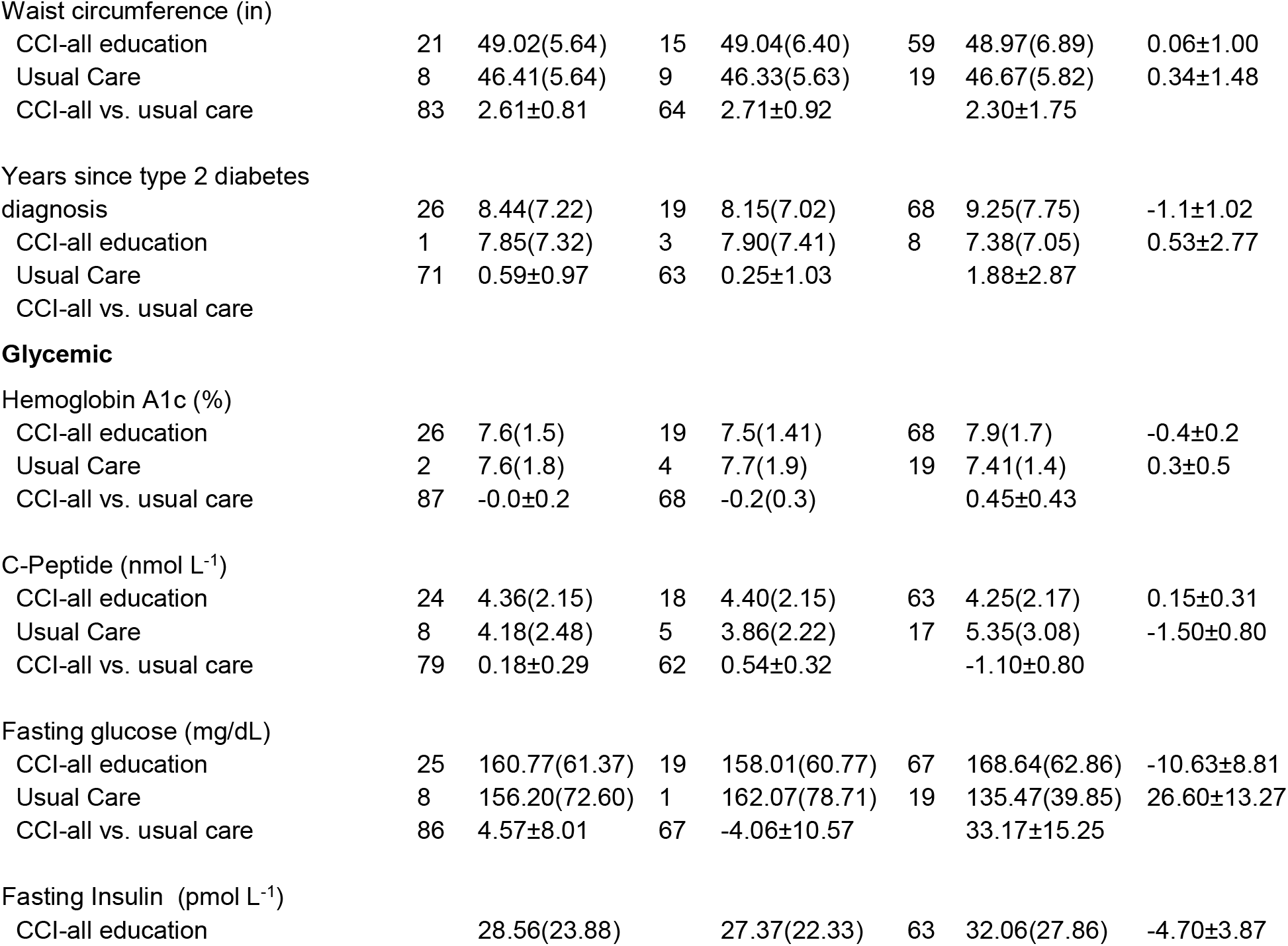
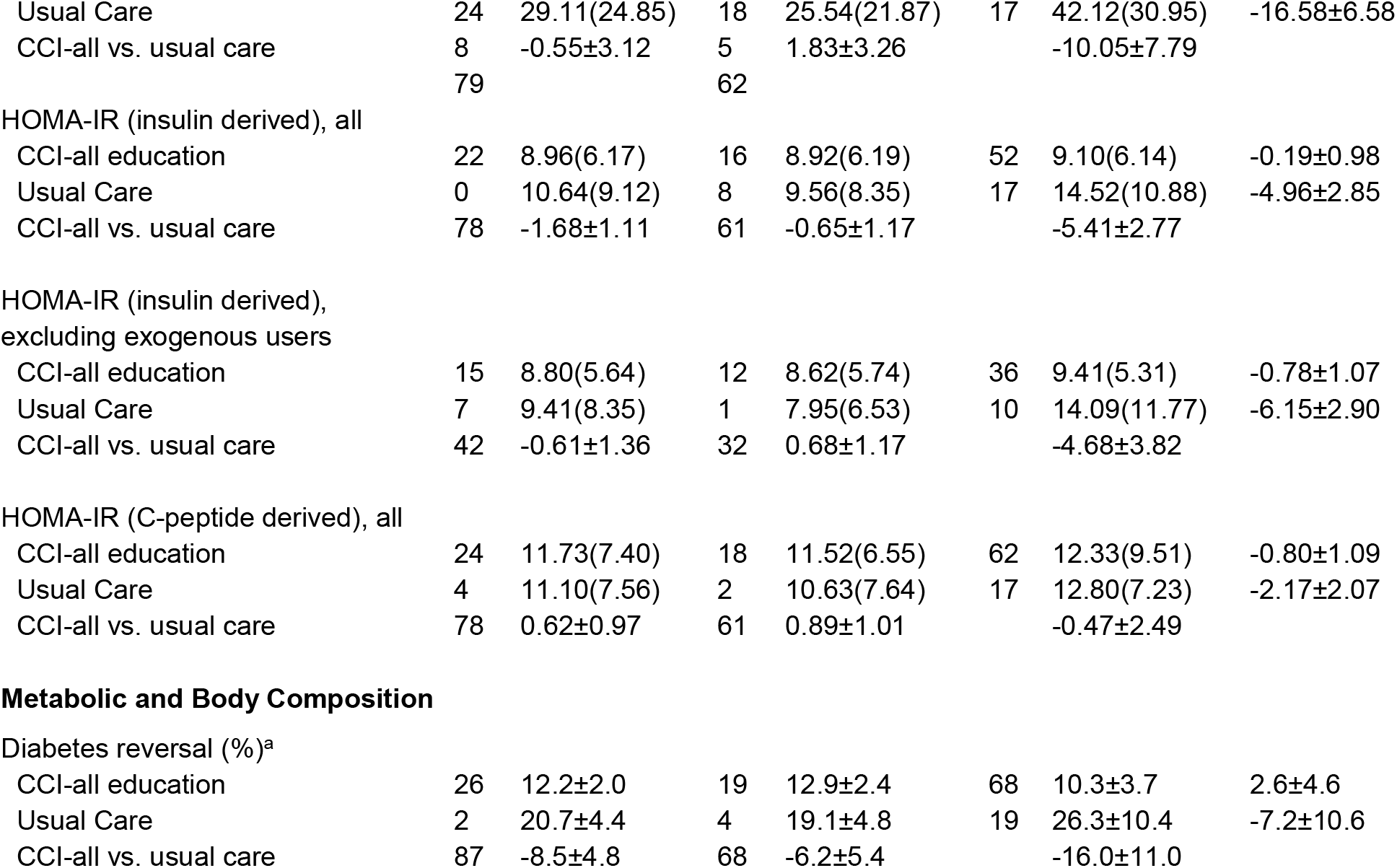
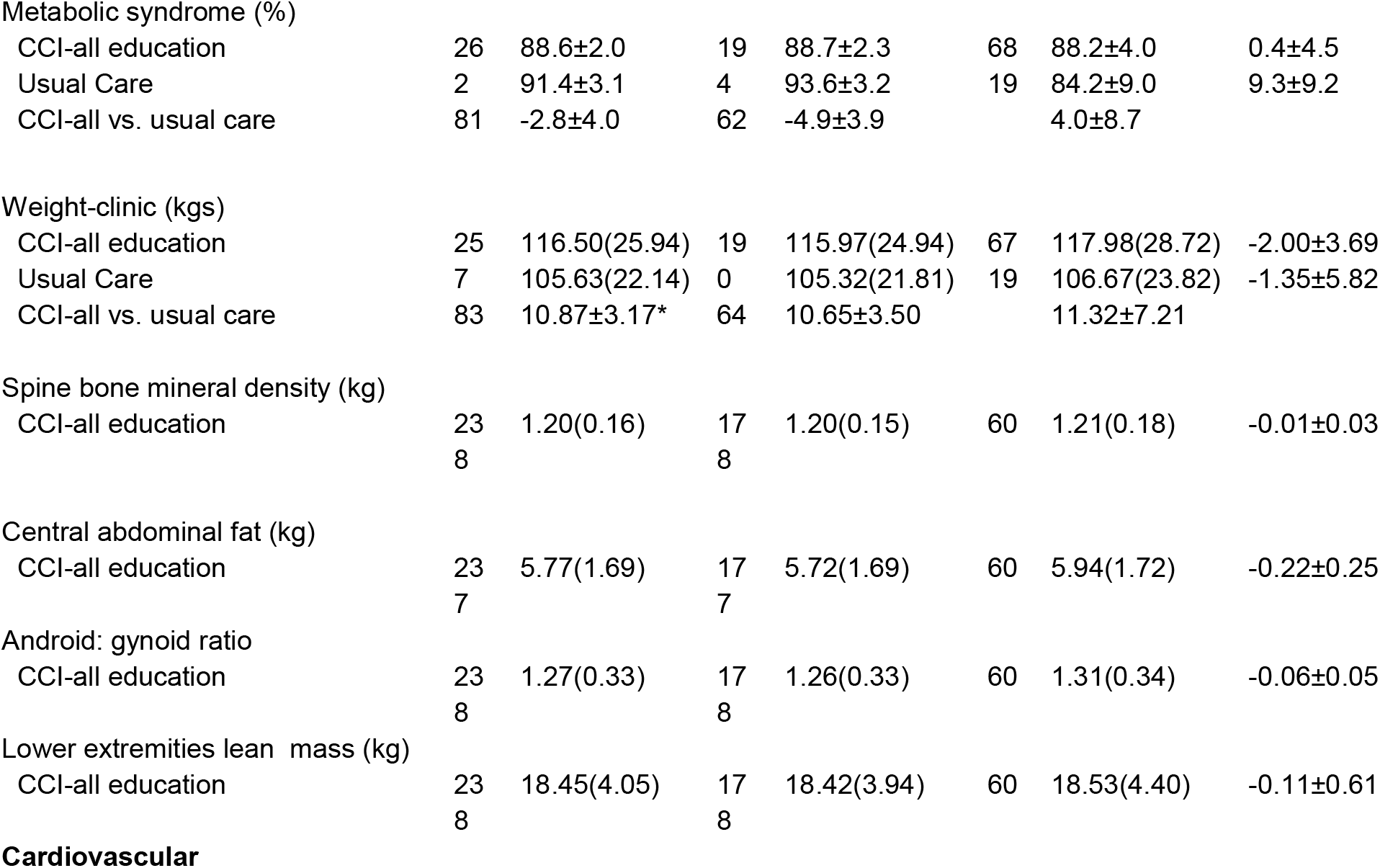
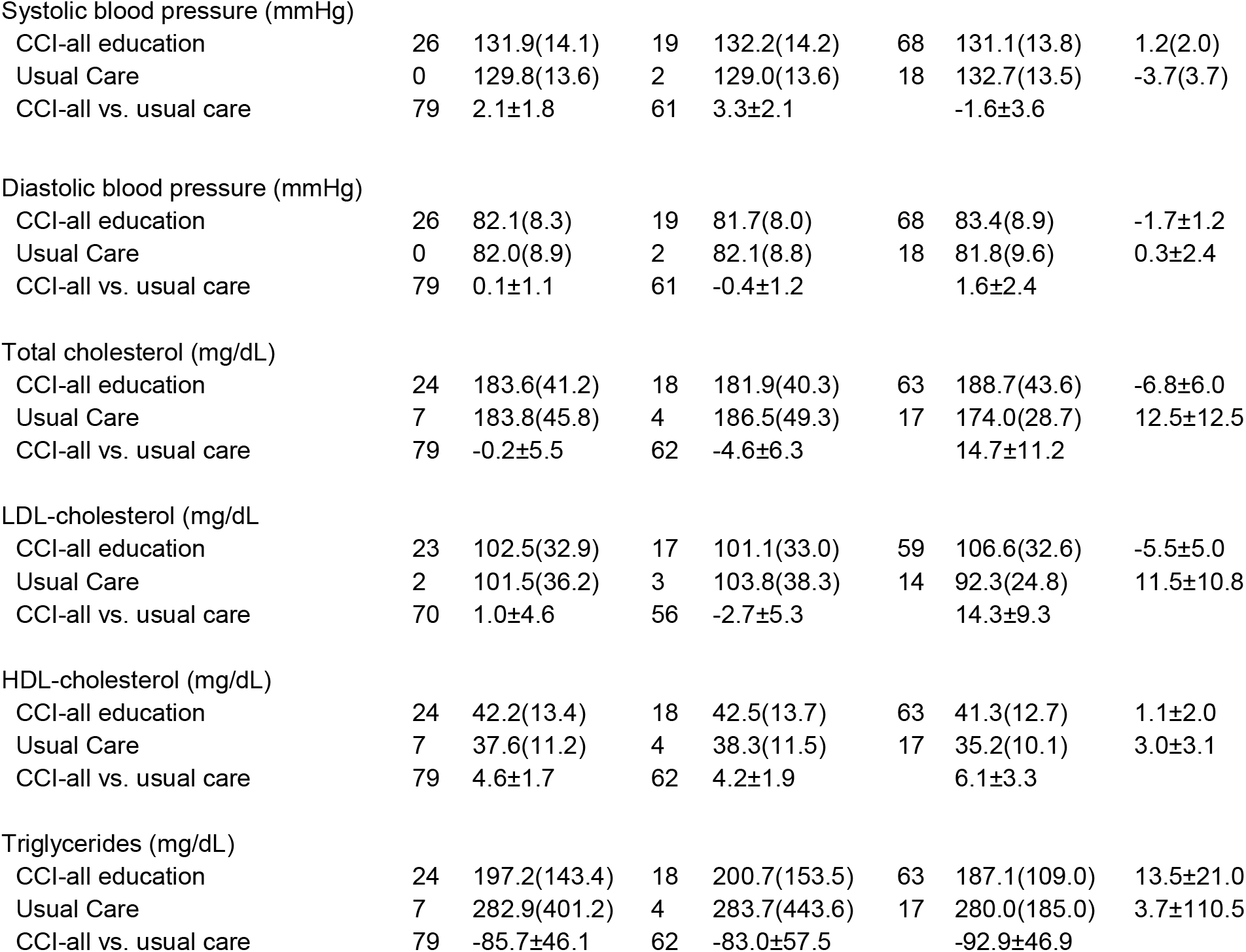
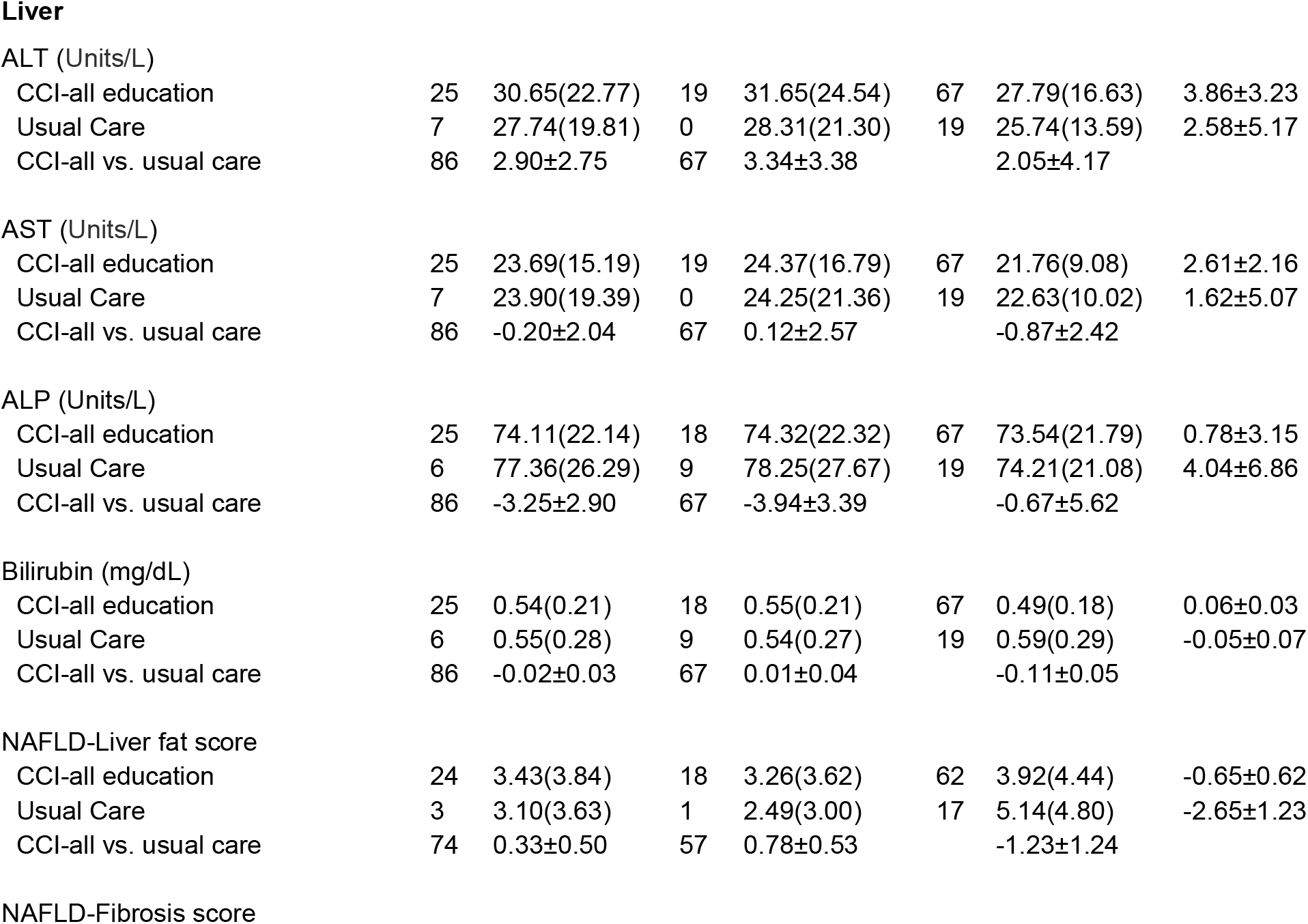
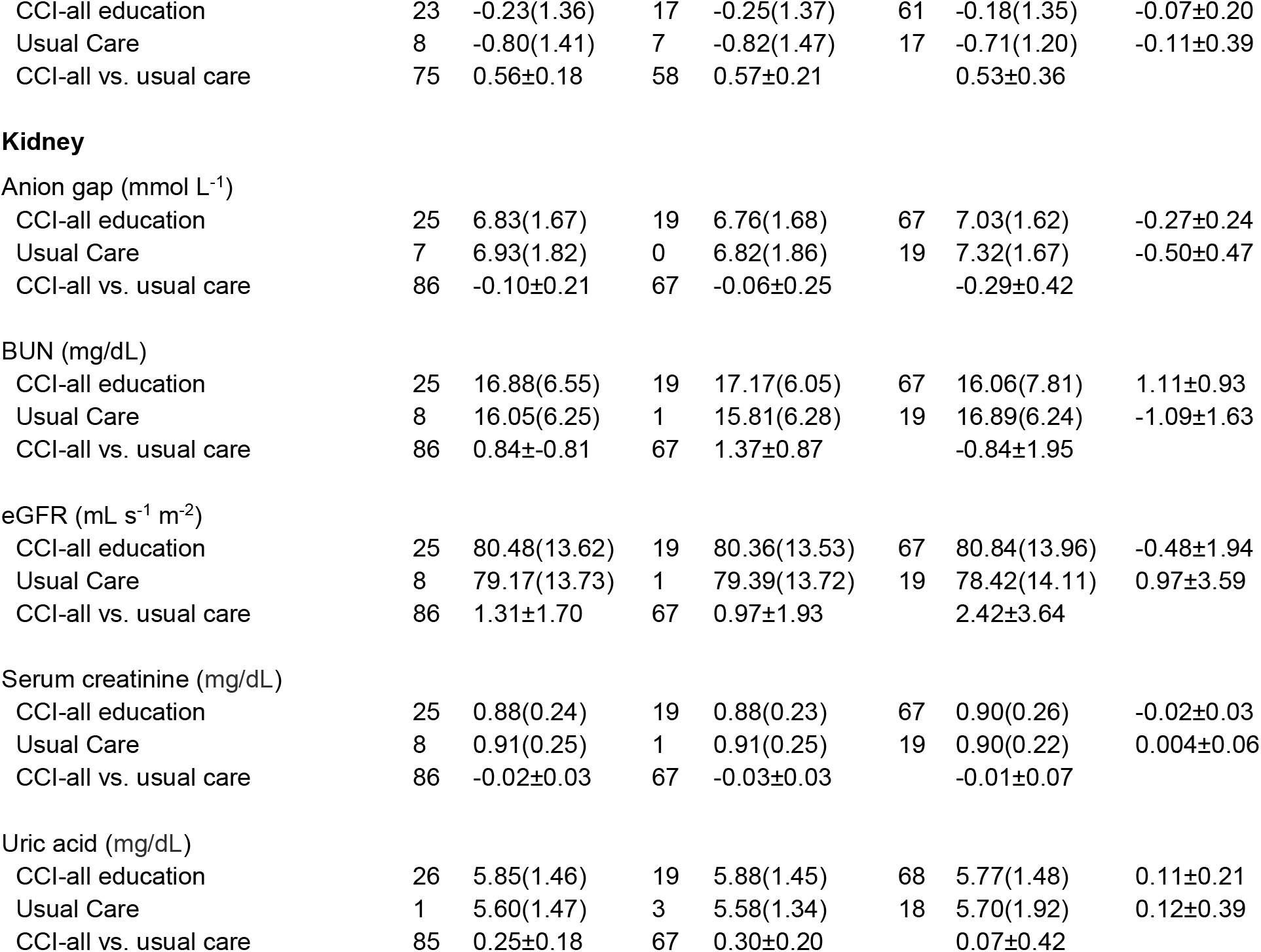
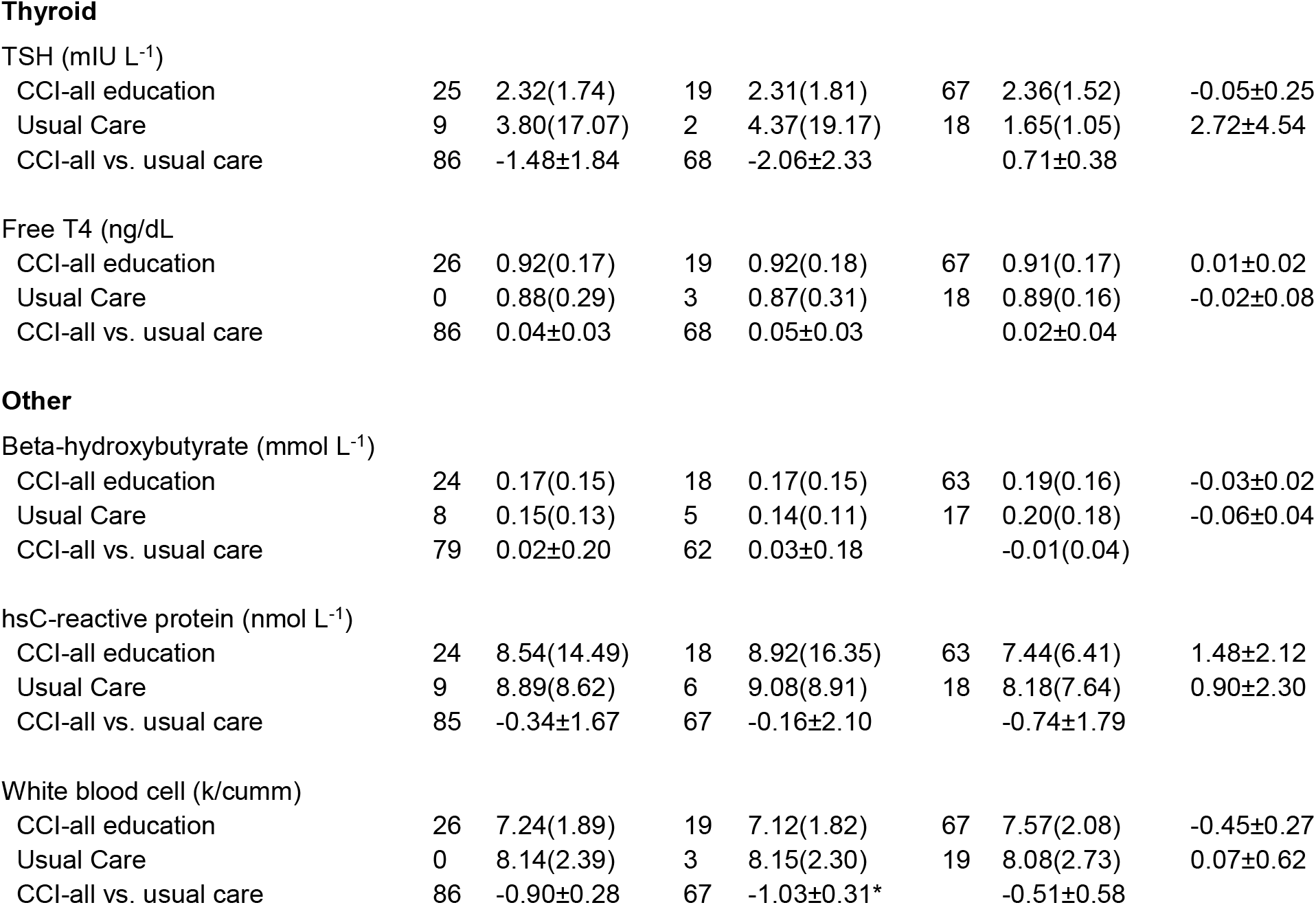
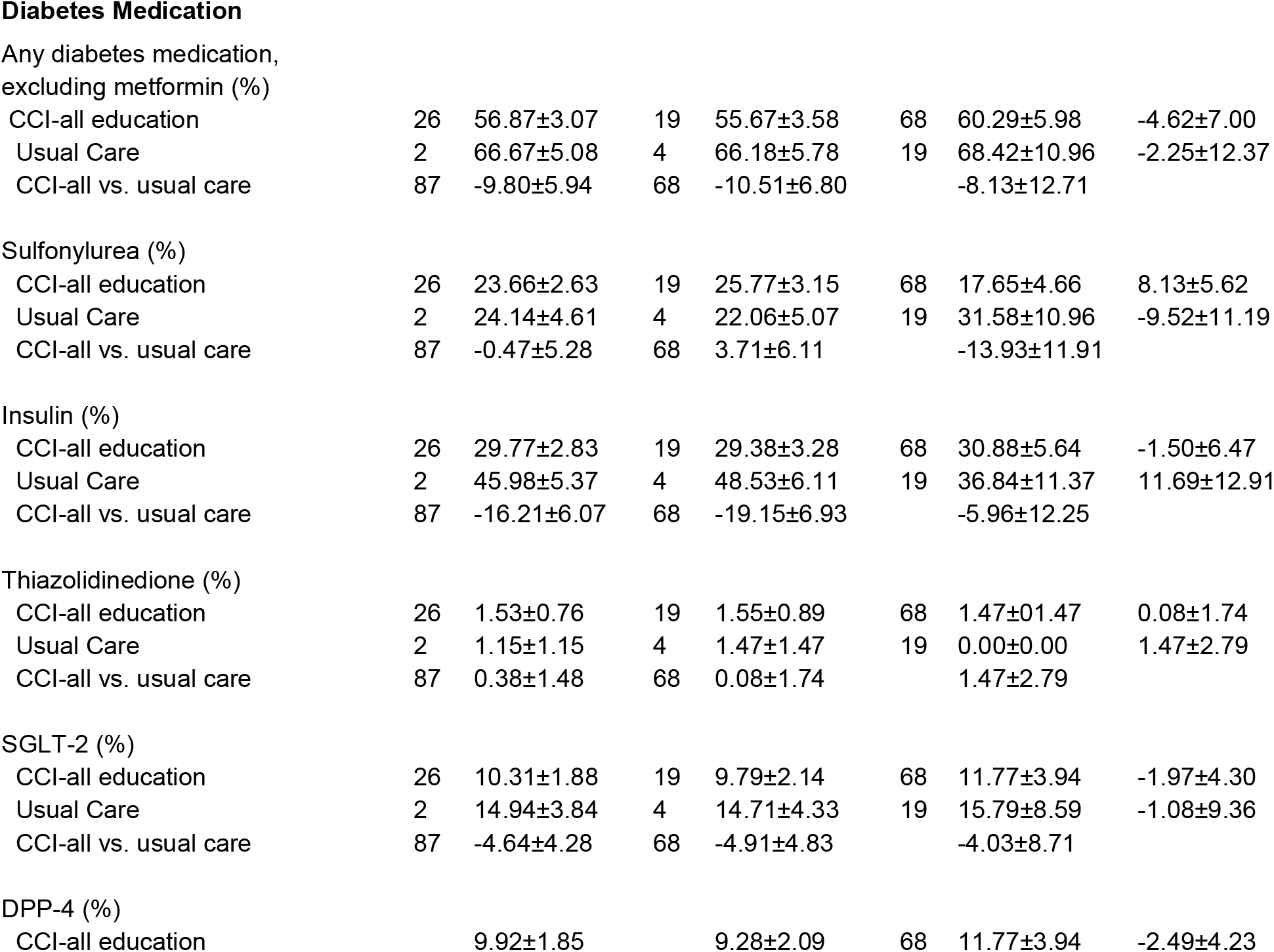
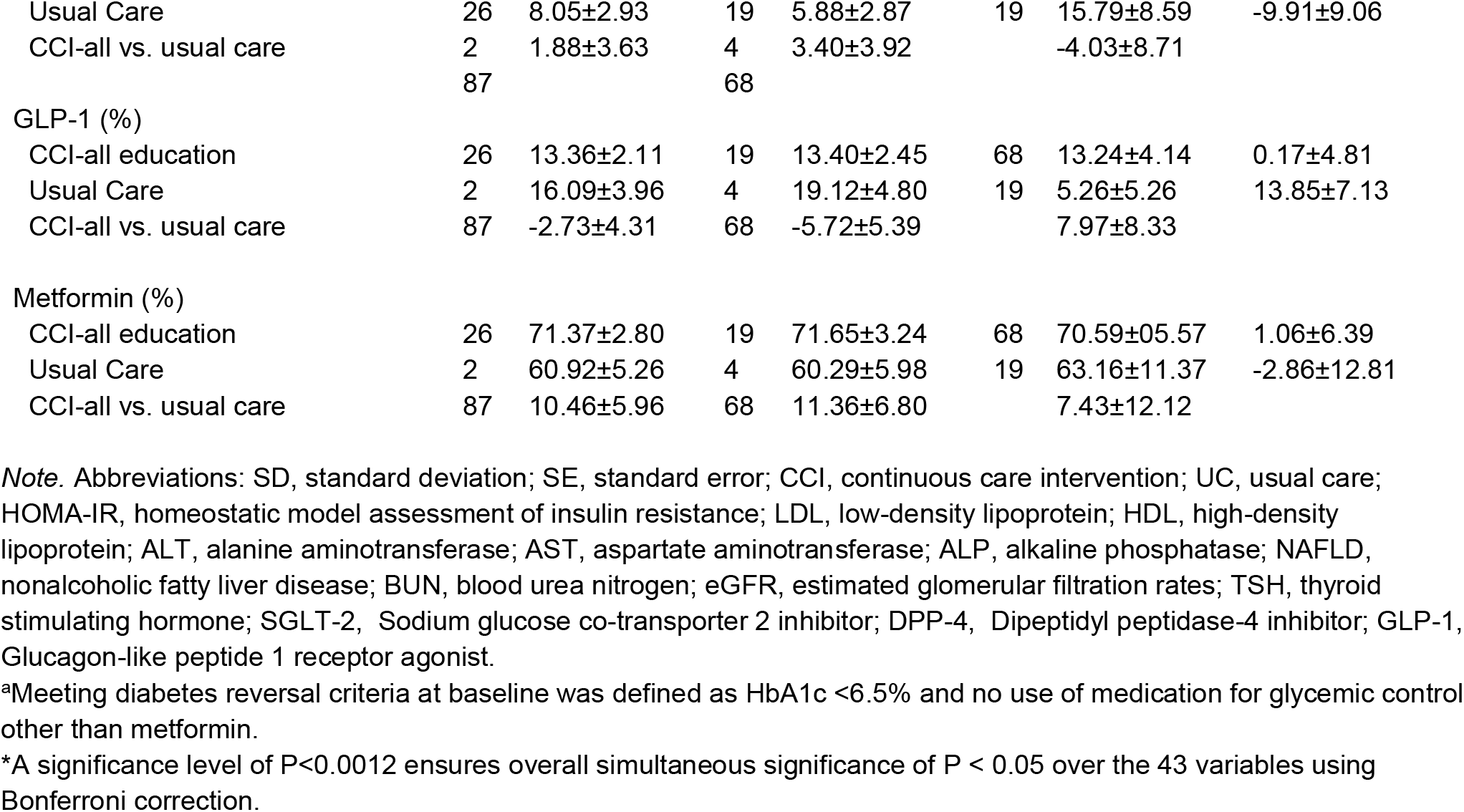
Baseline characteristics

|  | All |  | Completers with data |  | Dropout or missing data |  | Completers-Dropouts |
| --- | --- | --- | --- | --- | --- | --- | --- |
| | N | Mean (SD) or ±SE | N | Mean (SD) or ±SE | N | Mean (SD) or ±SE | Mean ± SE |
| Age (years) |  |  |  |  |  |  |  |
| CCI-all education | 26 | 53.8(8.4) | 19 | 54.4(8.2) | 68 | 51.9(8.7) | 2.5±1.2 |
| Usual Care | 2 | 52.3(9.5) | 4 | 51.4(9.4) | 19 | 55.6(9.5) | -4.2±2.4 |
| CCI-all vs. usual care | 87 | 1.4±1.1 | 68 | 3.0±1.2 | | -3.6±2.4 | |
| African American (%) |  |  |  |  |  |  |  |
| CCI-all education | 26 | 6.9±1.6 | 19 | 6.2±1.7 | 68 | 8.8±3.5 | -2.6±3.6 |
| Usual Care | 2 | 0.0±0.0 | 4 | 0.0±0.0 | 19 | 0.0±0.0 | — |
| CCI-all vs. usual care | 87 | 6.9±1.6* | 68 | 6.2±1.7* | | 8.8±3.5 | |
| Body mass index (kg m<sup>-2</sup>) |  |  |  |  |  |  |  |
| CCI-all education | 25 | 40.42(8.81) | 19 | 40.41(8.42) | 67 | 40.46(9.90) | -0.05±1.25 |
| Usual Care | 7 | 36.72(7.26) | 0 | 36.90(7.41) | 19 | 36.11(6.89) | 0.79±1.91 |
| CCI-all vs. usual care | 83 | 3.70±1.07* | 64 | 3.51±1.18 | | 4.34±2.43 | |
| Female (%) |  |  |  |  |  |  |  |
| CCI-all education | 26 | 66.79±2.92 | 19 | 65.98±3.41 | 68 | 69.12±5.64 | -3.14±6.66 |
| Usual Care | 2 | 58.62±5.31 | 4 | 60.29±5.98 | 19 | 52.63±11.77 | 7.66±12.90 |
| CCI-all vs. usual care | 87 | 8.17±6.06 | 68 | 5.69±6.76 | | 16.49±12.35 | |
| Waist circumference (in) |  |  |  |  |  |  |  |
| CCI-all education | 21 | 49.02(5.64) | 15 | 49.04(6.40) | 59 | 48.97(6.89) | 0.06±1.00 |
| Usual Care | 8 | 46.41(5.64) | 9 | 46.33(5.63) | 19 | 46.67(5.82) | 0.34±1.48 |
| CCI-all vs. usual care | 83 | 2.61±0.81 | 64 | 2.71±0.92 |  | 2.30±1.75 |  |
| Years since type 2 diabetes diagnosis |  |  |  |  |  |  |  |
| CCI-all education | 26 | 8.44(7.22) | 19 | 8.15(7.02) | 68 | 9.25(7.75) | -1.1±1.02 |
| Usual Care | 1 | 7.85(7.32) | 3 | 7.90(7.41) | 8 | 7.38(7.05) | 0.53±2.77 |
| CCI-all vs. usual care | 71 | 0.59±0.97 | 63 | 0.25±1.03 |  | 1.88±2.87 |  |
| **Glycemic** |  |  |  |  |  |  |  |
| Hemoglobin A1c (%) |  |  |  |  |  |  |  |
| CCI-all education | 26 | 7.6(1.5) | 19 | 7.5(1.41) | 68 | 7.9(1.7) | -0.4±0.2 |
| Usual Care | 2 | 7.6(1.8) | 4 | 7.7(1.9) | 19 | 7.41(1.4) | 0.3±0.5 |
| CCI-all vs. usual care | 87 | -0.0±0.2 | 68 | -0.2(0.3) |  | 0.45±0.43 |  |
| C-Peptide (nmol L<sup>-1</sup>) |  |  |  |  |  |  |  |
| CCI-all education | 24 | 4.36(2.15) | 18 | 4.40(2.15) | 63 | 4.25(2.17) | 0.15±0.31 |
| Usual Care | 8 | 4.18(2.48) | 5 | 3.86(2.22) | 17 | 5.35(3.08) | -1.50±0.80 |
| CCI-all vs. usual care | 79 | 0.18±0.29 | 62 | 0.54±0.32 |  | -1.10±0.80 |  |
| Fasting glucose (mg/dL) |  |  |  |  |  |  |  |
| CCI-all education | 25 | 160.77(61.37) | 19 | 158.01(60.77) | 67 | 168.64(62.86) | -10.63±8.81 |
| Usual Care | 8 | 156.20(72.60) | 1 | 162.07(78.71) | 19 | 135.47(39.85) | 26.60±13.27 |
| CCI-all vs. usual care | 86 | 4.57±8.01 | 67 | -4.06±10.57 |  | 33.17±15.25 |  |
| Fasting Insulin (pmol L<sup>-1</sup>) |  |  |  |  |  |  |  |
| CCI-all education |  | 28.56(23.88) |  | 27.37(22.33) | 63 | 32.06(27.86) | -4.70±3.87 |
| Usual Care | 24 | 29.11(24.85) | 18 | 25.54(21.87) | 17 | 42.12(30.95) | -16.58±6.58 |
| CCI-all vs. usual care | 8 | -0.55±3.12 | 5 | 1.83±3.26 |  | -10.05±7.79 |  |
|  | 79 |  | 62 |  |  |  |  |
| HOMA-IR (insulin derived), all |  |  |  |  |  |  |  |
| CCI-all education | 22 | 8.96(6.17) | 16 | 8.92(6.19) | 52 | 9.10(6.14) | -0.19±0.98 |
| Usual Care | 0 | 10.64(9.12) | 8 | 9.56(8.35) | 17 | 14.52(10.88) | -4.96±2.85 |
| CCI-all vs. usual care | 78 | -1.68±1.11 | 61 | -0.65±1.17 |  | -5.41±2.77 |  |
| HOMA-IR (insulin derived),<br>excluding exogenous users |  |  |  |  |  |  |  |
| CCI-all education | 15 | 8.80(5.64) | 12 | 8.62(5.74) | 36 | 9.41(5.31) | -0.78±1.07 |
| Usual Care | 7 | 9.41(8.35) | 1 | 7.95(6.53) | 10 | 14.09(11.77) | -6.15±2.90 |
| CCI-all vs. usual care | 42 | -0.61±1.36 | 32 | 0.68±1.17 |  | -4.68±3.82 |  |
| HOMA-IR (C-peptide derived), all |  |  |  |  |  |  |  |
| CCI-all education | 24 | 11.73(7.40) | 18 | 11.52(6.55) | 62 | 12.33(9.51) | -0.80±1.09 |
| Usual Care | 4 | 11.10(7.56) | 2 | 10.63(7.64) | 17 | 12.80(7.23) | -2.17±2.07 |
| CCI-all vs. usual care | 78 | 0.62±0.97 | 61 | 0.89±1.01 |  | -0.47±2.49 |  |
| Metabolic and Body Composition |  |  |  |  |  |  |  |
| Diabetes reversal (%)<sup>a</sup> |  |  |  |  |  |  |  |
| CCI-all education | 26 | 12.2±2.0 | 19 | 12.9±2.4 | 68 | 10.3±3.7 | 2.6±4.6 |
| Usual Care | 2 | 20.7±4.4 | 4 | 19.1±4.8 | 19 | 26.3±10.4 | -7.2±10.6 |
| CCI-all vs. usual care | 87 | -8.5±4.8 | 68 | -6.2±5.4 |  | -16.0±11.0 |  |
| Metabolic syndrome (%) |  |  |  |  |  |  |  |
| CCI-all education | 26 | 88.6±2.0 | 19 | 88.7±2.3 | 68 | 88.2±4.0 | 0.4±4.5 |
| Usual Care | 2 | 91.4±3.1 | 4 | 93.6±3.2 | 19 | 84.2±9.0 | 9.3±9.2 |
| CCI-all vs. usual care | 81 | -2.8±4.0 | 62 | -4.9±3.9 |  | 4.0±8.7 |  |
| Weight-clinic (kgs) |  |  |  |  |  |  |  |
| CCI-all education | 25 | 116.50(25.94) | 19 | 115.97(24.94) | 67 | 117.98(28.72) | -2.00±3.69 |
| Usual Care | 7 | 105.63(22.14) | 0 | 105.32(21.81) | 19 | 106.67(23.82) | -1.35±5.82 |
| CCI-all vs. usual care | 83 | 10.87±3.17* | 64 | 10.65±3.50 |  | 11.32±7.21 |  |
| Spine bone mineral density (kg) |  |  |  |  |  |  |  |
| CCI-all education | 23<br>8 | 1.20(0.16) | 17<br>8 | 1.20(0.15) | 60 | 1.21(0.18) | -0.01±0.03 |
| Central abdominal fat (kg) |  |  |  |  |  |  |  |
| CCI-all education | 23<br>7 | 5.77(1.69) | 17<br>7 | 5.72(1.69) | 60 | 5.94(1.72) | -0.22±0.25 |
| Android: gynoid ratio |  |  |  |  |  |  |  |
| CCI-all education | 23<br>8 | 1.27(0.33) | 17<br>8 | 1.26(0.33) | 60 | 1.31(0.34) | -0.06±0.05 |
| Lower extremities lean mass (kg) |  |  |  |  |  |  |  |
| CCI-all education | 23<br>8 | 18.45(4.05) | 17<br>8 | 18.42(3.94) | 60 | 18.53(4.40) | -0.11±0.61 |
| **Cardiovascular** |  |  |  |  |  |  |  |
| Systolic blood pressure (mmHg) |  |  |  |  |  |  |  |
| CCI-all education | 26 | 131.9(14.1) | 19 | 132.2(14.2) | 68 | 131.1(13.8) | 1.2(2.0) |
| Usual Care | 0 | 129.8(13.6) | 2 | 129.0(13.6) | 18 | 132.7(13.5) | -3.7(3.7) |
| CCI-all vs. usual care | 79 | 2.1±1.8 | 61 | 3.3±2.1 |  | -1.6±3.6 |  |
| Diastolic blood pressure (mmHg) |  |  |  |  |  |  |  |
| CCI-all education | 26 | 82.1(8.3) | 19 | 81.7(8.0) | 68 | 83.4(8.9) | -1.7±1.2 |
| Usual Care | 0 | 82.0(8.9) | 2 | 82.1(8.8) | 18 | 81.8(9.6) | 0.3±2.4 |
| CCI-all vs. usual care | 79 | 0.1±1.1 | 61 | -0.4±1.2 |  | 1.6±2.4 |  |
| Total cholesterol (mg/dL) |  |  |  |  |  |  |  |
| CCI-all education | 24 | 183.6(41.2) | 18 | 181.9(40.3) | 63 | 188.7(43.6) | -6.8±6.0 |
| Usual Care | 7 | 183.8(45.8) | 4 | 186.5(49.3) | 17 | 174.0(28.7) | 12.5±12.5 |
| CCI-all vs. usual care | 79 | -0.2±5.5 | 62 | -4.6±6.3 |  | 14.7±11.2 |  |
| LDL-cholesterol (mg/dL) |  |  |  |  |  |  |  |
| CCI-all education | 23 | 102.5(32.9) | 17 | 101.1(33.0) | 59 | 106.6(32.6) | -5.5±5.0 |
| Usual Care | 2 | 101.5(36.2) | 3 | 103.8(38.3) | 14 | 92.3(24.8) | 11.5±10.8 |
| CCI-all vs. usual care | 70 | 1.0±4.6 | 56 | -2.7±5.3 |  | 14.3±9.3 |  |
| HDL-cholesterol (mg/dL) |  |  |  |  |  |  |  |
| CCI-all education | 24 | 42.2(13.4) | 18 | 42.5(13.7) | 63 | 41.3(12.7) | 1.1±2.0 |
| Usual Care | 7 | 37.6(11.2) | 4 | 38.3(11.5) | 17 | 35.2(10.1) | 3.0±3.1 |
| CCI-all vs. usual care | 79 | 4.6±1.7 | 62 | 4.2±1.9 |  | 6.1±3.3 |  |
| Triglycerides (mg/dL) |  |  |  |  |  |  |  |
| CCI-all education | 24 | 197.2(143.4) | 18 | 200.7(153.5) | 63 | 187.1(109.0) | 13.5±21.0 |
| Usual Care | 7 | 282.9(401.2) | 4 | 283.7(443.6) | 17 | 280.0(185.0) | 3.7±110.5 |
| CCI-all vs. usual care | 79 | -85.7±46.1 | 62 | -83.0±57.5 |  | -92.9±46.9 |  |
| ALT (Units/L) |  |  |  |  |  |  |  |
| CCI-all education | 25 | 30.65(22.77) | 19 | 31.65(24.54) | 67 | 27.79(16.63) | 3.86±3.23 |
| Usual Care | 7 | 27.74(19.81) | 0 | 28.31(21.30) | 19 | 25.74(13.59) | 2.58±5.17 |
| CCI-all vs. usual care | 86 | 2.90±2.75 | 67 | 3.34±3.38 |  | 2.05±4.17 |  |
| AST (Units/L) |  |  |  |  |  |  |  |
| CCI-all education | 25 | 23.69(15.19) | 19 | 24.37(16.79) | 67 | 21.76(9.08) | 2.61±2.16 |
| Usual Care | 7 | 23.90(19.39) | 0 | 24.25(21.36) | 19 | 22.63(10.02) | 1.62±5.07 |
| CCI-all vs. usual care | 86 | -0.20±2.04 | 67 | 0.12±2.57 |  | -0.87±2.42 |  |
| ALP (Units/L) |  |  |  |  |  |  |  |
| CCI-all education | 25 | 74.11(22.14) | 18 | 74.32(22.32) | 67 | 73.54(21.79) | 0.78±3.15 |
| Usual Care | 6 | 77.36(26.29) | 9 | 78.25(27.67) | 19 | 74.21(21.08) | 4.04±6.86 |
| CCI-all vs. usual care | 86 | -3.25±2.90 | 67 | -3.94±3.39 |  | -0.67±5.62 |  |
| Bilirubin (mg/dL) |  |  |  |  |  |  |  |
| CCI-all education | 25 | 0.54(0.21) | 18 | 0.55(0.21) | 67 | 0.49(0.18) | 0.06±0.03 |
| Usual Care | 6 | 0.55(0.28) | 9 | 0.54(0.27) | 19 | 0.59(0.29) | -0.05±0.07 |
| CCI-all vs. usual care | 86 | -0.02±0.03 | 67 | 0.01±0.04 |  | -0.11±0.05 |  |
| NAFLD-Liver fat score |  |  |  |  |  |  |  |
| CCI-all education | 24 | 3.43(3.84) | 18 | 3.26(3.62) | 62 | 3.92(4.44) | -0.65±0.62 |
| Usual Care | 3 | 3.10(3.63) | 1 | 2.49(3.00) | 17 | 5.14(4.80) | -2.65±1.23 |
| CCI-all vs. usual care | 74 | 0.33±0.50 | 57 | 0.78±0.53 |  | -1.23±1.24 |  |
| NAFLD-Fibrosis score |  |  |  |  |  |  |  |
| CCI-all education | 23 | -0.23(1.36) | 17 | -0.25(1.37) | 61 | -0.18(1.35) | -0.07±0.20 |
| Usual Care | 8 | -0.80(1.41) | 7 | -0.82(1.47) | 17 | -0.71(1.20) | -0.11±0.39 |
| CCI-all vs. usual care | 75 | 0.56±0.18 | 58 | 0.57±0.21 |  | 0.53±0.36 |  |
| CCI-all education | 25 | 6.83(1.67) | 19 | 6.76(1.68) | 67 | 7.03(1.62) | -0.27±0.24 |
| Usual Care | 7 | 6.93(1.82) | 0 | 6.82(1.86) | 19 | 7.32(1.67) | -0.50±0.47 |
| CCI-all vs. usual care | 86 | -0.10±0.21 | 67 | -0.06±0.25 |  | -0.29±0.42 |  |
| BUN (mg/dL) |  |  |  |  |  |  |  |
| CCI-all education | 25 | 16.88(6.55) | 19 | 17.17(6.05) | 67 | 16.06(7.81) | 1.11±0.93 |
| Usual Care | 8 | 16.05(6.25) | 1 | 15.81(6.28) | 19 | 16.89(6.24) | -1.09±1.63 |
| CCI-all vs. usual care | 86 | 0.84±0.81 | 67 | 1.37±0.87 |  | -0.84±1.95 |  |
| eGFR (mL s<sup>-1</sup> m<sup>-2</sup>) |  |  |  |  |  |  |  |
| CCI-all education | 25 | 80.48(13.62) | 19 | 80.36(13.53) | 67 | 80.84(13.96) | -0.48±1.94 |
| Usual Care | 8 | 79.17(13.73) | 1 | 79.39(13.72) | 19 | 78.42(14.11) | 0.97±3.59 |
| CCI-all vs. usual care | 86 | 1.31±1.70 | 67 | 0.97±1.93 |  | 2.42±3.64 |  |
| Serum creatinine (mg/dL) |  |  |  |  |  |  |  |
| CCI-all education | 25 | 0.88(0.24) | 19 | 0.88(0.23) | 67 | 0.90(0.26) | -0.02±0.03 |
| Usual Care | 8 | 0.91(0.25) | 1 | 0.91(0.25) | 19 | 0.90(0.22) | 0.004±0.06 |
| CCI-all vs. usual care | 86 | -0.02±0.03 | 67 | -0.03±0.03 |  | -0.01±0.07 |  |
| Uric acid (mg/dL) |  |  |  |  |  |  |  |
| CCI-all education | 26 | 5.85(1.46) | 19 | 5.88(1.45) | 68 | 5.77(1.48) | 0.11±0.21 |
| Usual Care | 1 | 5.60(1.47) | 3 | 5.58(1.34) | 18 | 5.70(1.92) | 0.12±0.39 |
| CCI-all vs. usual care | 85 | 0.25±0.18 | 67 | 0.30±0.20 |  | 0.07±0.42 |  |
| TSH (mIU L<sup>-1</sup>) |  |  |  |  |  |  |  |
| CCI-all education | 25 | 2.32(1.74) | 19 | 2.31(1.81) | 67 | 2.36(1.52) | -0.05±0.25 |
| Usual Care | 9 | 3.80(17.07) | 2 | 4.37(19.17) | 18 | 1.65(1.05) | 2.72±4.54 |
| CCI-all vs. usual care | 86 | -1.48±1.84 | 68 | -2.06±2.33 |  | 0.71±0.38 |  |
| Free T4 (ng/dL) |  |  |  |  |  |  |  |
| CCI-all education | 26 | 0.92(0.17) | 19 | 0.92(0.18) | 67 | 0.91(0.17) | 0.01±0.02 |
| Usual Care | 0 | 0.88(0.29) | 3 | 0.87(0.31) | 18 | 0.89(0.16) | -0.02±0.08 |
| CCI-all vs. usual care | 86 | 0.04±0.03 | 68 | 0.05±0.03 |  | 0.02±0.04 |  |
| Beta-hydroxybutyrate (mmol L<sup>-1</sup>) |  |  |  |  |  |  |  |
| CCI-all education | 24 | 0.17(0.15) | 18 | 0.17(0.15) | 63 | 0.19(0.16) | -0.03±0.02 |
| Usual Care | 8 | 0.15(0.13) | 5 | 0.14(0.11) | 17 | 0.20(0.18) | -0.06±0.04 |
| CCI-all vs. usual care | 79 | 0.02±0.20 | 62 | 0.03±0.18 |  | -0.01(0.04) |  |
| hsC-reactive protein (nmol L<sup>-1</sup>) |  |  |  |  |  |  |  |
| CCI-all education | 24 | 8.54(14.49) | 18 | 8.92(16.35) | 63 | 7.44(6.41) | 1.48±2.12 |
| Usual Care | 9 | 8.89(8.62) | 6 | 9.08(8.91) | 18 | 8.18(7.64) | 0.90±2.30 |
| CCI-all vs. usual care | 85 | -0.34±1.67 | 67 | -0.16±2.10 |  | -0.74±1.79 |  |
| White blood cell (k/cumm) |  |  |  |  |  |  |  |
| CCI-all education | 26 | 7.24(1.89) | 19 | 7.12(1.82) | 67 | 7.57(2.08) | -0.45±0.27 |
| Usual Care | 0 | 8.14(2.39) | 3 | 8.15(2.30) | 19 | 8.08(2.73) | 0.07±0.62 |
| CCI-all vs. usual care | 86 | -0.90±0.28 | 67 | -1.03±0.31* |  | -0.51±0.58 |  |
| CCI-all education | 26 | 56.87±3.07 | 19 | 55.67±3.58 | 68 | 60.29±5.98 | -4.62±7.00 |
| Usual Care | 2 | 66.67±5.08 | 4 | 66.18±5.78 | 19 | 68.42±10.96 | -2.25±12.37 |
| CCI-all vs. usual care | 87 | -9.80±5.94 | 68 | -10.51±6.80 |  | -8.13±12.71 |  |
| Sulfonylurea (%) |  |  |  |  |  |  |  |
| CCI-all education | 26 | 23.66±2.63 | 19 | 25.77±3.15 | 68 | 17.65±4.66 | 8.13±5.62 |
| Usual Care | 2 | 24.14±4.61 | 4 | 22.06±5.07 | 19 | 31.58±10.96 | -9.52±11.19 |
| CCI-all vs. usual care | 87 | -0.47±5.28 | 68 | 3.71±6.11 |  | -13.93±11.91 |  |
| Insulin (%) |  |  |  |  |  |  |  |
| CCI-all education | 26 | 29.77±2.83 | 19 | 29.38±3.28 | 68 | 30.88±5.64 | -1.50±6.47 |
| Usual Care | 2 | 45.98±5.37 | 4 | 48.53±6.11 | 19 | 36.84±11.37 | 11.69±12.91 |
| CCI-all vs. usual care | 87 | -16.21±6.07 | 68 | -19.15±6.93 |  | -5.96±12.25 |  |
| Thiazolidinedione (%) |  |  |  |  |  |  |  |
| CCI-all education | 26 | 1.53±0.76 | 19 | 1.55±0.89 | 68 | 1.47±01.47 | 0.08±1.74 |
| Usual Care | 2 | 1.15±1.15 | 4 | 1.47±1.47 | 19 | 0.00±0.00 | 1.47±2.79 |
| CCI-all vs. usual care | 87 | 0.38±1.48 | 68 | 0.08±1.74 |  | 1.47±2.79 |  |
| SGLT-2 (%) |  |  |  |  |  |  |  |
| CCI-all education | 26 | 10.31±1.88 | 19 | 9.79±2.14 | 68 | 11.77±3.94 | -1.97±4.30 |
| Usual Care | 2 | 14.94±3.84 | 4 | 14.71±4.33 | 19 | 15.79±8.59 | -1.08±9.36 |
| CCI-all vs. usual care | 87 | -4.64±4.28 | 68 | -4.91±4.83 |  | -4.03±8.71 |  |
| DPP-4 (%) |  |  |  |  |  |  |  |
| CCI-all education |  | 9.92±1.85 |  | 9.28±2.09 | 68 | 11.77±3.94 | -2.49±4.23 |
| Usual Care | 26 | 8.05±2.93 | 19 | 5.88±2.87 | 19 | 15.79±8.59 | -9.91±9.06 |
| CCI-all vs. usual care | 2 | 1.88±3.63 | 4 | 3.40±3.92 |  | -4.03±8.71 |  |
|  | 87 |  | 68 |  |  |  |  |
| GLP-1 (%) |  |  |  |  |  |  |  |
| CCI-all education | 26 | 13.36±2.11 | 19 | 13.40±2.45 | 68 | 13.24±4.14 | 0.17±4.81 |
| Usual Care | 2 | 16.09±3.96 | 4 | 19.12±4.80 | 19 | 5.26±5.26 | 13.85±7.13 |
| CCI-all vs. usual care | 87 | -2.73±4.31 | 68 | -5.72±5.39 |  | 7.97±8.33 |  |
| Metformin (%) |  |  |  |  |  |  |  |
| CCI-all education | 26 | 71.37±2.80 | 19 | 71.65±3.24 | 68 | 70.59±05.57 | 1.06±6.39 |
| Usual Care | 2 | 60.92±5.26 | 4 | 60.29±5.98 | 19 | 63.16±11.37 | -2.86±12.81 |
| CCI-all vs. usual care | 87 | 10.46±5.96 | 68 | 11.36±6.80 |  | 7.43±12.12 |  |

*Note.* Abbreviations: SD, standard deviation; SE, standard error; CCI, continuous care intervention; UC, usual care; HOMA-IR, homeostatic model assessment of insulin resistance; LDL, low-density lipoprotein; HDL, high-density lipoprotein; ALT, alanine aminotransferase; AST, aspartate aminotransferase; ALP, alkaline phosphatase; NAFLD, nonalcoholic fatty liver disease; BUN, blood urea nitrogen; eGFR, estimated glomerular filtration rates; TSH, thyroid stimulating hormone; SGLT-2, Sodium glucose co-transporter 2 inhibitor; DPP-4, Dipeptidyl peptidase-4 inhibitor; GLP-1, Glucagon-like peptide 1 receptor agonist.
<sup>a</sup>Meeting diabetes reversal criteria at baseline was defined as HbA1c <6.5% and no use of medication for glycemic control other than metformin.
\*A significance level of P<0.0012 ensures overall simultaneous significance of P < 0.05 over the 43 variables using Bonferroni correction.

The prevalence of T2D (diabetes reversal, partial and complete remission), metabolic syndrome, suspected steatosis and absence of fibrosis were evaluated at 2 years in the CCI and UC groups using the criteria provided in Supplementary Table 2 (assignment references listed in the supplementary). Assignment of metabolic syndrome was based on the presence of three of the five defined criteria according to measured laboratory and anthropometric variables; pharmacological treatment for any of the conditions was not considered.

Adverse events encountered in the study were reported to the Principal Investigator and reviewed by the Institutional Review Board (IRB).

### Laboratory and body composition measures

Clinical anthropometrics and laboratory blood analytes measurements were obtained at baseline, 1 year, and 2 years from the CCI and UC participants. Details of the methods were previously published(7,19). All blood analytes were measured at a Clinical Laboratory Improvement Amendments (CLIA) certified laboratory. The CCI participants were also assessed for total body composition changes at baseline, 1 and 2 years using dual X-ray absorptiometry (DXA) (Lunar GE Prodigy, Madison, WI) and analyzed using GE Encore software(v11.10, Madison, WI). The details of the DXA procedure and analyses are listed in the **supplementary data (Methods section)**.

### Statistical analyses

All analyses were conducted using SPSS statistical software (Version 25.0, Armonk, NY). A detailed description of the statistical method is included in the **supplementary data (Methods section)**. Briefly, we conducted intent-to-treat analyses to assess study outcomes. For continuous study outcomes, linear mixed-effects (LMM) models were used to assess within-group changes from baseline to 2 years and between-group differences at 2 years. For dichotomous disease outcomes, generalized estimating equation models were used. Changes in diabetes medication use and insulin dosage from baseline to 2 years were assessed using McNemar’s tests with continuity correction when appropriate and paired t-tests. Available data only was used to assess changes in medication use, which was routinely adjusted as part of the intervention protocol. Data from the two CCI educational groups were combined because no group differences were found, as in our prior time points(7,S45). Completers-only analyses were also conducted for all outcomes and results appear in the supplementary material. For all study analyses, nominal significance levels (P) are presented in the tables. A significance level of *P*<0.0012 ensures overall simultaneous significance of *P*<0.05 over the 43 variables using Bonferroni correction.

## Results

### Participant characteristics

Table 1 presents baseline characteristics of the 262 CCI and 87 UC participants. Participants did not differ between groups in demographic characteristics, except the proportion of African Americans was higher in the CCI group. Baseline characteristics were well-matched between the groups, except for mean weight and BMI, which were higher in the CCI group. There were no significant differences between completers and dropouts on baseline characteristics for either group.

### Retention and long-term dietary adherence

One hundred ninety four participants (of 262; 74%) remained enrolled in the CCI at 2 years (Figure 1), as did 78% of the UC group participants (68 of 87). CCI participant-reported reasons for dropout included: intervening life events (e.g. family emergencies), difficulty attending or completing laboratory and clinic visits associated with the trial, and insufficient motivation for participation in the intervention. At both 1 and 2 years, laboratory measured blood BHB was 0.3 ± 0.0 mmol L^-1^, about 1.5 fold higher than the baseline value (0.2 ± 0.0 mmol L^-1^). The mean laboratory BHB level was stable from 1 to 2 years, and 61.5% (n=161) of participants reported a blood BHB measurement ≥0.5mmol L^-1^ in the app at least once between 1 and 2 years.

**Figure 1.**
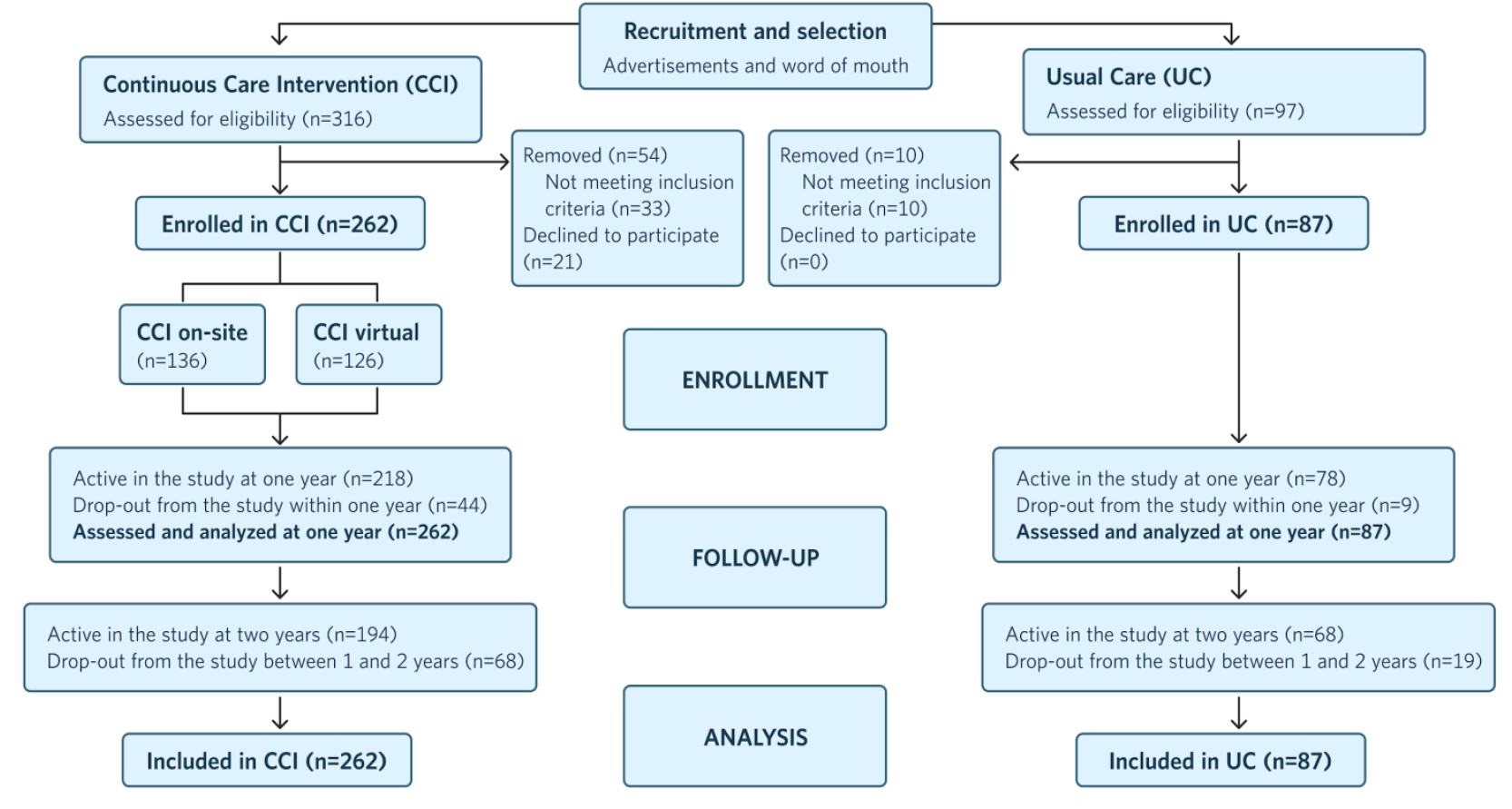
Flow chart of participants in each stage of the study from recruitment to 2 years post-enrollment and analysis.

All adjusted within and between group changes in study outcomes for the CCI and UC groups appear in Table 2.

**Table 2.**
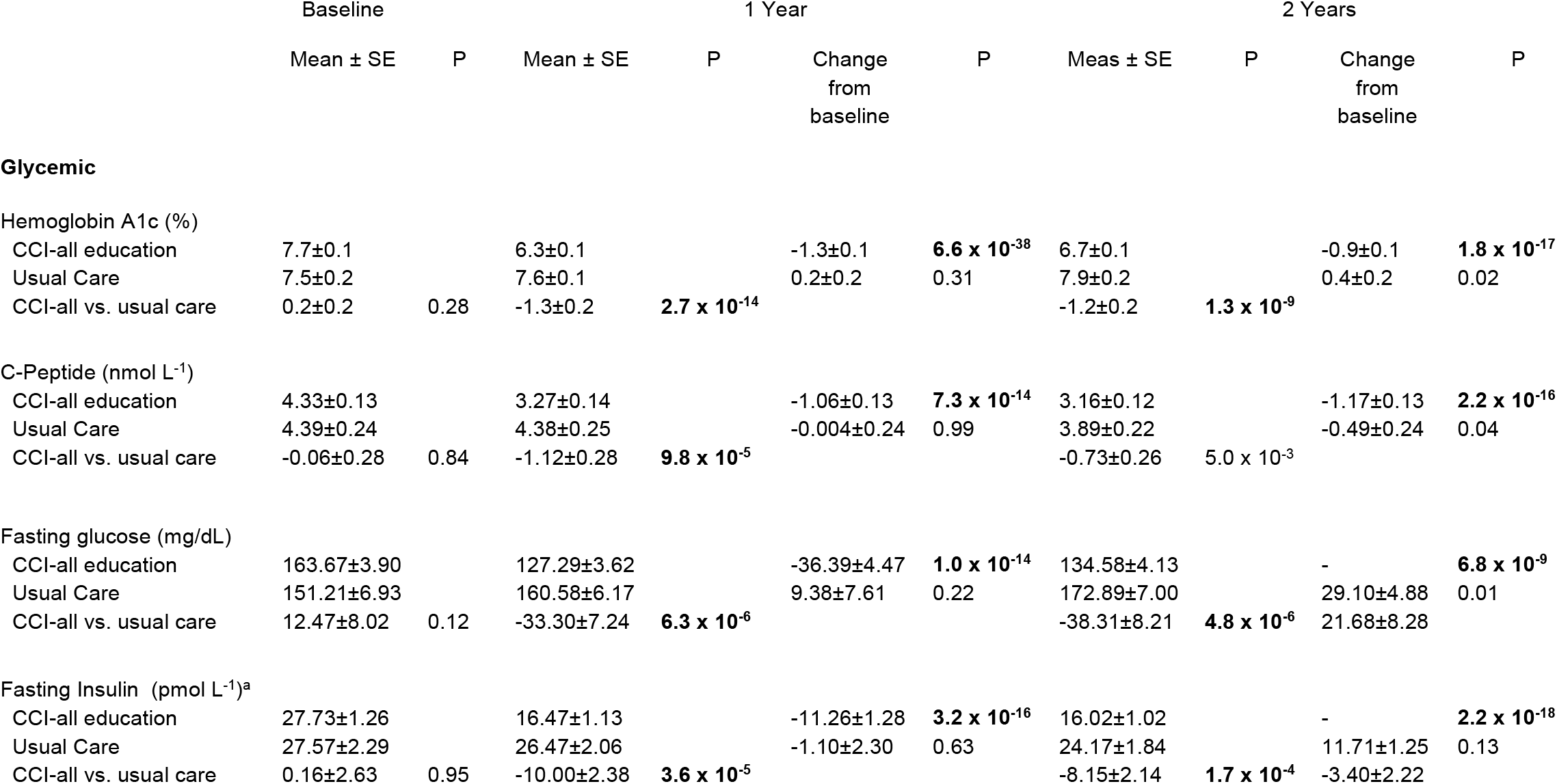
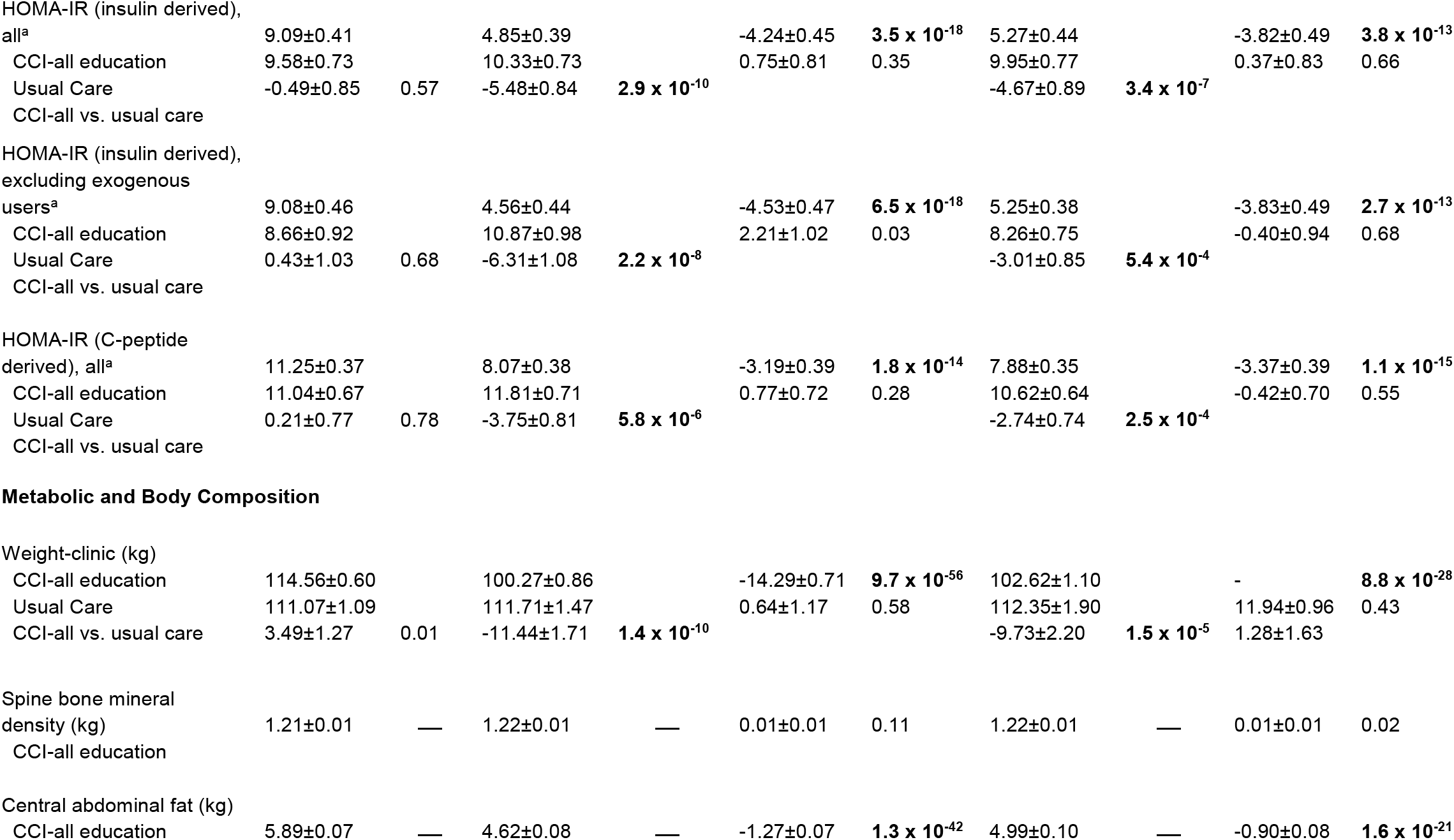
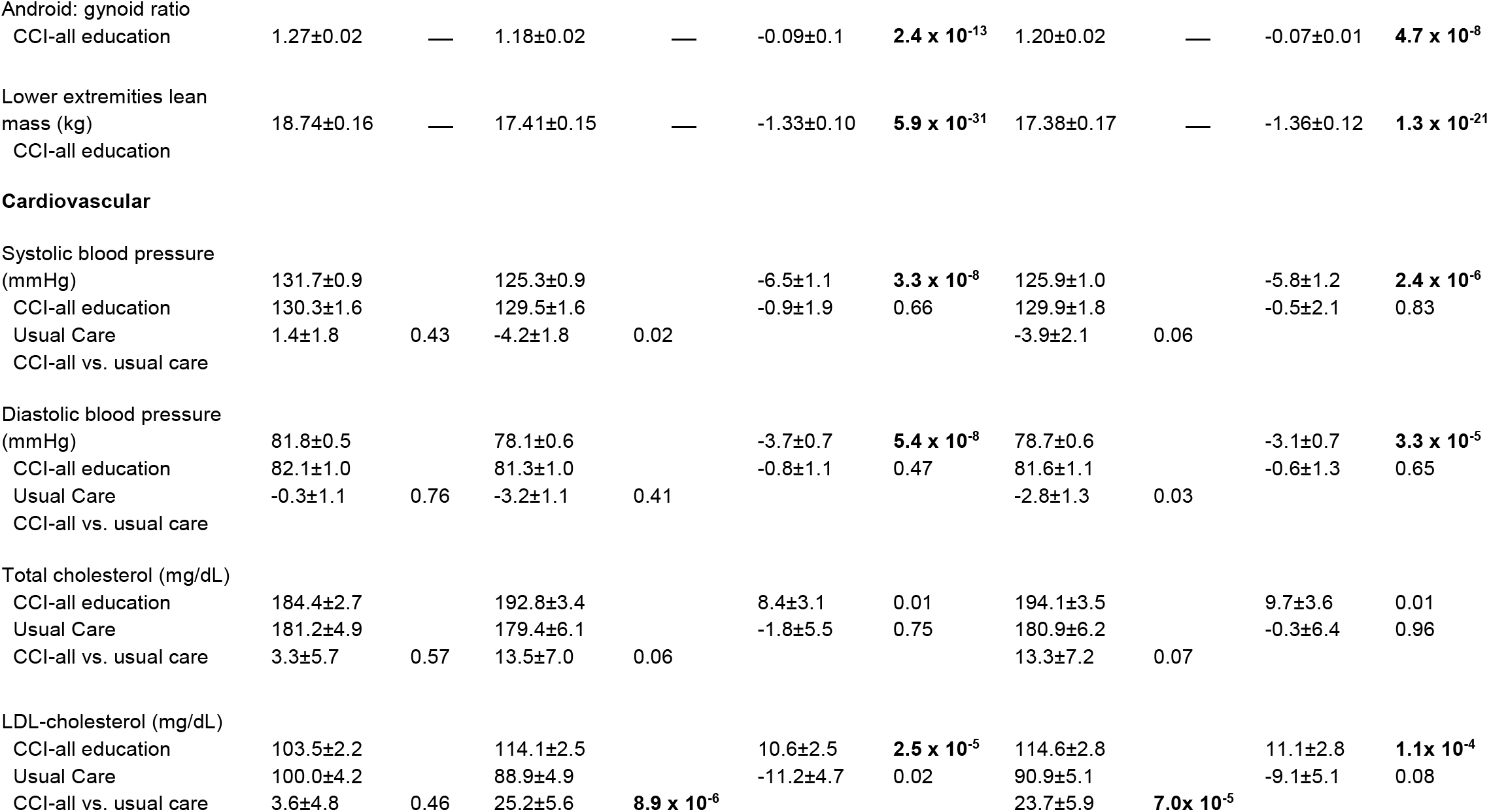
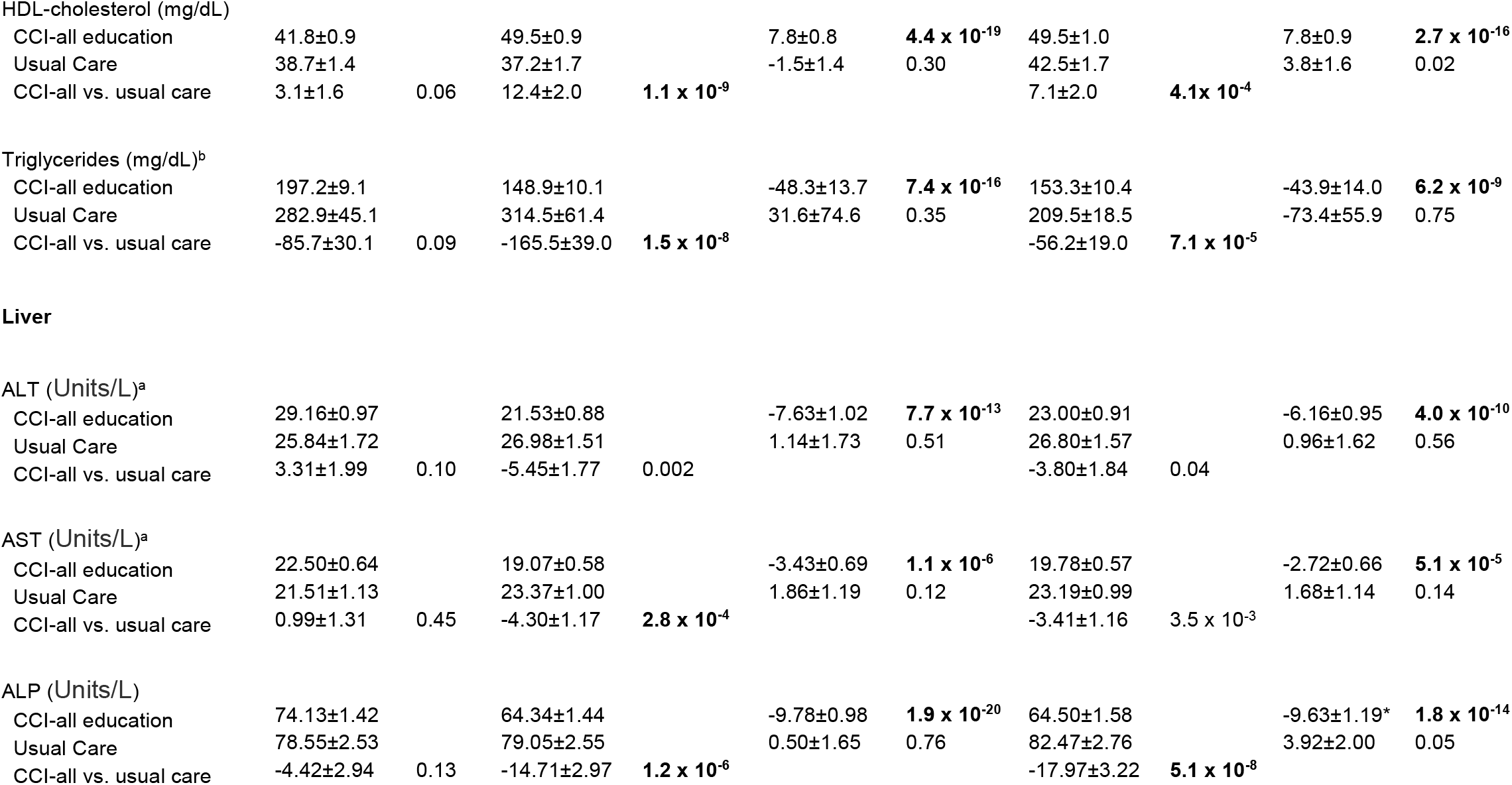
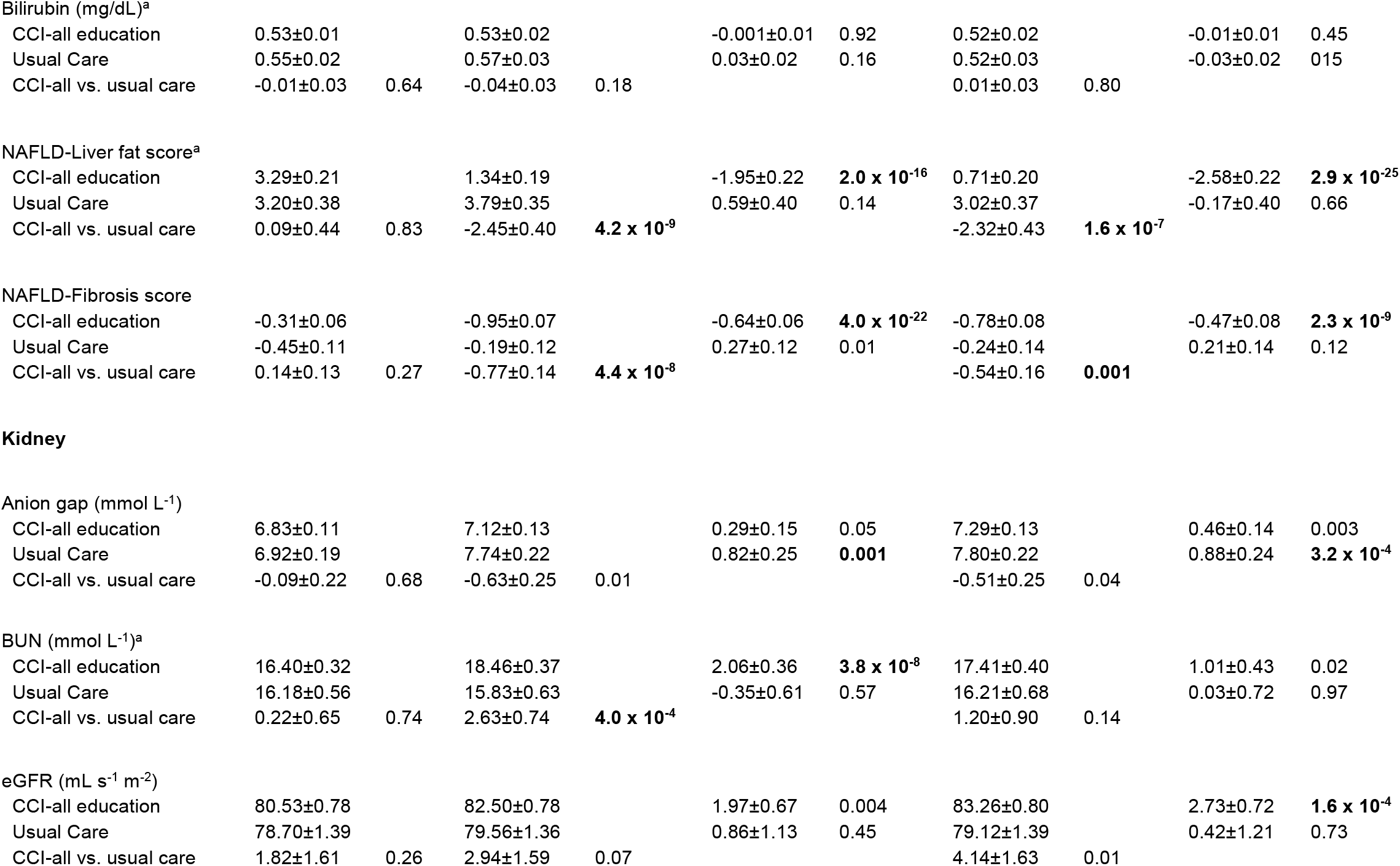
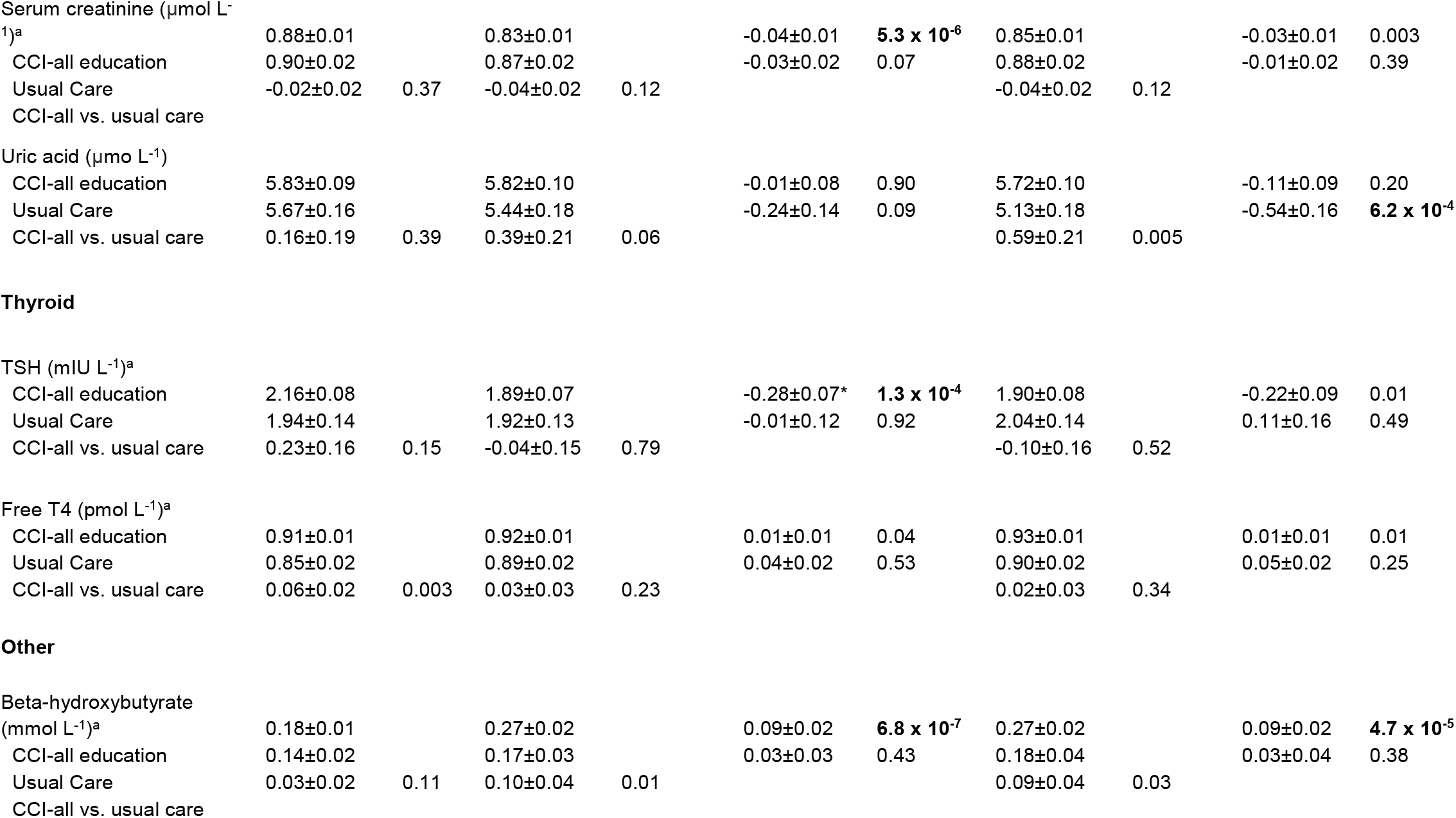
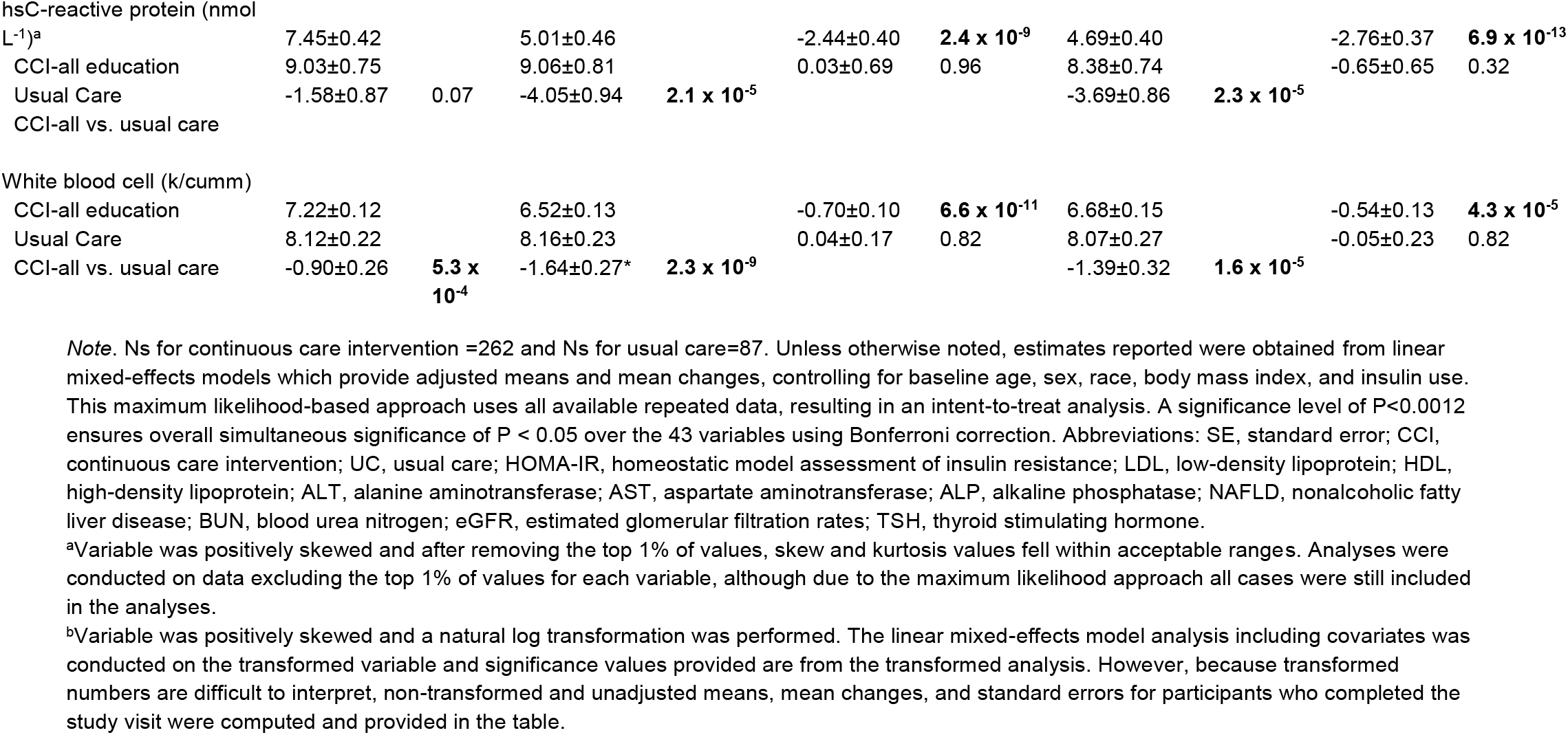
Adjusted mean changes over time

|  | Baseline |  | 1 Year |  |  |  | 2 Years |  |  |  |
| --- | --- | --- | --- | --- | --- | --- | --- | --- | --- | --- |
|  | Mean ± SE | P | Mean ± SE | P | Change from baseline | P | Meas ± SE | P | Change from baseline | P |
| **Glycemic** |  |  |  |  |  |  |  |  |  |  |
| Hemoglobin A1c (%) |  |  |  |  |  |  |  |  |  |  |
| CCI-all education | 7.7±0.1 |  | 6.3±0.1 |  | -1.3±0.1 | **6.6 x 10<sup>-38</sup>** | 6.7±0.1 |  | -0.9±0.1 | **1.8 x 10<sup>-17</sup>** |
| Usual Care | 7.5±0.2 |  | 7.6±0.1 |  | 0.2±0.2 | 0.31 | 7.9±0.2 |  | 0.4±0.2 | 0.02 |
| CCI-all vs. usual care | 0.2±0.2 | 0.28 | -1.3±0.2 | **2.7 x 10<sup>-14</sup>** |  |  | -1.2±0.2 | **1.3 x 10<sup>-9</sup>** |  |  |
| C-Peptide (nmol L<sup>-1</sup>) |  |  |  |  |  |  |  |  |  |  |
| CCI-all education | 4.33±0.13 |  | 3.27±0.14 |  | -1.06±0.13 | **7.3 x 10<sup>-14</sup>** | 3.16±0.12 |  | -1.17±0.13 | **2.2 x 10<sup>-16</sup>** |
| Usual Care | 4.39±0.24 |  | 4.38±0.25 |  | -0.004±0.24 | 0.99 | 3.89±0.22 |  | -0.49±0.24 | 0.04 |
| CCI-all vs. usual care | -0.06±0.28 | 0.84 | -1.12±0.28 | **9.8 x 10<sup>-5</sup>** |  |  | -0.73±0.26 | 5.0 x 10<sup>-3</sup> |  |  |
| Fasting glucose (mg/dL) |  |  |  |  |  |  |  |  |  |  |
| CCI-all education | 163.67±3.90 |  | 127.29±3.62 |  | -36.39±4.47 | **1.0 x 10<sup>-14</sup>** | 134.58±4.13 |  | - | **6.8 x 10<sup>-9</sup>** |
| Usual Care | 151.21±6.93 |  | 160.58±6.17 |  | 9.38±7.61 | 0.22 | 172.89±7.00 |  | 29.10±4.88 | 0.01 |
| CCI-all vs. usual care | 12.47±8.02 | 0.12 | -33.30±7.24 | **6.3 x 10<sup>-6</sup>** |  |  | -38.31±8.21 | **4.8 x 10<sup>-6</sup>** | 21.68±8.28 |  |
| Fasting Insulin (pmol L<sup>-1</sup>)<sup>a</sup> |  |  |  |  |  |  |  |  |  |  |
| CCI-all education | 27.73±1.26 |  | 16.47±1.13 |  | -11.26±1.28 | **3.2 x 10<sup>-16</sup>** | 16.02±1.02 |  | - | **2.2 x 10<sup>-18</sup>** |
| Usual Care | 27.57±2.29 |  | 26.47±2.06 |  | -1.10±2.30 | 0.63 | 24.17±1.84 |  | 11.71±1.25 | 0.13 |
| CCI-all vs. usual care | 0.16±2.63 | 0.95 | -10.00±2.38 | **3.6 x 10<sup>-5</sup>** |  |  | -8.15±2.14 | **1.7 x 10<sup>-4</sup>** | -3.40±2.22 |  |
| HOMA-IR (insulin derived), all<sup>a</sup> | 9.09±0.41 |  | 4.85±0.39 |  | -4.24±0.45 | **3.5 x 10<sup>-18</sup>** | 5.27±0.44 |  | -3.82±0.49 | **3.8 x 10<sup>-13</sup>** |
| CCI-all education | 9.58±0.73 |  | 10.33±0.73 |  | 0.75±0.81 | 0.35 | 9.95±0.77 |  | 0.37±0.83 | 0.66 |
| Usual Care | -0.49±0.85 | 0.57 | -5.48±0.84 | **2.9 x 10<sup>-10</sup>** |  |  | -4.67±0.89 | **3.4 x 10<sup>-7</sup>** |  |  |
| CCI-all vs. usual care |  |  |  |  |  |  |  |  |  |  |
| HOMA-IR (insulin derived), excluding exogenous users<sup>a</sup> | 9.08±0.46 |  | 4.56±0.44 |  | -4.53±0.47 | **6.5 x 10<sup>-18</sup>** | 5.25±0.38 |  | -3.83±0.49 | **2.7 x 10<sup>-13</sup>** |
| CCI-all education | 8.66±0.92 |  | 10.87±0.98 |  | 2.21±1.02 | 0.03 | 8.26±0.75 |  | -0.40±0.94 | 0.68 |
| Usual Care | 0.43±1.03 | 0.68 | -6.31±1.08 | **2.2 x 10<sup>-8</sup>** |  |  | -3.01±0.85 | **5.4 x 10<sup>-4</sup>** |  |  |
| CCI-all vs. usual care |  |  |  |  |  |  |  |  |  |  |
| HOMA-IR (C-peptide derived), all<sup>a</sup> | 11.25±0.37 |  | 8.07±0.38 |  | -3.19±0.39 | **1.8 x 10<sup>-14</sup>** | 7.88±0.35 |  | -3.37±0.39 | **1.1 x 10<sup>-15</sup>** |
| CCI-all education | 11.04±0.67 |  | 11.81±0.71 |  | 0.77±0.72 | 0.28 | 10.62±0.64 |  | -0.42±0.70 | 0.55 |
| Usual Care | 0.21±0.77 | 0.78 | -3.75±0.81 | **5.8 x 10<sup>-6</sup>** |  |  | -2.74±0.74 | **2.5 x 10<sup>-4</sup>** |  |  |
| CCI-all vs. usual care |  |  |  |  |  |  |  |  |  |  |
| Metabolic and Body Composition |  |  |  |  |  |  |  |  |  |  |
| Weight-clinic (kg) |  |  |  |  |  |  |  |  |  |  |
| CCI-all education | 114.56±0.60 |  | 100.27±0.86 |  | -14.29±0.71 | **9.7 x 10<sup>-56</sup>** | 102.62±1.10 |  | - | **8.8 x 10<sup>-28</sup>** |
| Usual Care | 111.07±1.09 |  | 111.71±1.47 |  | 0.64±1.17 | 0.58 | 112.35±1.90 |  | 11.94±0.96 | 0.43 |
| CCI-all vs. usual care | 3.49±1.27 | 0.01 | -11.44±1.71 | **1.4 x 10<sup>-10</sup>** |  |  | -9.73±2.20 | **1.5 x 10<sup>-5</sup>** | 1.28±1.63 |  |
| Spine bone mineral density (kg) | 1.21±0.01 | — | 1.22±0.01 | — | 0.01±0.01 | 0.11 | 1.22±0.01 | — | 0.01±0.01 | 0.02 |
| CCI-all education |  |  |  |  |  |  |  |  |  |  |
| Central abdominal fat (kg) |  |  |  |  |  |  |  |  |  |  |
| CCI-all education | 5.89±0.07 | — | 4.62±0.08 | — | -1.27±0.07 | **1.3 x 10<sup>-42</sup>** | 4.99±0.10 | — | -0.90±0.08 | **1.6 x 10<sup>-21</sup>** |
| Android: gynoid ratio |  |  |  |  |  |  |  |  |  |  |
| CCI-all education | 1.27±0.02 | — | 1.18±0.02 | — | -0.09±0.1 | **2.4 x 10<sup>-13</sup>** | 1.20±0.02 | — | -0.07±0.01 | **4.7 x 10<sup>-8</sup>** |
| Lower extremities lean mass (kg) |  |  |  |  |  |  |  |  |  |  |
| CCI-all education | 18.74±0.16 | — | 17.41±0.15 | — | -1.33±0.10 | **5.9 x 10<sup>-31</sup>** | 17.38±0.17 | — | -1.36±0.12 | **1.3 x 10<sup>-21</sup>** |
| **Cardiovascular** |  |  |  |  |  |  |  |  |  |  |
| Systolic blood pressure (mmHg) | 131.7±0.9 |  | 125.3±0.9 |  | -6.5±1.1 | **3.3 x 10<sup>-8</sup>** | 125.9±1.0 |  | -5.8±1.2 | **2.4 x 10<sup>-6</sup>** |
| CCI-all education | 130.3±1.6 |  | 129.5±1.6 |  | -0.9±1.9 | 0.66 | 129.9±1.8 |  | -0.5±2.1 | 0.83 |
| Usual Care | 1.4±1.8 | 0.43 | -4.2±1.8 | 0.02 |  |  | -3.9±2.1 | 0.06 |  |  |
| CCI-all vs. usual care |  |  |  |  |  |  |  |  |  |  |
| Diastolic blood pressure (mmHg) | 81.8±0.5 |  | 78.1±0.6 |  | -3.7±0.7 | **5.4 x 10<sup>-8</sup>** | 78.7±0.6 |  | -3.1±0.7 | **3.3 x 10<sup>-5</sup>** |
| CCI-all education | 82.1±1.0 |  | 81.3±1.0 |  | -0.8±1.1 | 0.47 | 81.6±1.1 |  | -0.6±1.3 | 0.65 |
| Usual Care | -0.3±1.1 | 0.76 | -3.2±1.1 | 0.41 |  |  | -2.8±1.3 | 0.03 |  |  |
| CCI-all vs. usual care |  |  |  |  |  |  |  |  |  |  |
| Total cholesterol (mg/dL) |  |  |  |  |  |  |  |  |  |  |
| CCI-all education | 184.4±2.7 |  | 192.8±3.4 |  | 8.4±3.1 | 0.01 | 194.1±3.5 |  | 9.7±3.6 | 0.01 |
| Usual Care | 181.2±4.9 |  | 179.4±6.1 |  | -1.8±5.5 | 0.75 | 180.9±6.2 |  | -0.3±6.4 | 0.96 |
| CCI-all vs. usual care | 3.3±5.7 | 0.57 | 13.5±7.0 | 0.06 |  |  | 13.3±7.2 | 0.07 |  |  |
| LDL-cholesterol (mg/dL) |  |  |  |  |  |  |  |  |  |  |
| CCI-all education | 103.5±2.2 |  | 114.1±2.5 |  | 10.6±2.5 | **2.5 x 10<sup>-5</sup>** | 114.6±2.8 |  | 11.1±2.8 | **1.1x 10<sup>-4</sup>** |
| Usual Care | 100.0±4.2 |  | 88.9±4.9 |  | -11.2±4.7 | 0.02 | 90.9±5.1 |  | -9.1±5.1 | 0.08 |
| CCI-all vs. usual care | 3.6±4.8 | 0.46 | 25.2±5.6 | **8.9 x 10<sup>-6</sup>** |  |  | 23.7±5.9 | **7.0x 10<sup>-5</sup>** |  |  |
| HDL-cholesterol (mg/dL) |  |  |  |  |  |  |  |  |  |  |
| CCI-all education | 41.8±0.9 |  | 49.5±0.9 |  | 7.8±0.8 | **4.4 x 10<sup>-19</sup>** | 49.5±1.0 |  | 7.8±0.9 | **2.7 x 10<sup>-16</sup>** |
| Usual Care | 38.7±1.4 |  | 37.2±1.7 |  | -1.5±1.4 | 0.30 | 42.5±1.7 |  | 3.8±1.6 | 0.02 |
| CCI-all vs. usual care | 3.1±1.6 | 0.06 | 12.4±2.0 | **1.1 x 10<sup>-9</sup>** |  |  | 7.1±2.0 | **4.1x 10<sup>-4</sup>** |  |  |
| Triglycerides (mg/dL)<sup>b</sup> |  |  |  |  |  |  |  |  |  |  |
| CCI-all education | 197.2±9.1 |  | 148.9±10.1 |  | -48.3±13.7 | **7.4 x 10<sup>-16</sup>** | 153.3±10.4 |  | -43.9±14.0 | **6.2 x 10<sup>-9</sup>** |
| Usual Care | 282.9±45.1 |  | 314.5±61.4 |  | 31.6±74.6 | 0.35 | 209.5±18.5 |  | -73.4±55.9 | 0.75 |
| CCI-all vs. usual care | -85.7±30.1 | 0.09 | -165.5±39.0 | **1.5 x 10<sup>-8</sup>** |  |  | -56.2±19.0 | **7.1 x 10<sup>-5</sup>** |  |  |
| Liver |  |  |  |  |  |  |  |  |  |  |
| ALT (Units/L)<sup>a</sup> |  |  |  |  |  |  |  |  |  |  |
| CCI-all education | 29.16±0.97 |  | 21.53±0.88 |  | -7.63±1.02 | **7.7 x 10<sup>-13</sup>** | 23.00±0.91 |  | -6.16±0.95 | **4.0 x 10<sup>-10</sup>** |
| Usual Care | 25.84±1.72 |  | 26.98±1.51 |  | 1.14±1.73 | 0.51 | 26.80±1.57 |  | 0.96±1.62 | 0.56 |
| CCI-all vs. usual care | 3.31±1.99 | 0.10 | -5.45±1.77 | 0.002 |  |  | -3.80±1.84 | 0.04 |  |  |
| AST (Units/L)<sup>a</sup> |  |  |  |  |  |  |  |  |  |  |
| CCI-all education | 22.50±0.64 |  | 19.07±0.58 |  | -3.43±0.69 | **1.1 x 10<sup>-6</sup>** | 19.78±0.57 |  | -2.72±0.66 | **5.1 x 10<sup>-5</sup>** |
| Usual Care | 21.51±1.13 |  | 23.37±1.00 |  | 1.86±1.19 | 0.12 | 23.19±0.99 |  | 1.68±1.14 | 0.14 |
| CCI-all vs. usual care | 0.99±1.31 | 0.45 | -4.30±1.17 | **2.8 x 10<sup>-4</sup>** |  |  | -3.41±1.16 | 3.5 x 10<sup>-3</sup> |  |  |
| ALP (Units/L) |  |  |  |  |  |  |  |  |  |  |
| CCI-all education | 74.13±1.42 |  | 64.34±1.44 |  | -9.78±0.98 | **1.9 x 10<sup>-20</sup>** | 64.50±1.58 |  | -9.63±1.19* | **1.8 x 10<sup>-14</sup>** |
| Usual Care | 78.55±2.53 |  | 79.05±2.55 |  | 0.50±1.65 | 0.76 | 82.47±2.76 |  | 3.92±2.00 | 0.05 |
| CCI-all vs. usual care | -4.42±2.94 | 0.13 | -14.71±2.97 | **1.2 x 10<sup>-6</sup>** |  |  | -17.97±3.22 | **5.1 x 10<sup>-8</sup>** |  |  |
| Bilirubin (mg/dL)<sup>a</sup> |  |  |  |  |  |  |  |  |  |  |
| CCI-all education | 0.53±0.01 |  | 0.53±0.02 |  | -0.001±0.01 | 0.92 | 0.52±0.02 |  | -0.01±0.01 | 0.45 |
| Usual Care | 0.55±0.02 |  | 0.57±0.03 |  | 0.03±0.02 | 0.16 | 0.52±0.03 |  | -0.03±0.02 | 0.15 |
| CCI-all vs. usual care | -0.01±0.03 | 0.64 | -0.04±0.03 | 0.18 |  |  | 0.01±0.03 | 0.80 |  |  |
| NAFLD-Liver fat score<sup>a</sup> |  |  |  |  |  |  |  |  |  |  |
| CCI-all education | 3.29±0.21 |  | 1.34±0.19 |  | -1.95±0.22 | **2.0 x 10<sup>-16</sup>** | 0.71±0.20 |  | -2.58±0.22 | **2.9 x 10<sup>-25</sup>** |
| Usual Care | 3.20±0.38 |  | 3.79±0.35 |  | 0.59±0.40 | 0.14 | 3.02±0.37 |  | -0.17±0.40 | 0.66 |
| CCI-all vs. usual care | 0.09±0.44 | 0.83 | -2.45±0.40 | **4.2 x 10<sup>-9</sup>** |  |  | -2.32±0.43 | **1.6 x 10<sup>-7</sup>** |  |  |
| NAFLD-Fibrosis score |  |  |  |  |  |  |  |  |  |  |
| CCI-all education | -0.31±0.06 |  | -0.95±0.07 |  | -0.64±0.06 | **4.0 x 10<sup>-22</sup>** | -0.78±0.08 |  | -0.47±0.08 | **2.3 x 10<sup>-9</sup>** |
| Usual Care | -0.45±0.11 |  | -0.19±0.12 |  | 0.27±0.12 | 0.01 | -0.24±0.14 |  | 0.21±0.14 | 0.12 |
| CCI-all vs. usual care | 0.14±0.13 | 0.27 | -0.77±0.14 | **4.4 x 10<sup>-8</sup>** |  |  | -0.54±0.16 | **0.001** |  |  |
| Kidney |  |  |  |  |  |  |  |  |  |  |
| Anion gap (mmol L<sup>-1</sup>) |  |  |  |  |  |  |  |  |  |  |
| CCI-all education | 6.83±0.11 |  | 7.12±0.13 |  | 0.29±0.15 | 0.05 | 7.29±0.13 |  | 0.46±0.14 | 0.003 |
| Usual Care | 6.92±0.19 |  | 7.74±0.22 |  | 0.82±0.25 | **0.001** | 7.80±0.22 |  | 0.88±0.24 | **3.2 x 10<sup>-4</sup>** |
| CCI-all vs. usual care | -0.09±0.22 | 0.68 | -0.63±0.25 | 0.01 |  |  | -0.51±0.25 | 0.04 |  |  |
| BUN (mmol L<sup>-1</sup>)<sup>a</sup> |  |  |  |  |  |  |  |  |  |  |
| CCI-all education | 16.40±0.32 |  | 18.46±0.37 |  | 2.06±0.36 | **3.8 x 10<sup>-8</sup>** | 17.41±0.40 |  | 1.01±0.43 | 0.02 |
| Usual Care | 16.18±0.56 |  | 15.83±0.63 |  | -0.35±0.61 | 0.57 | 16.21±0.68 |  | 0.03±0.72 | 0.97 |
| CCI-all vs. usual care | 0.22±0.65 | 0.74 | 2.63±0.74 | **4.0 x 10<sup>-4</sup>** |  |  | 1.20±0.90 | 0.14 |  |  |
| eGFR (mL s<sup>-1</sup> m<sup>-2</sup>) |  |  |  |  |  |  |  |  |  |  |
| CCI-all education | 80.53±0.78 |  | 82.50±0.78 |  | 1.97±0.67 | 0.004 | 83.26±0.80 |  | 2.73±0.72 | **1.6 x 10<sup>-4</sup>** |
| Usual Care | 78.70±1.39 |  | 79.56±1.36 |  | 0.86±1.13 | 0.45 | 79.12±1.39 |  | 0.42±1.21 | 0.73 |
| CCI-all vs. usual care | 1.82±1.61 | 0.26 | 2.94±1.59 | 0.07 |  |  | 4.14±1.63 | 0.01 |  |  |
| Serum creatinine (μmol L<sup>-1</sup>)<sup>a</sup> |  |  |  |  |  |  |  |  |  |  |
|  | 0.88±0.01 |  | 0.83±0.01 |  | -0.04±0.01 | **5.3 x 10<sup>-6</sup>** | 0.85±0.01 |  | -0.03±0.01 | 0.003 |
| CCI-all education | 0.90±0.02 |  | 0.87±0.02 |  | -0.03±0.02 | 0.07 | 0.88±0.02 |  | -0.01±0.02 | 0.39 |
| Usual Care | -0.02±0.02 | 0.37 | -0.04±0.02 | 0.12 |  |  | -0.04±0.02 | 0.12 |  |  |
| CCI-all vs. usual care |  |  |  |  |  |  |  |  |  |  |
| Uric acid (μmo L<sup>-1</sup>) |  |  |  |  |  |  |  |  |  |  |
| CCI-all education | 5.83±0.09 |  | 5.82±0.10 |  | -0.01±0.08 | 0.90 | 5.72±0.10 |  | -0.11±0.09 | 0.20 |
| Usual Care | 5.67±0.16 |  | 5.44±0.18 |  | -0.24±0.14 | 0.09 | 5.13±0.18 |  | -0.54±0.16 | **6.2 x 10<sup>-4</sup>** |
| CCI-all vs. usual care | 0.16±0.19 | 0.39 | 0.39±0.21 | 0.06 |  |  | 0.59±0.21 | 0.005 |  |  |
| Thyroid |  |  |  |  |  |  |  |  |  |  |
| TSH (mIU L<sup>-1</sup>)<sup>a</sup> |  |  |  |  |  |  |  |  |  |  |
| CCI-all education | 2.16±0.08 |  | 1.89±0.07 |  | -0.28±0.07* | **1.3 x 10<sup>-4</sup>** | 1.90±0.08 |  | -0.22±0.09 | 0.01 |
| Usual Care | 1.94±0.14 |  | 1.92±0.13 |  | -0.01±0.12 | 0.92 | 2.04±0.14 |  | 0.11±0.16 | 0.49 |
| CCI-all vs. usual care | 0.23±0.16 | 0.15 | -0.04±0.15 | 0.79 |  |  | -0.10±0.16 | 0.52 |  |  |
| Free T4 (pmol L<sup>-1</sup>)<sup>a</sup> |  |  |  |  |  |  |  |  |  |  |
| CCI-all education | 0.91±0.01 |  | 0.92±0.01 |  | 0.01±0.01 | 0.04 | 0.93±0.01 |  | 0.01±0.01 | 0.01 |
| Usual Care | 0.85±0.02 |  | 0.89±0.02 |  | 0.04±0.02 | 0.53 | 0.90±0.02 |  | 0.05±0.02 | 0.25 |
| CCI-all vs. usual care | 0.06±0.02 | 0.003 | 0.03±0.03 | 0.23 |  |  | 0.02±0.03 | 0.34 |  |  |
| Other |  |  |  |  |  |  |  |  |  |  |
| Beta-hydroxybutyrate (mmol L<sup>-1</sup>)<sup>a</sup> |  |  |  |  |  |  |  |  |  |  |
|  | 0.18±0.01 |  | 0.27±0.02 |  | 0.09±0.02 | **6.8 x 10<sup>-7</sup>** | 0.27±0.02 |  | 0.09±0.02 | **4.7 x 10<sup>-5</sup>** |
| CCI-all education | 0.14±0.02 |  | 0.17±0.03 |  | 0.03±0.03 | 0.43 | 0.18±0.04 |  | 0.03±0.04 | 0.38 |
| Usual Care | 0.03±0.02 | 0.11 | 0.10±0.04 | 0.01 |  |  | 0.09±0.04 | 0.03 |  |  |
| CCI-all vs. usual care |  |  |  |  |  |  |  |  |  |  |
| hsC-reactive protein (nmol L<sup>-1</sup>)<sup>a</sup> | 7.45±0.42 |  | 5.01±0.46 |  | -2.44±0.40 | **2.4 x 10<sup>-9</sup>** | 4.69±0.40 |  | -2.76±0.37 | **6.9 x 10<sup>-13</sup>** |
| CCI-all education | 9.03±0.75 |  | 9.06±0.81 |  | 0.03±0.69 | 0.96 | 8.38±0.74 |  | -0.65±0.65 | 0.32 |
| Usual Care | -1.58±0.87 | 0.07 | -4.05±0.94 | **2.1 x 10<sup>-5</sup>** |  |  | -3.69±0.86 | **2.3 x 10<sup>-5</sup>** |  |  |
| CCI-all vs. usual care |  |  |  |  |  |  |  |  |  |  |
| White blood cell (k/cumm) |  |  |  |  |  |  |  |  |  |  |
| CCI-all education | 7.22±0.12 |  | 6.52±0.13 |  | -0.70±0.10 | **6.6 x 10<sup>-11</sup>** | 6.68±0.15 |  | -0.54±0.13 | **4.3 x 10<sup>-5</sup>** |
| Usual Care | 8.12±0.22 |  | 8.16±0.23 |  | 0.04±0.17 | 0.82 | 8.07±0.27 |  | -0.05±0.23 | 0.82 |
| CCI-all vs. usual care | -0.90±0.26 | **5.3 x 10<sup>-4</sup>** | -1.64±0.27* | **2.3 x 10<sup>-9</sup>** |  |  | -1.39±0.32 | **1.6 x 10<sup>-5</sup>** |  |  |

*Note.* Ns for continuous care intervention =262 and Ns for usual care=87. Unless otherwise noted, estimates reported were obtained from linear mixed-effects models which provide adjusted means and mean changes, controlling for baseline age, sex, race, body mass index, and insulin use. This maximum likelihood-based approach uses all available repeated data, resulting in an intent-to-treat analysis. A significance level of P<0.0012 ensures overall simultaneous significance of P < 0.05 over the 43 variables using Bonferroni correction. Abbreviations: SE, standard error; CCI, continuous care intervention; UC, usual care; HOMA-IR, homeostatic model assessment of insulin resistance; LDL, low-density lipoprotein; HDL, high-density lipoprotein; ALT, alanine aminotransferase; AST, aspartate aminotransferase; ALP, alkaline phosphatase; NAFLD, nonalcoholic fatty liver disease; BUN, blood urea nitrogen; eGFR, estimated glomerular filtration rates; TSH, thyroid stimulating hormone.
<sup>a</sup>Variable was positively skewed and after removing the top 1% of values, skew and kurtosis values fell within acceptable ranges. Analyses were conducted on data excluding the top 1% of values for each variable, although due to the maximum likelihood approach all cases were still included in the analyses.
<sup>b</sup>Variable was positively skewed and a natural log transformation was performed. The linear mixed-effects model analysis including covariates was conducted on the transformed variable and significance values provided are from the transformed analysis. However, because transformed numbers are difficult to interpret, non-transformed and unadjusted means, mean changes, and standard errors for participants who completed the study visit were computed and provided in the table.

### Glycemic outcomes

From baseline to 2 years (Table 2), significant reductions in HbA1c (0.9% unit decrease, −12% relative to baseline, P=1.8×10^-17^; Figure 2A), C-peptide (−27%, P=2.2×10^-16^), fasting glucose (−18%, P=6.8×10^-9^), fasting insulin (−42%, P=2.2×10^-18^, Figure 2B), insulin-derived HOMA-IR excluding exogenous insulin users (−42%, P=2.7×10^-13^), and C-peptide-derived HOMA-IR (−30%, P=1.1×10^-15^) were observed in the CCI group, whereas no changes occurred in the UC group (Supplementary Figures 1A and 1B) (Table 2). There were also significant between-group (CCI vs. UC) differences observed at 2 years in HbA1c, fasting glucose, fasting insulin, insulin-derived HOMA-IR excluding exogenous users, and C-peptide-derived HOMA-IR, with the CCI group having lower glycemic marker means (Table 2).

**Figure 2.**
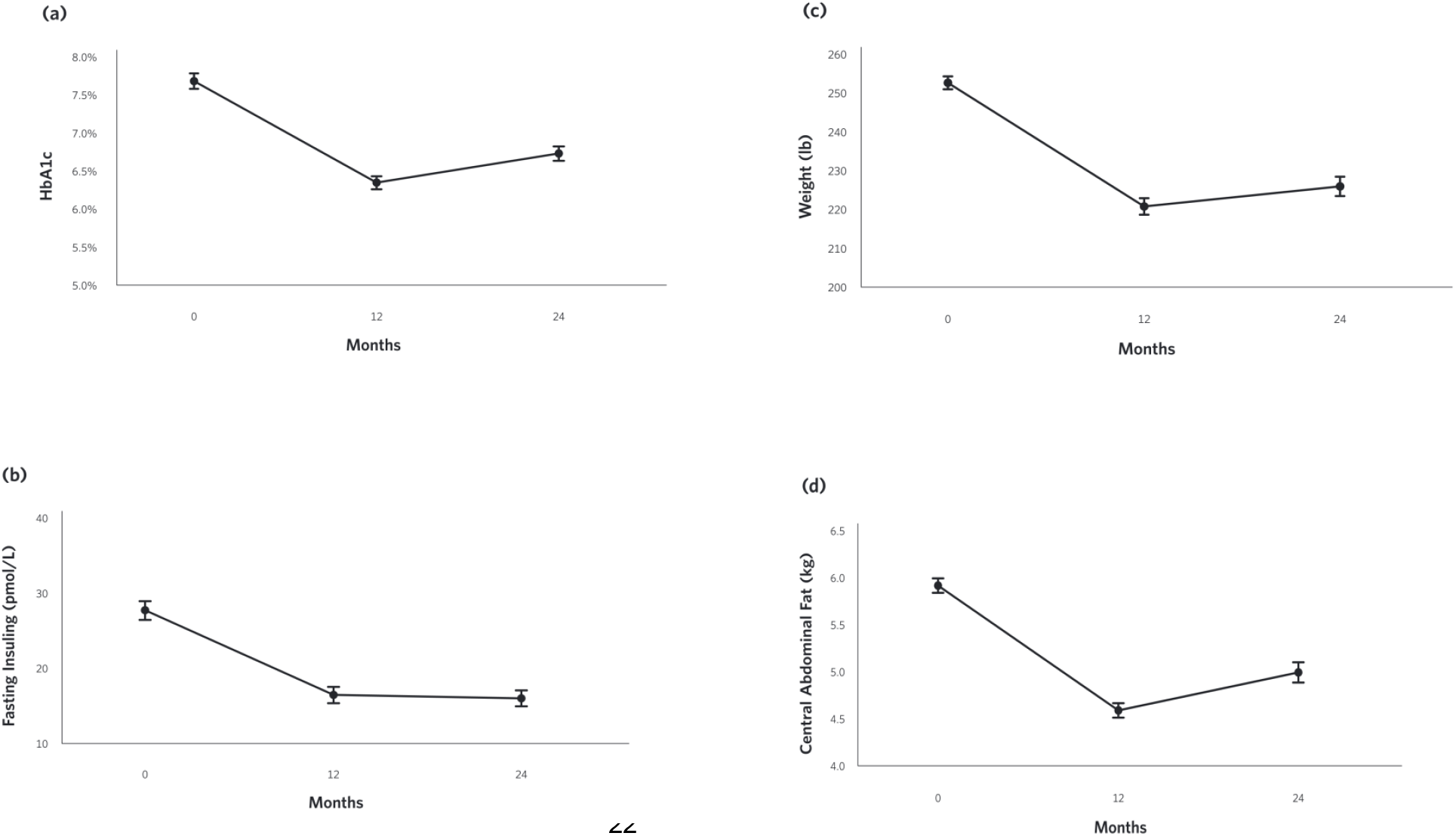
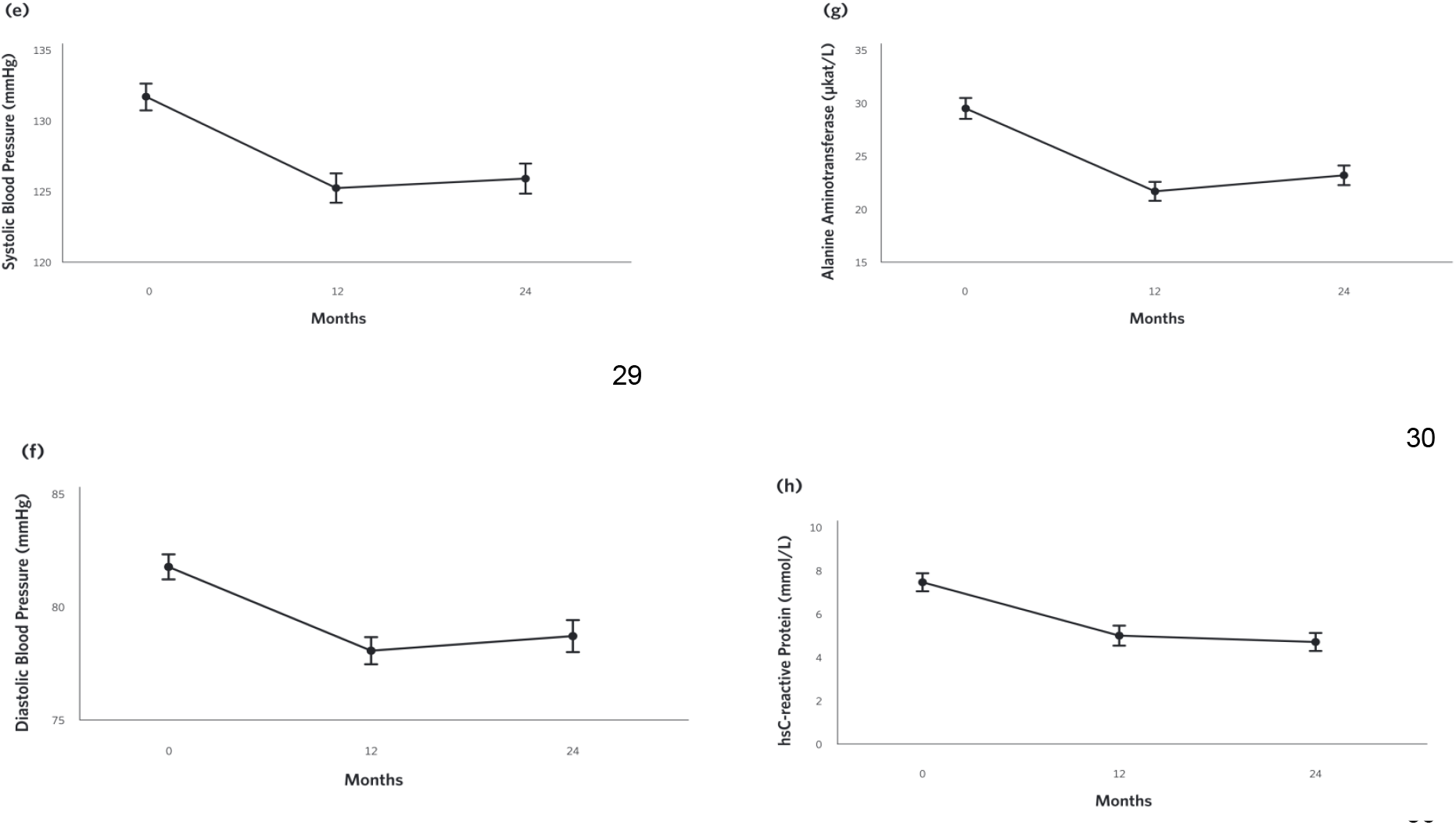
Adjusted mean changes from baseline to 2-years in the CCI group for (A) HbA1c, (B) Fasting insulin, (C) Weight, (D) Central Abdominal Fat [CAF], (E) Systolic Blood Pressure, (F) Diastolic Blood Pressure (G) Alanine aminotransferase (ALT), and (H) High sensitive C-reactive protein (hsCRP).

### Metabolic and body composition outcomes

At 2 years, mean weight change from baseline was −10% (P=8.8×10^-28^; Figure 2C) in the CCI group, whereas no change was observed in the UC group (Supplementary Figure 1C). Among CCI patients, 74% had ≥ 5% weight loss compared to only 14% of UC patients (Supplementary Figure 2; completers analysis). Consistent with the weight loss observed, the CCI group had reductions in abdominal fat content, with decreases in CAF (−15%, P=1.6×10^-21^, Figure 2D) and the A/G ratio (−6%, P=4.7×10^-8^) from baseline to 2 years (Table 2). The CCI group’s total spine BMD remained unchanged from baseline to 2 years after correction for multiple comparisons (Table 2). The changes in the average LELM in the CCI are included in the Table 2, and further elaborated in the **supplementary data (Discussion section)**.

### Cardiovascular risk factor outcomes

Decreases in systolic (−4%, P= 2.4×10^-6^, Figure 2E) and diastolic (−4%, P= 3.3×10^-5^, Figure 2F) blood pressures and triglycerides (−22%, P=6.2×10^-9^) were observed in the CCI but not UC group at 2 years (Table 2, Supplementary Figures 3A and 3B). The CCI group’s HDL-cholesterol (+19%, P= 2.7×10^-16^) and LDL-cholesterol (+11%, P=1.1×10^-4^) both increased from baseline to two years, whereas no changes were observed in the UC group (Table 2). No changes in total cholesterol were observed in either the CCI or UC group. At 2 years, the CCI group had higher HDL-cholesterol, higher LDL-cholesterol, and lower triglycerides than UC. No between-group differences were observed at 2 years for systolic or diastolic blood pressure or total cholesterol (Table 2).

**Figure 3.**
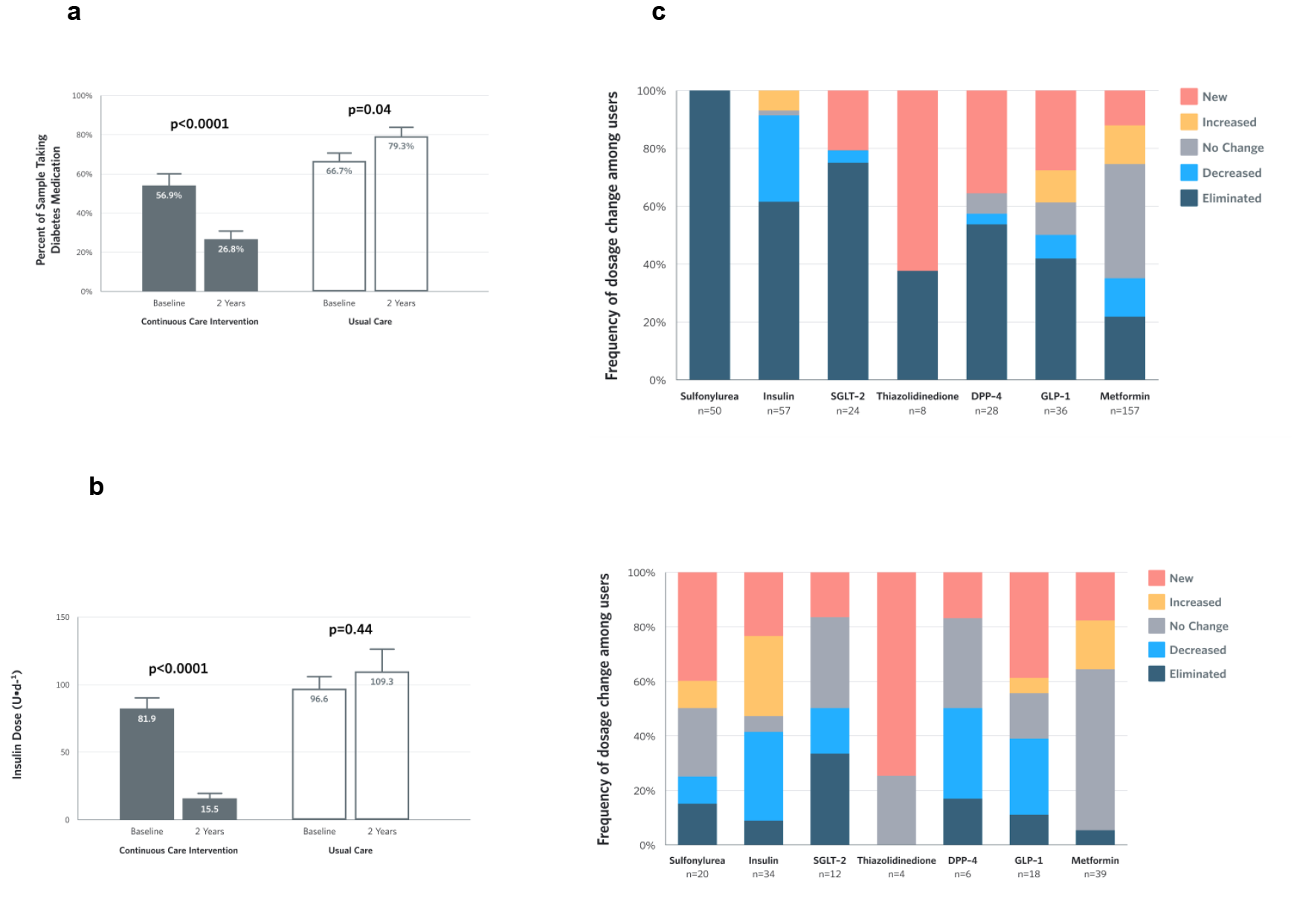
Medication and insulin dose changes from baseline to 2 years for CCI and UC group completers. (A) Percent of completers taking diabetes medications, excluding metformin. (B) Mean + SE prescribed insulin dose among baseline users. (C) Frequency of medication dosage and use change among prescribed users by diabetes medication class.

### Liver-related outcomes

From baseline to 2 years, the CCI group’s ALT (−21%, P=4.0×10^-10^; Table 2, Figure 2G), AST (-12%, P=5.1×10^-5^), ALP (−13%, P=1.8×10^-14^), NLF (−78%, P=2.9×10^-25^) and NFS (−60%, P=2.3×10^-9^) were reduced, whereas no changes were observed in UC (e.g. ALT; Supplementary Figure 3C; Table 2). No bonferroni-corrected group differences were observed for bilirubin,ALT, nor AST at 2 years (Table 2).

### Kidney, thyroid, and inflammation outcomes

The eGFR increased in the CCI (+3%, P=1.6×10^-4^, Table 2) but not UC group at 2 years. The UC but not CCI group had increased anion gap and decreased uric acid (Table 2). No bonferroni-corrected within-group changes in BUN, serum creatinine, TSH, or Free T4 were observed in either the CCI or UC group from baseline to 2 years. No between-group differences were observed for any thyroid- or kidney-related markers at 2 years (Table 2).

From baseline to 2 years, decreases in the CCI group’s hsCRP (−37%, P=6.9×10^-13^, Table 2, Figure 2H) and white blood cell count (−7%, P=4.3×10^-5^) were observed. No changes were observed in the UC group (Supplementary Figure 3D). At 2 years, both markers of inflammation were lower in the CCI group compared to the UC group (Table 2).

### Diabetes Medication

All within-group changes in diabetes medication use among study completers appear in eTable 3 (ns are listed in the table). The proportion of CCI completers taking any diabetes medication (excluding metformin) decreased from 55.7% at baseline to 26.8% at 2 years (P=1.3×10^-11^, Figure 3A). Reductions in the use of diabetes medication classes included insulin (29.8% at baseline to 11.3% at 2 years, P=9.1×10^-9^) and sulfonylureas (23.7% at baseline and 0% at 2 years, P=4.2×10^-12^). At 2 years, no changes in the proportions of CCI completers taking SGLT-2 inhibitors (10.3% to 3.1%, P=0.01), DPP-4 (9.9% to 6.7%, P=0.42), GLP-1 agonists (13.4% to 10.8%, P=0.42), thiazolidinediones (1.5% to 2.6%, P=0.73), or metformin (71.4% to 63.9%, P=0.05) were observed after correction for multiple comparisons. No changes in use of any diabetes medication (excluding metformin) or individual diabetes medication classes were observed in the UC completers from baseline to 2 years. The mean dose for insulin-using participants at baseline decreased among CCI participants by 81% (P= 2.6×10^-12^) at 2 years, but not in UC participants (+13%, P=0.45) (see Figure 3B). For participants who remained insulin-users at 2 years, the mean dose also decreased in the CCI group by 61% (P=9.2×10^-5^) but not UC group (+19%, P=0.29). Among participants prescribed each diabetes medication class, the proportion with each dosage change (eliminated, reduced, unchanged, increased, or newly added) at 2 years in each group appears in Figure 3C.

### Disease Outcomes

All within-group changes and between-group differences in disease outcomes among the CCI and UC group participants appear in supplementary Table 4 (intent-to-treat analyses were conducted; all below n=262). The proportion of participants meeting the defined criteria for diabetes reversal at 2 years increased 41.4% (from 12.1% at baseline to 53.5% at 2 years, P<0.0×10^-36^) in the CCI group, whereas no Bonferroni-corrected change was observed in the UC group (7.1% absolute decrease, P=0.04). In addition, diabetes remission (partial or complete) was observed in 46 (17.6%) participants in the CCI group and two (2.4%) of the UC participants at 2 years. Complete remission was observed in 17 (6.7%) CCI participants and none (0%) of the UC participants at 2 years.

At 2 years, 27.2% of CCI participants and 6.5% of UC patients showed resolution of metabolic syndrome. The proportion of participants with metabolic syndrome decreased from baseline to 2 years in the CCI (from 89.1% to 61.9%, P= 4.9×10^-15^) but not UC group. The two years improvements of suspected steatosis and fibrosis status are included in the supplementary Tables 4 and 5.

### Safety and adverse events

In the CCI group, there were no reported serious adverse events between one and two years attributed to the intervention or that resulted in discontinuation, including no reported episodes of ketoacidosis or severe hypoglycemia requiring assistance. Adverse events occurring in the first year of intervention (n=6) were previously reported[10]. Details of the adverse events are included in the supplementary data (Results section).

## Discussion

Following 2 years of a remote continuous care intervention supporting medical and lifestyle changes, the CCI participants demonstrated improved HbA1c, fasting glucose and insulin, and HOMA-IR. Pharmaceutical interventions of 1.5 to 3 years duration report HbA1c reductions of 0.2 to 1.0% with DPP-4 inhibitors, SGLT-2 inhibitors and GLP-1 agonists(20,21,S46–S48). The HbA1c reduction of 0.9% with this CCI is comparable to that observed in pharmaceutical trials, but is achieved while discontinuing 67.0% of diabetes-specific prescriptions including most insulin and all sulfonylureas that engender risks for weight gain and hypoglycemia(22,23). Comparable improvements in glycemic control and reduced medication were not observed in UC participants recruited from the same healthcare system, suggesting that the CCI improves diabetes management relative to usual care. Other interventions using carbohydrate restriction reported variable long-term glycemic improvement outcomes(24–26,S49–S51). The 0.9% absolute (12% relative) HbA1c reduction observed at 2 years is consistent with low carbohydrate studies reporting HbA1c reductions of 8-15% at 2 to 3.5 years (25,26,S49,S51) with medication reduction. Two others studies reported no changes in HbA1c from baseline to 2 years, even though the low carbohydrate arm reduced HbA1c in the first 6 months(24,S50). This study observed a modest increase in HbA1c and weight between 1 and 2 years in CCI participants suggesting some reduction in long-term effectiveness. Interestingly, insulin-levels show no regression toward baseline from 1 to 2 years indicating long-term improvement in hyperinsulinemia, an important component of diabetes pathology(8,27).

Criticisms of low-carbohydrate diets relate to poor adherence and long-term sustainability(16,28). In this CCI, self-monitoring combined with continuous remote-monitoring and feedback from the care team, including behavioral support and nutrition advice via the app, may have improved accountability and engagement(S52). In addition to glucose and weight tracking, dietary adherence was monitored by blood ketones. The 2 year BHB increase above baseline demonstrates sustained dietary modification. While laboratory BHB levels were increased from baseline, nutritional ketosis (≥0.5 mM) was observed in only a minority (14.1%) of participants at 2 years. On average, patient-measured BHB was ≥0.5 mM for 32.8% of measurements over the 2 years (eFigure 4). This reveals an opportunity to increase adherence to nutritional ketosis for patients not achieving their desired health outcomes while prompting future research investigating the association between dietary adherence and health improvements.

A majority of the CCI participants (53.5%) met criteria for diabetes reversal at 2 years while 17.6% achieved diabetes remission (i.e. glycemic control without medication use) based on intent-to-treat with multiple imputation. The percentage of all CCI enrollees (N=262) with verified reversal and remission requiring both completion of two years of the trial and an obtained laboratory value for HbA1c were 37.8% and 14.9%, respectively. CCI diabetes reversal exceeds remission as metformin prescriptions were usually continued given its role in preventing disease progression(7,29), preserving β-cell function(29) and in treatment of pre-diabetes per guidelines (28). Partial and complete remission rates of 2.4% and 0. 2% per year, respectively, have been reported in 122,781 T2D patients receiving standard diabetes care(3). The two-year remission rate (both partial and complete) in the CCI (17.6%) is higher than that achieved through intensive lifestyle intervention (ILI) in the Look AHEAD trial (9.2%)(4). Greater diabetes remission in the CCI versus Look AHEAD ILI could result from differences in the dietary intervention(14), patients’ ability to self-select their lifestyle or effectiveness of continuous remote care. Length of time with a T2D diagnosis is a factor in remission, with longer time since diagnosis resulting in lower remission(3,4,6,S53). Despite a mean of 8.4 years since diagnosis among CCI participants, the remission rate was higher than the Look AHEAD trial where its participants had a median of 5 years(4) since diabetes diagnosis.

Participants in the CCI achieved 10% mean weight loss (−11.9kg) at 2 years. CCI weight loss was comparable to observed weight loss following surgical gastric banding (−10.7kg) at 2 years(29). Previous studies consistently report that weight loss increases the likelihood of T2D remission(3,4,6). CCI participants also improved blood pressure, triglycerides, and HDL-cholesterol. Total cholesterol was unchanged and calculated LDL-cholesterol was increased at 2 years, but was not different from the LDL-cholesterol level observed at one year (+0.51, P=0.85). Despite the rise in LDL-cholesterol, the CCI cohort improved in 22 out of 26 CVD markers at one year(19). This includes a decrease in small LDL-particles and large VLDL-P and an increase in LDL-particle size with no changes in ApoB(19), a marker considered a better predictor of CVD risk than LDL-cholesterol(19,30,S54). Non-elevated LDL cholesterol values together with higher triglycerides and lower HDL-cholesterol are common in patients with abdominal obesity, T2D, and metabolic syndrome(31,S55,S56); these individuals often still have elevated atherogenic lipoproteins such as non-HDL(32,S57), small LDL particles(31,S58), and VLDL(31,S58). In the CCI group, non-HDL cholesterol did not change significantly from baseline to 2 years and several cardiovascular risk factors across various physiological systems improved, suggesting that the rise in LDL-cholesterol may not be associated with increased atherogenic risk(33).

The CCI group had a reduction in visceral fat content, CAF and A/G ratio. This is consistent with other low-carbohydrate interventions reporting visceral fat reduction as a component of weight loss(18,24,34,35,S59). Anatomical distribution of fat around the abdominal area (“android” obesity) is associated with T2D(36,S60) and other comorbidities such as metabolic syndrome(37) and NAFLD(38,S61). The alleviation of visceral fat in the CCI group was concurrent with resolution of metabolic syndrome at 2 years, while sustaining one-year improvements of liver enzymes(7), steatosis and fibrosis (39 in press, S62-S67). While studies in animal models(40,S68,S69) and children treated with ketogenic diets(41,S70) have suggested retardation in skeletal development and reduction in BMD, in this study of T2D adults the CCI group had no change in total spine BMD over two years. Our results are consistent with other adult ketogenic dietary studies that reported no bone mass loss in short-term(34,S71) or long-term follow-up of 2(35,S72) and 5(S73) years. The differing findings of ketogenic diet on bone mass between adults and children could be due to differential effects on developed and mineralized versus developing bones(42).

### Strengths and limitations

This study’s strengths include its size and prospective, longitudinal data collection from two participant groups (CCI and UC) which allowed statistical analysis by LMMs to investigate intervention time and treatment effects. While not randomized, the participants’ self-selection of intervention may contribute to the observed high retention and predicts real-life clinical management of chronic disease. The study also included patients prescribed insulin and with long-standing disease, groups often excluded from prior studies. The multi-component aspect of the intervention involving regular biomarker monitoring and access to a a remote care team may have improved the patients’ long-term dietary adherence and engagement. The dietary advice including encouraging participants to restrict carbohydrates, moderate protein intake, and eat to satiety may also help in maintaining long-term effectiveness. Weaknesses of this study include the lack of randomization and limited racial diversity. Interpretation of DXA body composition was limited to subregion analyses due to to the scanner not accomodating the patients’ complete body.

### Conclusions

At 2 years, the CCI, including remote medical management with instruction in nutritional ketosis, led to improvements in blood glucose, insulin, HbA1c, weight, blood pressure, triglycerides, liver function, and inflammation and reduced dependence upon medication. These long-term benefits were achieved concurrent with reduced prevalence of metabolic syndrome and visceral adiposity. The CCI had no adverse effect on bone mineral density. The CCI group also had higher prevalence of diabetes reversal and remission compared to the UC group following a standard diabetes care program. These results provide strong evidence that sustained improvement in diabetes status can be achieved through the continuous remote monitoring and accountability mechanisms provided by this multi-component CCI including recommendations for low carbohydrate nutrition.

## Supplementary Method

### Study Interventions

#### Continuous care intervention (CCI)

Briefly, participants in the CCI were provided access to a web-based software application (app), which was used to provide telemedicine communication, online resources and biomarker tracking tools. The participants used the app to upload and monitor their reportable biomarkers including body weight and blood glucose and beta-hydroxybutryrate (BHB). Biomarkers allowed for daily feedback to the care team and individualization of patient instruction. Frequency of reporting was personalized over time based on care needs. Participants were advised to achieve nutritional ketosis (blood BHB level at 0.5 to 3.0 mmol L^-1^) through sufficient carbohydrate restriction (initially <30g day^-1^ but gradually increased based on personal carbohydrate tolerance and health goals, primarily control of glucose and weight). Participants’ daily protein intake was initially targeted at a level of 1.5g kg^-1^ of a medium-frame ideal weight body and further individualized based on biomarkers. Participants were instructed to include sufficient dietary fat in meals to achieve satiety without tracking energy intake. Nutrition education directed consumption of monounsaturated and saturated fat with sufficient intake of omega-3 and omega-6 polyunsaturated fats. The participants were also encouraged to consume sufficient fluid, vitamins and minerals including sodium and magnesium, especially if signs of mineral deficiency were encountered (e.g. decreased circulating volume)(1,2).

The web-based app was also used by participants to communicate with their remote care team consisting of a health coach and medical provider. The remote care team provided education and support regarding dietary changes, behavior modification techniques for maintenance of lifestyle changes and actively directed changes for diabetes and antihypertensive medications as part of the intervention. Metformin prescriptions were continued except for contraindication, intolerance, or patient request given its efficacy for T2D prevention [3]. Education modules covered core concepts related to the dietary changes for achieving nutritional ketosis, and adaptation to and maintenance of the diet(1,2). Participants selected their preferred education mode (CCI-virtual, n=126 or CCI-onsite, n=136) during recruitment. The CCI-virtual group received care and education primarily via app-based communication. The CCI-onsite group received care and education via clinic-based group meetings (weekly for 12 weeks, bi-weekly for 12 weeks, monthly for 6 months, and then quarterly in the second year). All participants had access to the app for communication with their care team, online resources, biomarker tracking and the opportunity to participate in an online peer community for social support.

#### Usual Care (UC)

The participants recruited for usual care (UC) received care from their primary care physician or endocrinologist and were counseled by a registered dietician as part of a diabetes education program. These participants received the American Diabetes Association (ADA) recommendations on nutrition, lifestyle and diabetes management(3). No modification of their care was made for the study. This group was used as a reference control to study the effect of disease progression over 2 years in a cohort of participants prospectively recruited from the same geography and healthcare system. Figure 1 depicts the study flow from recruitment to 2 years post-enrollment.

### Body composition measures

The CCI participants’ total body composition was measured at baseline, one year and two years using dual-energy X-ray absorptiometry (DXA) (Lunar GE Prodigy, Madison, WI). Participants were scanned while wearing light clothing using standard clinical imaging procedures. The scans obtained were analyzed using GE Encore software(v11.10, Madison, WI). In many obese patients, full body scans were not obtained due to the scanner not accomodating the patient’s complete body resulting in issues such as cropping of the arms and/or overlapping of arms with the chest(4,5). To address these limitations, changes in bone density and fat and lean mass were assessed using subregions rather than the full body scan. We assessed changes in the bone mass by evaluating total spine bone mineral density (BMD) from baseline to 2 years(6). For assessment of fat mass, we manually selected the central abdominal fat (CAF) region using the software and evaluated the changes in CAF over time, as previously suggested for overweight individuals(4,7). Furthermore, we assessed changes in the android:gynoid (A/G) ratio by time. Due to lack of proper arm lean mass measurement, we analyzed the lower extremities lean mass (LELM) to assess weight-related changes in lean mass over time(8,9).

### Statistical analyses

All analyses were conducted using SPSS statistical software (Version 25.0, Armonk, NY). First, we examined the assumptions of normality and linearity. According to Kline’s (2011) (10) guidelines, 14 outcomes (i.e., fasting insulin, insulin and C-peptide-derived HOMA-IR scores, triglycerides, ALT, AST, bilirubin, N-LFS, BUN, serum creatinine, TSH, Free T4, hsCRP, and BHB) were positively skewed. We explored two approaches to handling the skewed variables: natural log-transformations and removing the top 1% of values. For N-LFS which includes both positive and negative values, a modulus log-transformation was performed instead of a natural log-transformation(11). For most variables, both approaches resulted in new skew and kurtosis values within the acceptable range. One variable (triglycerides) was only corrected via log-transformation, whereas two variables (C-peptide-derived HOMA-IR and TSH) were only corrected by removing the top 1% of values. For the other variables, we conducted sensitivity analyses to compare the two approaches. Because the results did not differ between the approaches and because interpretation of outcomes is more difficult with transformed variables, we report results from the approach of removing the top 1% of values for all variables except triglycerides. For triglycerides, analyses were performed and p-values reported on the log-transformed variable but the means and standard errors reported were computed from the untransformed variable. Next, we ran independent sample t-tests to examine differences in baseline characteristics between CCI and UC, and completers and dropouts.

We performed linear mixed-effects models (LMMs) to assess (1) within-group changes in the continuous study outcomes from baseline to 2 years and (2) between-group differences (CCI vs. UC) in the study outcomes at 2 years. The LMMs included fixed effects for time, group (CCI vs. UC), and a time by group interaction. Covariates included baseline age, sex, race (African American vs. other), BMI, and insulin use. This maximum likelihood-based approach uses all available repeated data, resulting in an intent-to-treat analysis. An unstructured covariance structure was specified for all models to account for correlations between repeated measures.

Within-group changes and between-group differences in dichotomous disease outcome variables; i.e., diabetes reversal, diabetes remission (partial or complete) and complete remission(12), metabolic syndrome(13,14), steatosis(15), fibrosis(16) were assessed, controlling for baseline age, sex, race, time since diagnosis, BMI, and insulin use. For this set of analyses, multiple imputation was used to replace missing values from baseline and 2 years with a set of plausible values, facilitating an intent-to-treat analysis (all ns=262). Missing values were estimated from 40 imputations (17) from logistic regression. Within-group changes from baseline to 2 years and between-group differences at 2 years were assessed using generalized estimating equations with binary logistic models and unstructured covariance matrices.

We also examined changes in participants’ diabetes medication use. First, we compared rates of diabetes medication use within groups from baseline to 2 years using McNemar’s test with continuity correction when appropriate. Next, we calculated the proportion of participants in each group with each diabetes medication class eliminated, reduced, not changed, increased, or added. Paired t-tests were used to assess within-group changes in insulin dosages from baseline to 2 years among participants taking insulin at baseline and among participants taking insulin at both baseline and 2 years.

We conducted a second set of the analyses with 2-year completers only. Results of the completers-only analyses appear in eTable 3 and 5. Given that 2 different modes (virtual and onsite) were utilized for delivery of the CCI group educational content, we also conducted another set of analyses to assess whether differences existed between the groups on all analyses of primary outcomes. As in our prior time points (1,18), no group differences were found; thus, the data from the two CCI educational groups were combined for this report. For all study analyses, nominal significance levels (P) are presented in the tables. A significance level of P<0.0012 ensures overall simultaneous significance of *P*<0.05 over the 43 variables using Bonferroni correction.

### Supplementary Results

#### Safety and adverse events

During the second year of intervention, nine adverse events were reported including: one breast cancer diagnosis, one mycosis fungoides, one onset of atrial fibrillation (Afib) with heart failure, one onset of migraine, two cases of chest pain (one resulting in stent placement), one pulmonary effusion, and two pulmonary embolisms (one following orthopedic surgery and one with benign ovarian mass/Afib). In the UC group, adverse events occuring in the first year were previously reported(1), and in the second year, adverse events occurred in six participants: one death from liver cancer, one hospitalization from recurrent seizure, one ureteropelvic junction obstruction from kidney stone, one cerebrovascular accident with left side weakness and sensory disturbances, one chest pain requiring percutaneous coronary intervention, and one deep vein thrombosis.

### Supplementary Discussion

#### Lower extremities lean mass (LELM)

In this study, the CCI group had a reduction (7.0%, 1.3kg) in the calculated LELM. Most lean mass loss was encountered in the first year without further reduction in year 2. Studies have reported that obese adults have about 20% higher thigh muscle mass than those with normal weight(19,20). The reduced upper body load burden achieved through weight loss might explain the reduction of LELM. This reflects an appropriate post-weight decrease in muscle mass rather than muscle deficiency(21,22). Weight loss (~10%) induced by energy restriction resulted in slightly higher lean mass loss than the CCI (8.4% appendicular lean mass and 7.6% total lean mass loss at 20 weeks)(23). Total lean mass loss from 10% weight reduction by bariatric surgery is reported in the range of 7.3 to 15.9% from baseline(24,25). Greater weight loss is usually associated with more lean mass loss(26–28). Approximately 25% of diet-induced weight loss (without exercise) often arises from FFM(29). In the present intervention, FFM loss contributed an estimated 14% to the lower extremity weight loss. The lower proportion of FFM loss in the CCI group, despite higher percentage of weight loss, may be due to the adequate dietary protein recommendations (30,31). Since ~73% of FFM is water, the observed reduction of LELM in the first year of intervention may have arisen from natriuresis and water loss during keto-adaptation(32,33).

**Supplementary Table 1.** Equations for calculating HOMA-IR (insulin-derived), HOMA-IR (c-peptide derived), LDL-cholesterol, NAFLD liver fat score (NLF) and NAFLD fibrosis score (NFS)

| Score | Equation |
| --- | --- |
| HOMA-IR (insulin derived) | [fasting insulin (mU/L) x fasting glucose (mg/dL)]/ 405 |
| HOMA-IR (c-peptide derived) | Calculation performed using spreadsheet downloaded from<br>http://www.dtu.ox.ac.uk/homacalculator/ |
| Friedewald LDL-cholesterol | total cholesterol (mg/dL) - HDL cholesterol (mg/dL) - [TG (mg/dL)/5] |
| NAFLD liver fat score (N-LFS) | -2.89 + 1.18 x metabolic syndrome (yes=1 or no=0) + 0.45 x type 2 diabetes<br>(yes=2 or no=0)* + 0.15 x fasting insulin (mU/l) + 0.04 x fasting serum AST (U/L) –<br>0.94 x AST/ALT |
| NAFLD fibrosis score (NFS) | -1.675 + 0.037 × Age (yrs) + 0.094 × BMI (kg/m<sup>2</sup>) + 1.13 × IFG/diabetes (yes = 1,<br>no = 0) + 0.99 × AST/ALT ratio – 0.013 × Platelet (×10<sup>9</sup>/L) –0.66 × Albumin (g/dl) |

**Supplementary Table 2.**
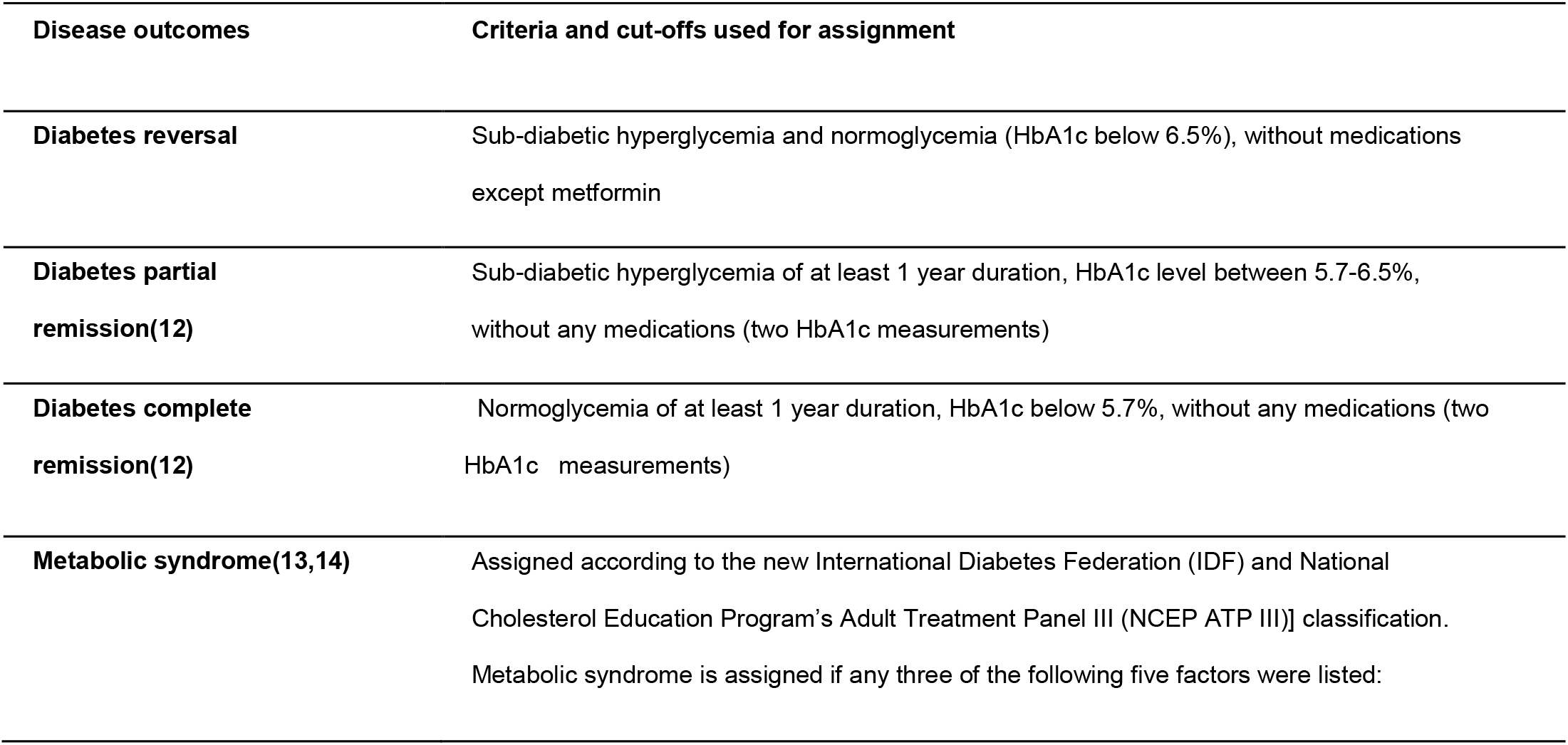
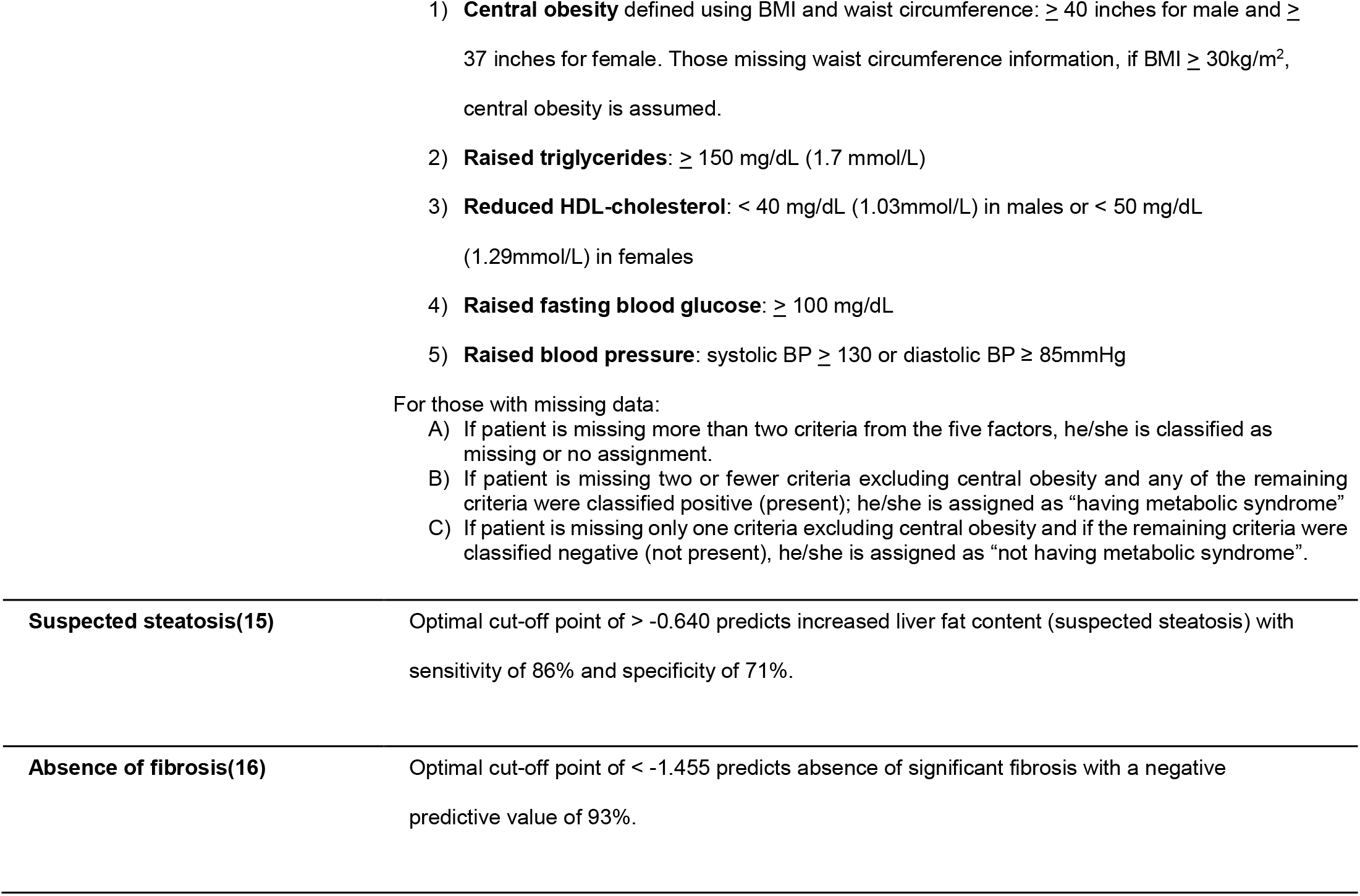
Criteria and cut-offs for diabetes reversal, diabetes partial remission, diabetes complete remission, metabolic syndrome, steatosis and absence of fibrosis

**Supplementary Table 3.**
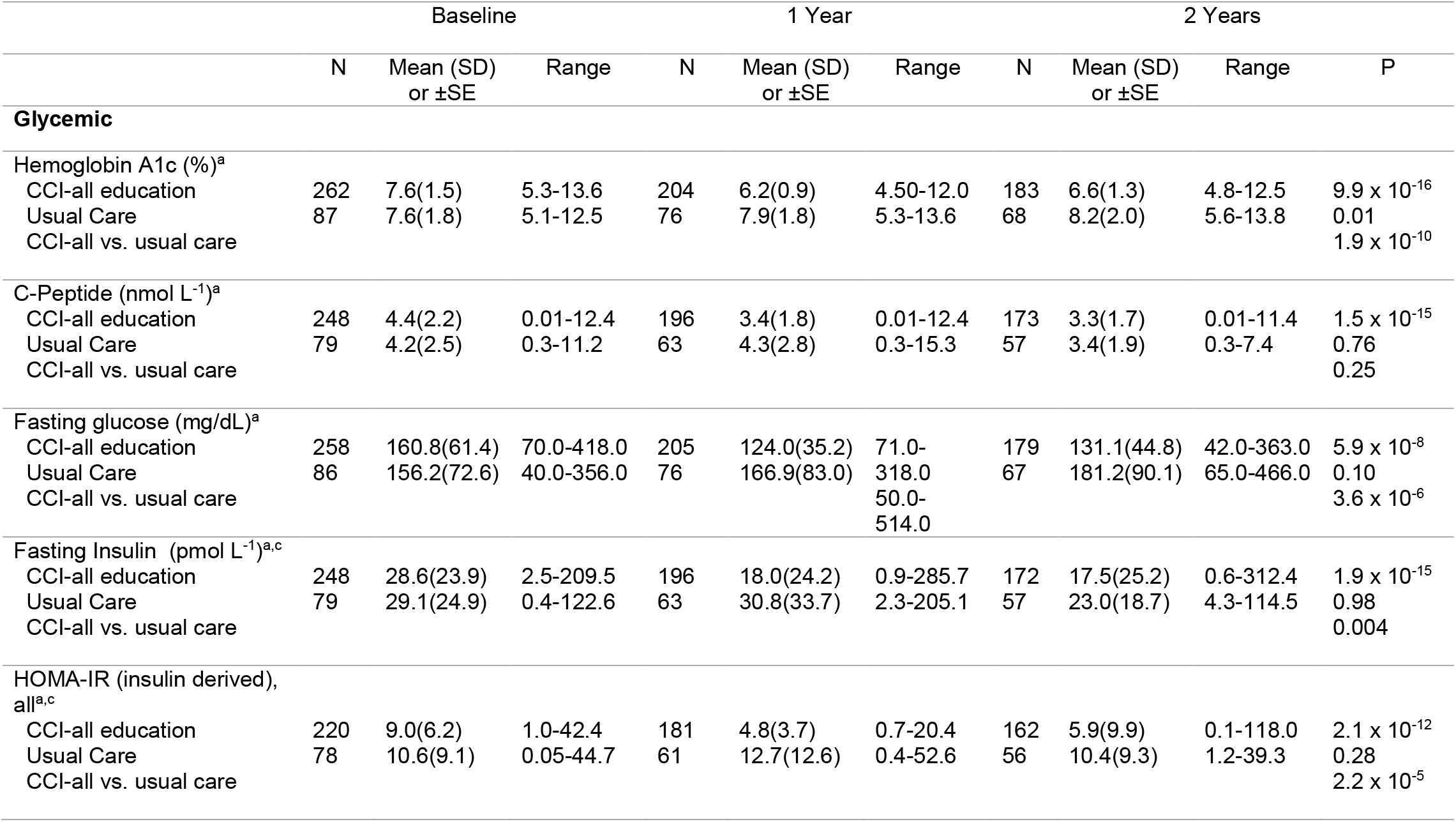
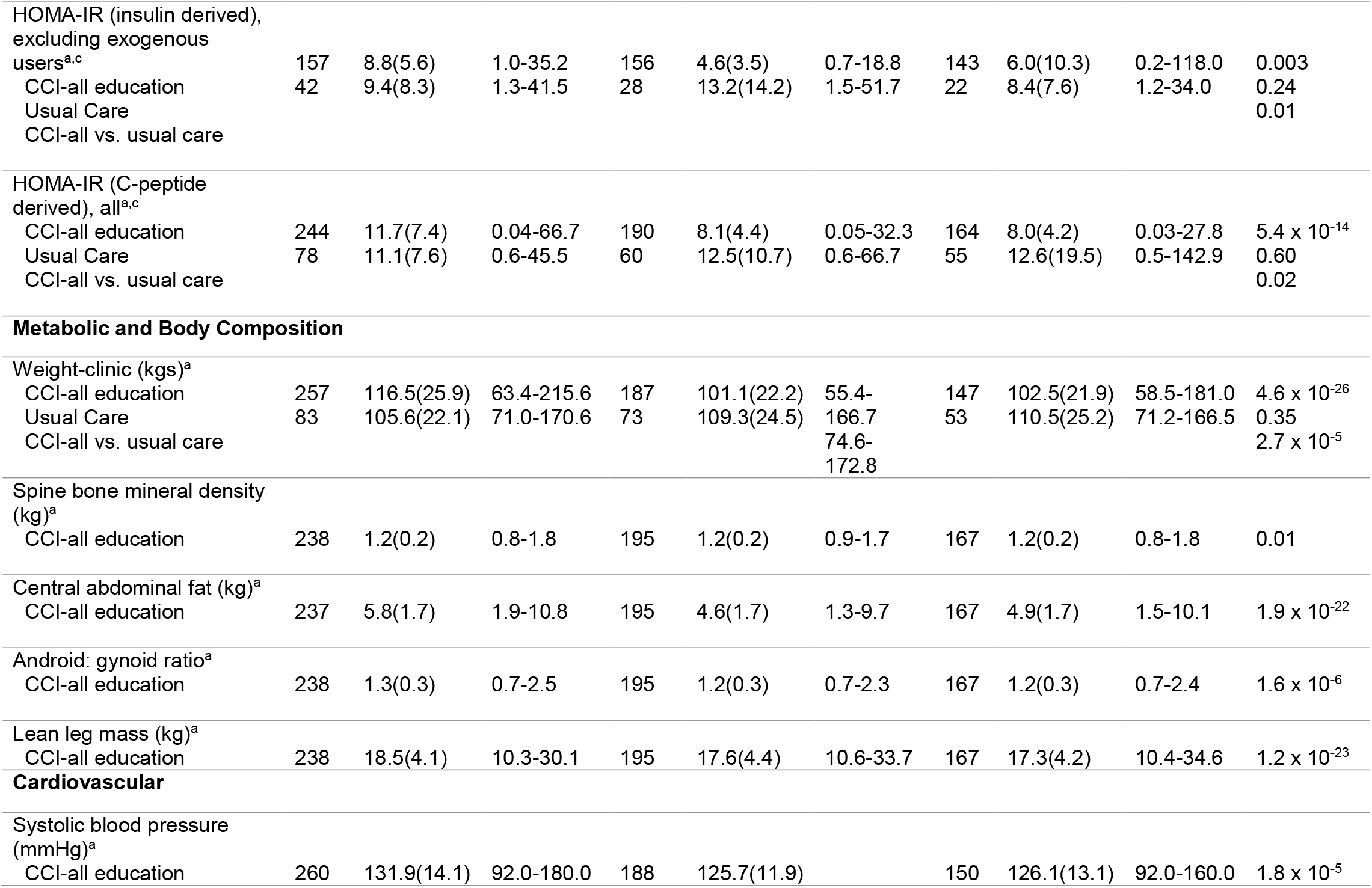
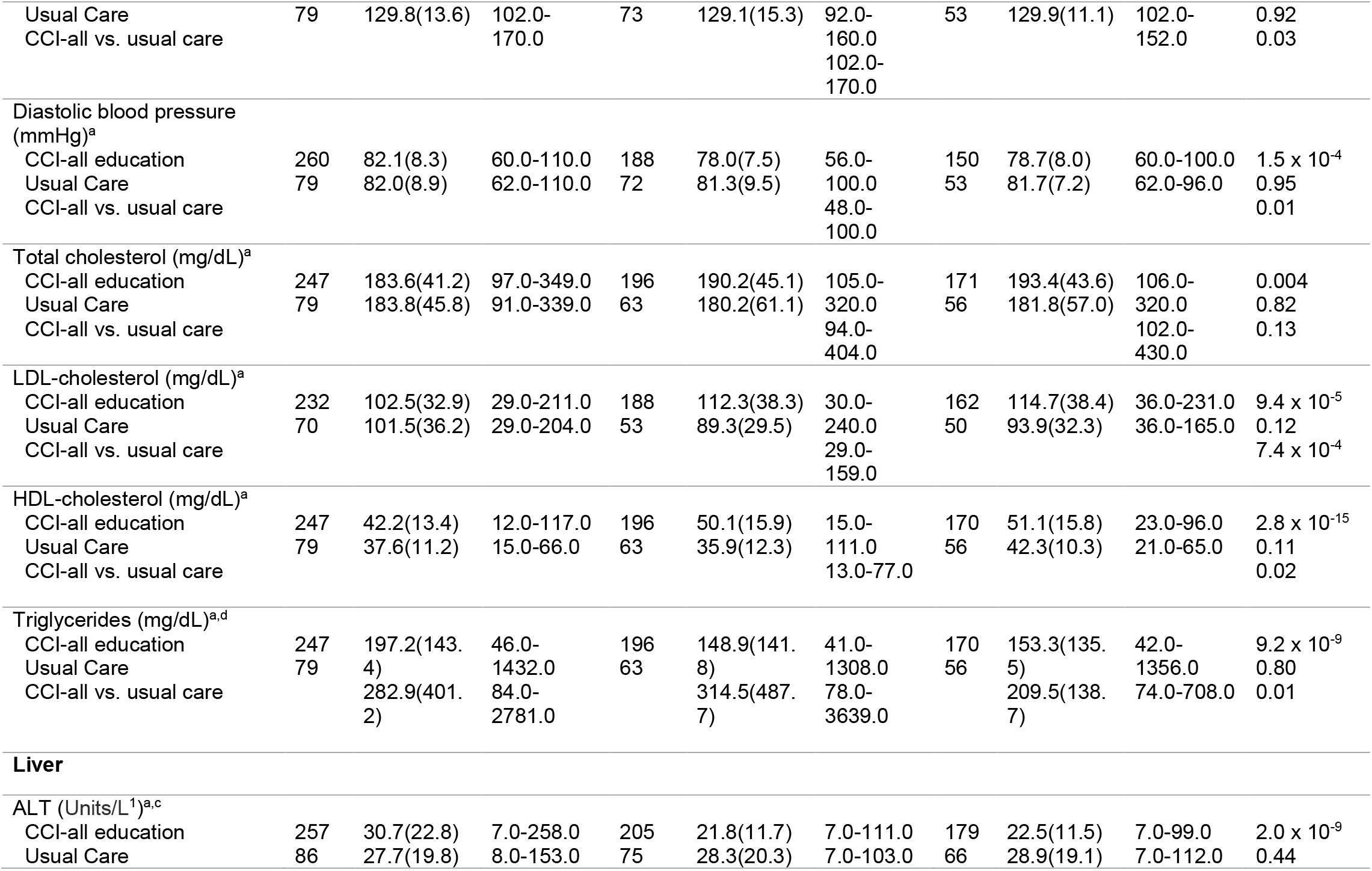
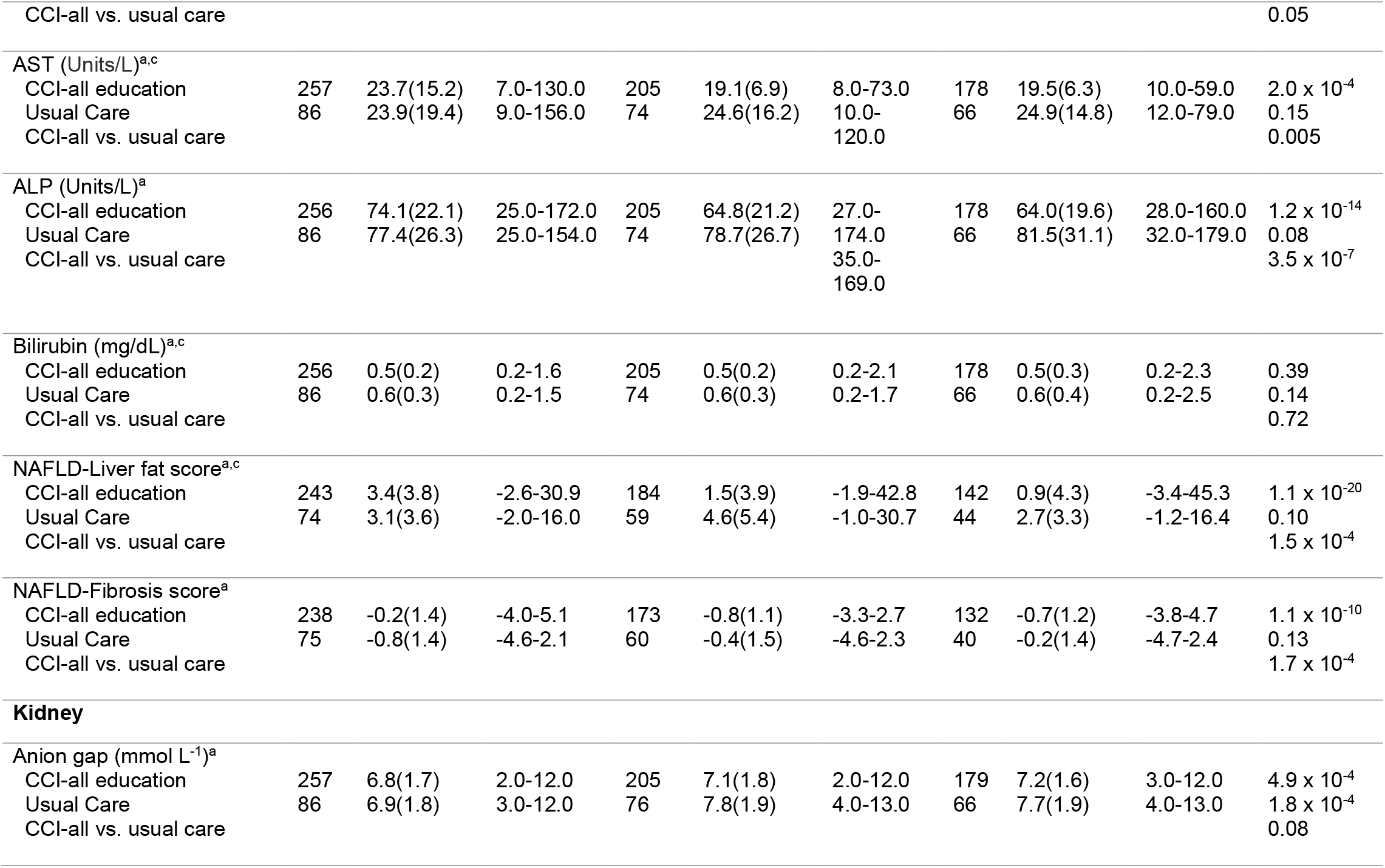
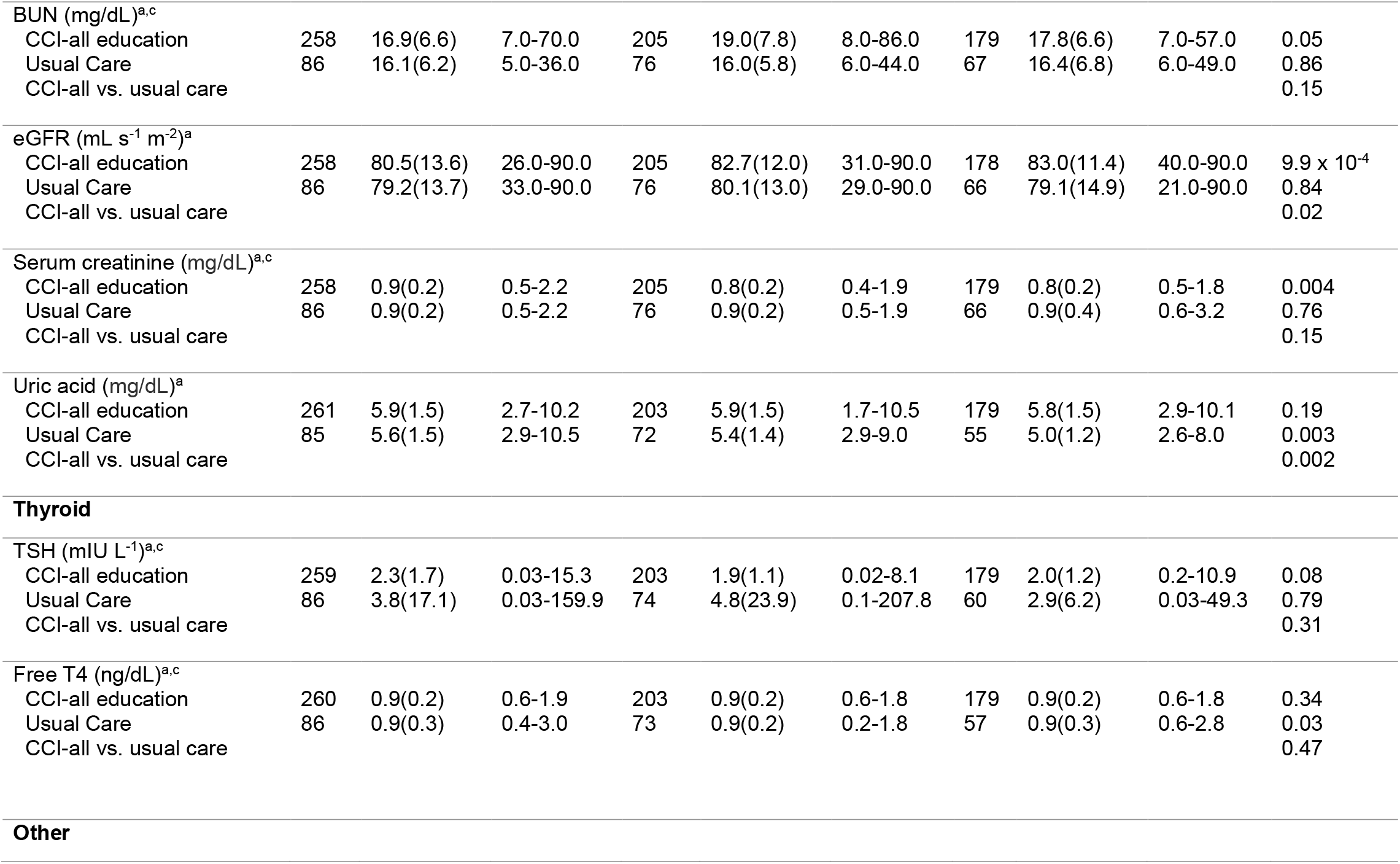
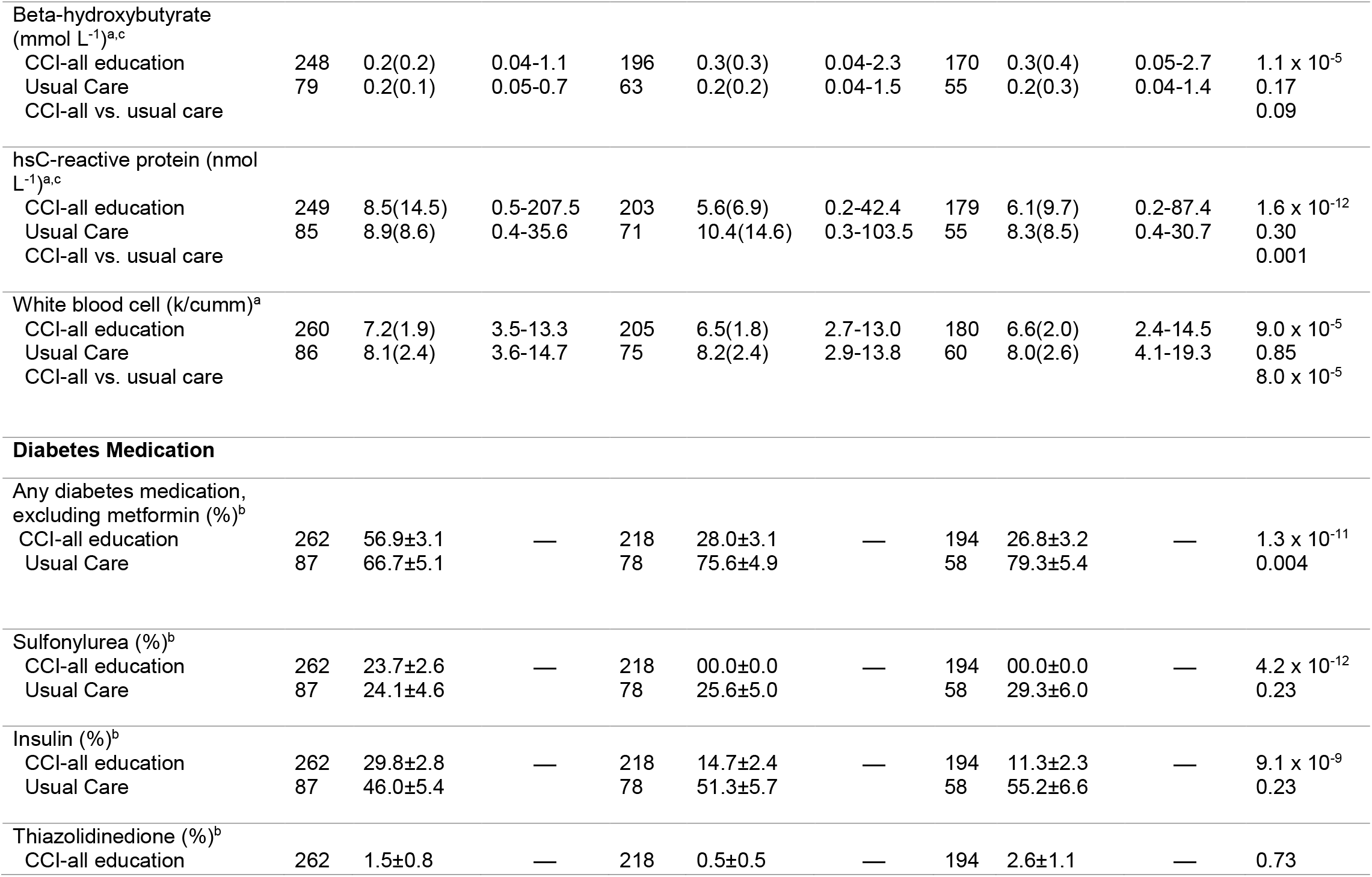
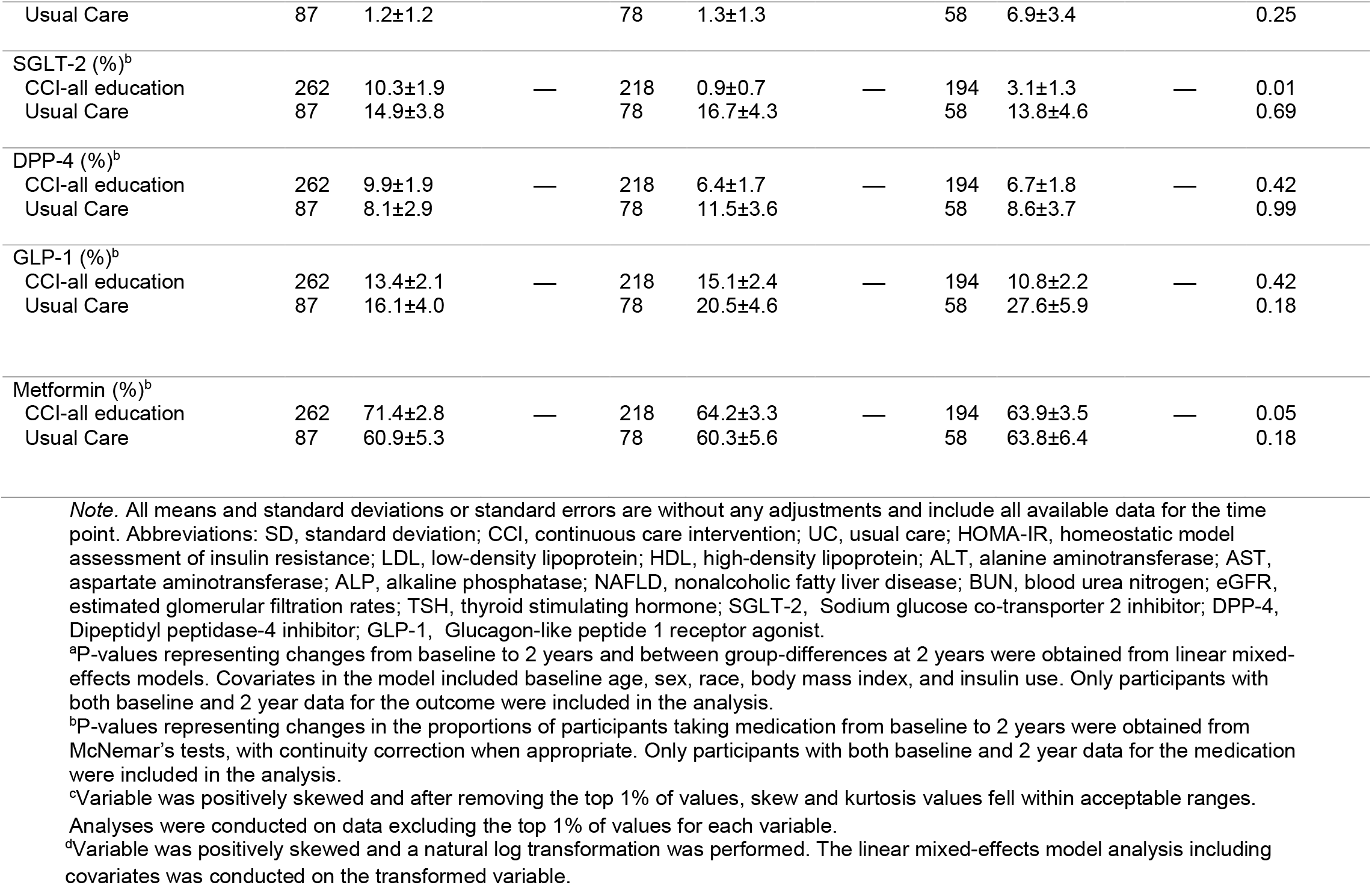
Descriptives and results of completer-only analyses

**Supplementary Table 4.** Disease outcomes in CCI and UC participants after 2 years (Intent-to-treat analysis with imputation)

| Disease Outcomes | Continuous Care Intervention<br>(n=262) |  |  | Usual Care (n=87) |  |  | Between<br>group |
| --- | --- | --- | --- | --- | --- | --- | --- |
|  | Baseline | 2 Years | P | Baseline | 2 Years | P | P |
| **Diabetes Reversal (%)** | 12.1±2.0 | 53.5±3.4 | <0.0x10<sup>-36</sup> | 16.4±4.5 | 9.3±3.9 | 0.04 | <0.0x10<sup>-36</sup> |
| **Diabetes Remission (%)<sup>a</sup>** | — | 17.6±2.5 | — | — | 2.4±1.7 | — | 5.1x10<sup>-9</sup> |
| **Complete Remission (%)** | — | 6.7±1.6 | — | — | 0.0±0.0 | — | 1.1x10<sup>-5</sup> |
| **Metabolic Syndrome (%)** | 89.1±2.0 | 61.9±4.0 | 4.9x10<sup>-15</sup> | 92.4±3.3 | 85.9±5.1 | 0.24 | 4.7x10<sup>-7</sup> |
| **Suspected Steatosis (%)** | 95.8±1.4 | 67.4±4.2 | <0.0x10<sup>-36</sup> | 94.7±3.0 | 89.0±5.1 | 0.16 | 2.5x10<sup>-7</sup> |
| **Absence of Fibrosis (%)** | 18.3±2.5 | 30.8±4.0 | 1.4x10<sup>-5</sup> | 24.9±5.4 | 15.9±5.8 | 0.08 | 4x10<sup>-3</sup> |

*Note.* Percentages and standard errors are provided. Estimates were obtained from generalized estimating equation models which provide adjusted proportions, controlling for baseline age, sex, race, time since diagnosis, body mass index, and insulin use. Multiple imputation was used to replace missing values, facilitating intent-to-treat analyses. A significance level of P<0.0012 ensures overall simultaneous significance of P < 0.05 over the 43 study variables using Bonferroni correction.
<sup>a</sup>Diabetes remission includes both partial and complete remission.

**Supplementary Table 5.** Disease outcomes in CCI and UC participants after 2 years (Completers-only analysis)

| Disease Outcomes | Continuous Care Intervention |  |  |  |  | Usual Care (n=87) |  |  |  |  | Between group |
| --- | --- | --- | --- | --- | --- | --- | --- | --- | --- | --- | --- |
|  | N | Baseline | N | 2 Years | P | N | Baseline | N | 2 Years | P | P |
| **Diabetes Reversal (%)** | 262 | 12.2±2.0 | 181 | 54.7±3.7 | <0.0x10<sup>-36</sup> | 87 | 20.7±4.4 | 57 | 10.5±4.1 | 0.07 | 5.4x10<sup>-15</sup> |
| **Diabetes Remission (%)<sup>a</sup>** | — | — | 208 | 18.8±2.7 | — | — | — | 79 | 2.5±1.8 | — | 1.6x10<sup>-8</sup> |
| **Complete Remission (%)** | — | — | 210 | 6.7±1.7 | — | — | — | 81 | 0.0±0.0 | — | 1.1x10<sup>-4</sup> |
| **Metabolic Syndrome (%)** | 262 | 88.6±2.0 | 154 | 63.0±3.9 | 9.9x10<sup>-11</sup> | 81 | 91.4±3.1 | 54 | 87.0±4.6 | 0.51 | 8.9x10<sup>-5</sup> |
| **Suspected Steatosis (%)** | 243 | 96.3±1.2 | 142 | 67.6±3.9 | 7.7x10<sup>-13</sup> | 74 | 94.6±2.6 | 44 | 88.6±4.8 | 0.40 | 3.9x10<sup>-5</sup> |
| **Absence of Fibrosis (%)** | 238 | 18.1±2.5 | 132 | 29.6±4.0 | 0.003 | 75 | 28.0±5.2 | 40 | 17.5±6.1 | 0.58 | 0.09 |

*Note.* All percentages and standard errors are without any adjustments and include all available data for the time point. P-values representing within-group changes from baseline to 2 years and between-group differences at 2 years were obtained from generalized estimating equation models. Covariates in the model included baseline age, sex, race, time since diagnosis, body mass index, and insulin use. Only participants with both baseline and 2 year data for the outcome were included in the analysis. A significance level of $P < 0.0012$ ensures overall simultaneous significance of $P < 0.05$ over the 43 study variables using Bonferroni correction.
<sup>a</sup>Diabetes remission includes both partial and complete remission.

### Supplementary Figures Legend

**Supplementary Figure 1.**
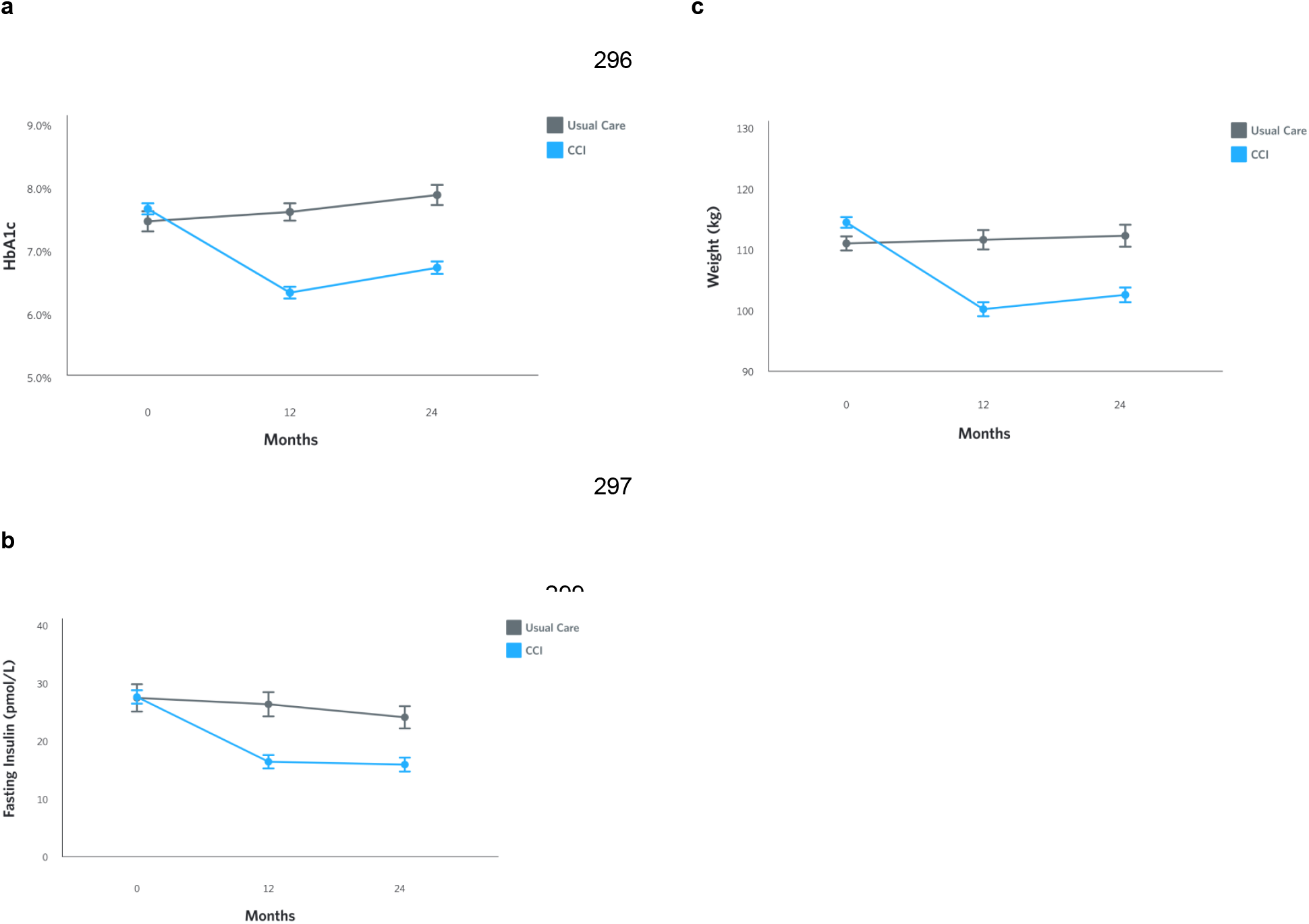
Adjusted mean changes (CCI versus UC) from baseline to 2-years in (A) HbA1c, (B) Fasting insulin, (C) Weight.

**Supplementary Figure 2.**
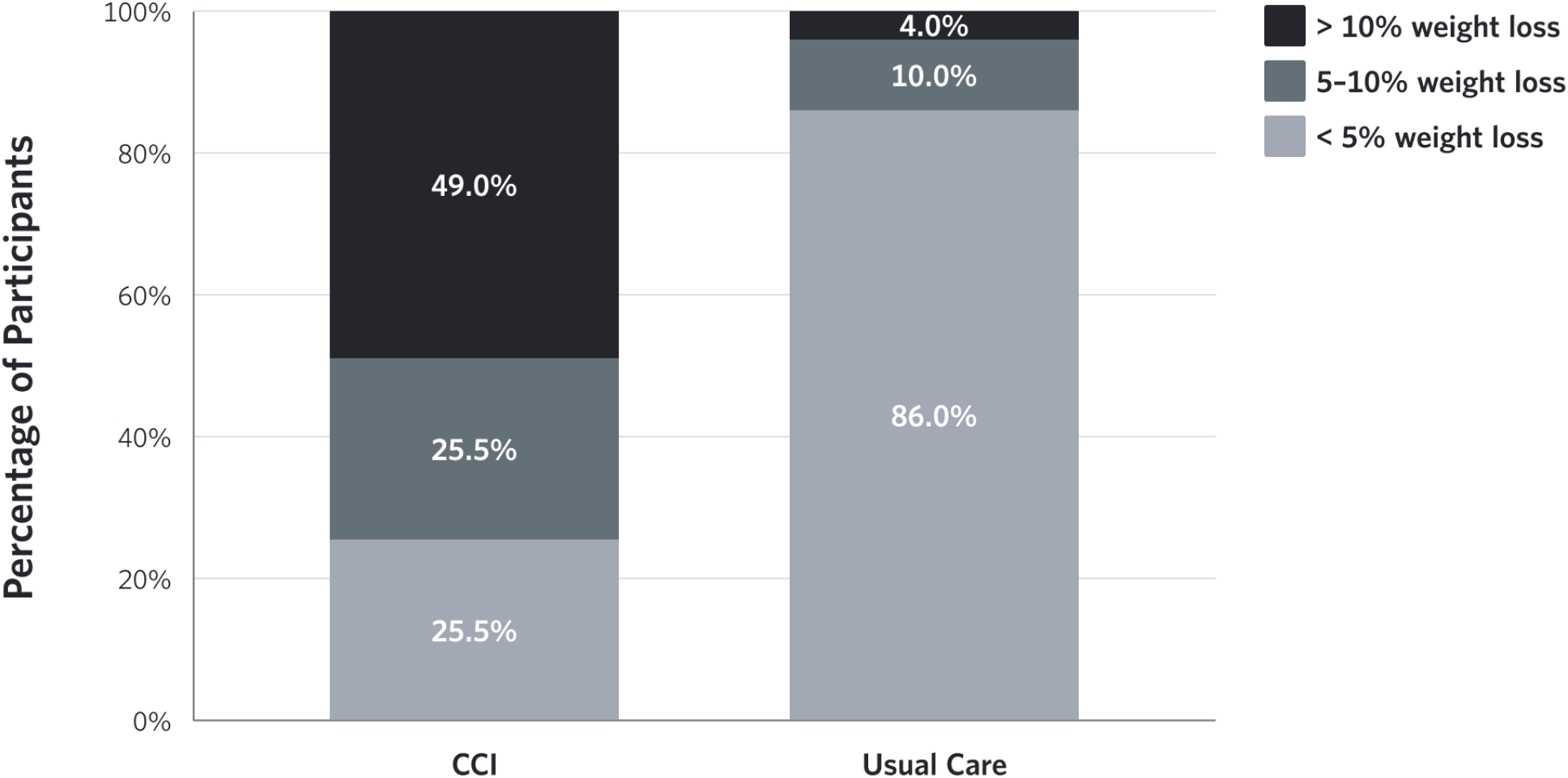
Stratification of participants based on weight change (%) categories in each intervention groups, UC and CCI, among completers. Category <5% includes participants with weight gain.

**Supplementary Figure 3.**
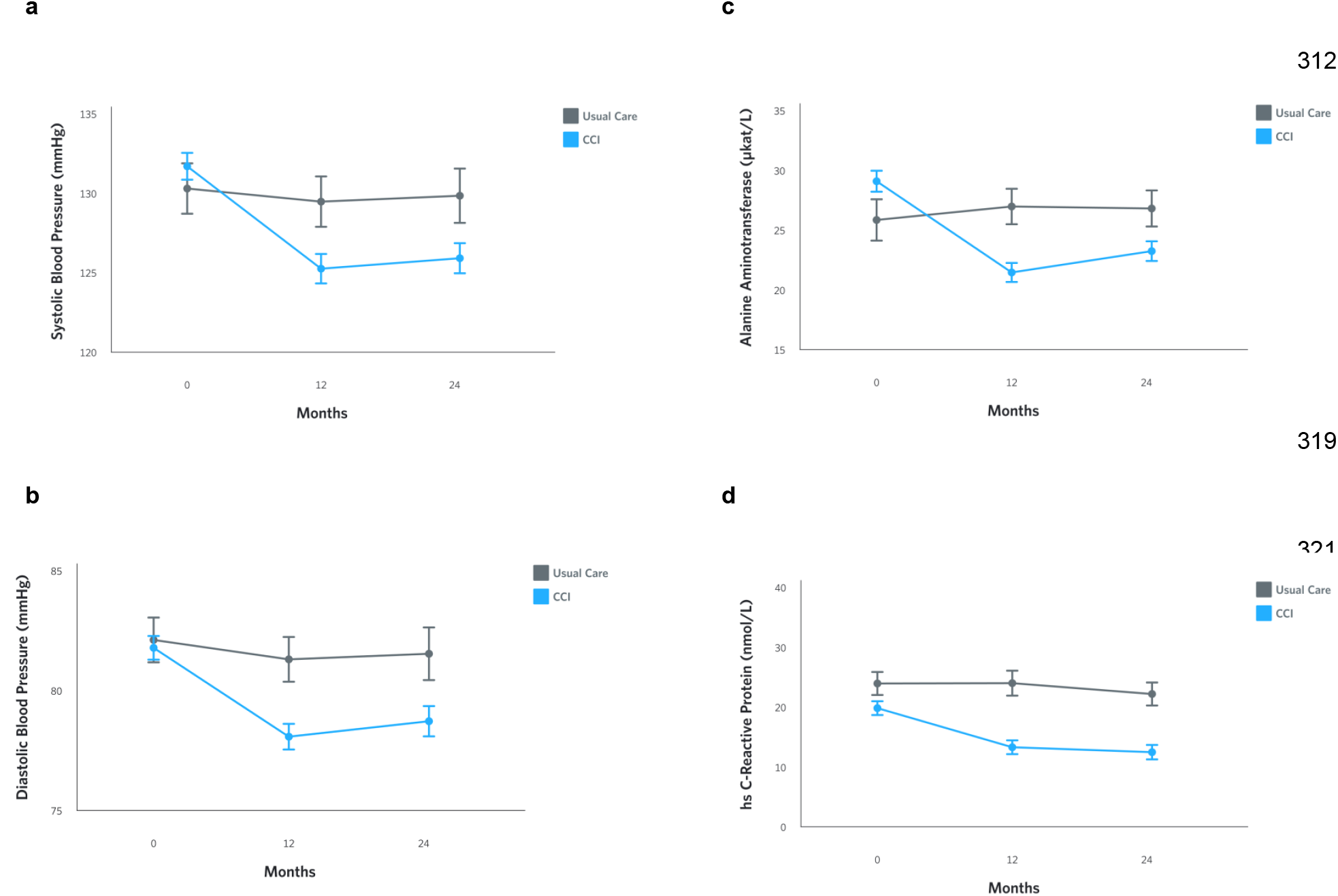
Adjusted mean changes (CCI versus UC) from baseline to 2-years in (A) Systolic Blood Pressure, (B) Diastolic Blood Pressure, (C) Alanine aminotransferase (ALT), and (D) High sensitive C-reactive protein (hsCRP).

**Supplementary Figure 4.**
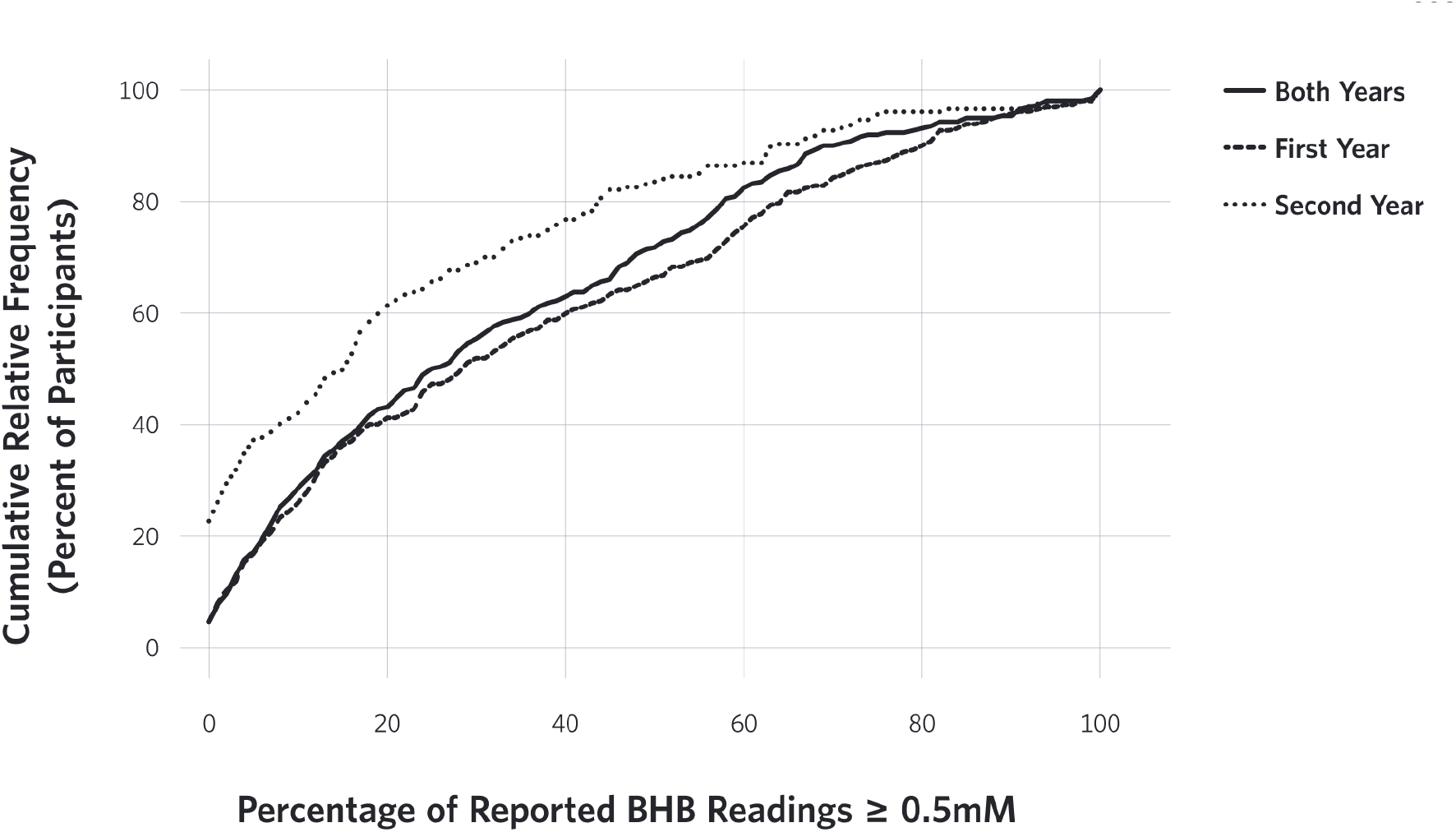
Cumulative relative frequency (%) of percentage participants reporting BHB > 0.5mM at first, second and both years of the study. The differences in the distribution of participants reporting BHB > 0.5mM between one and two years are illustrated in the figure.

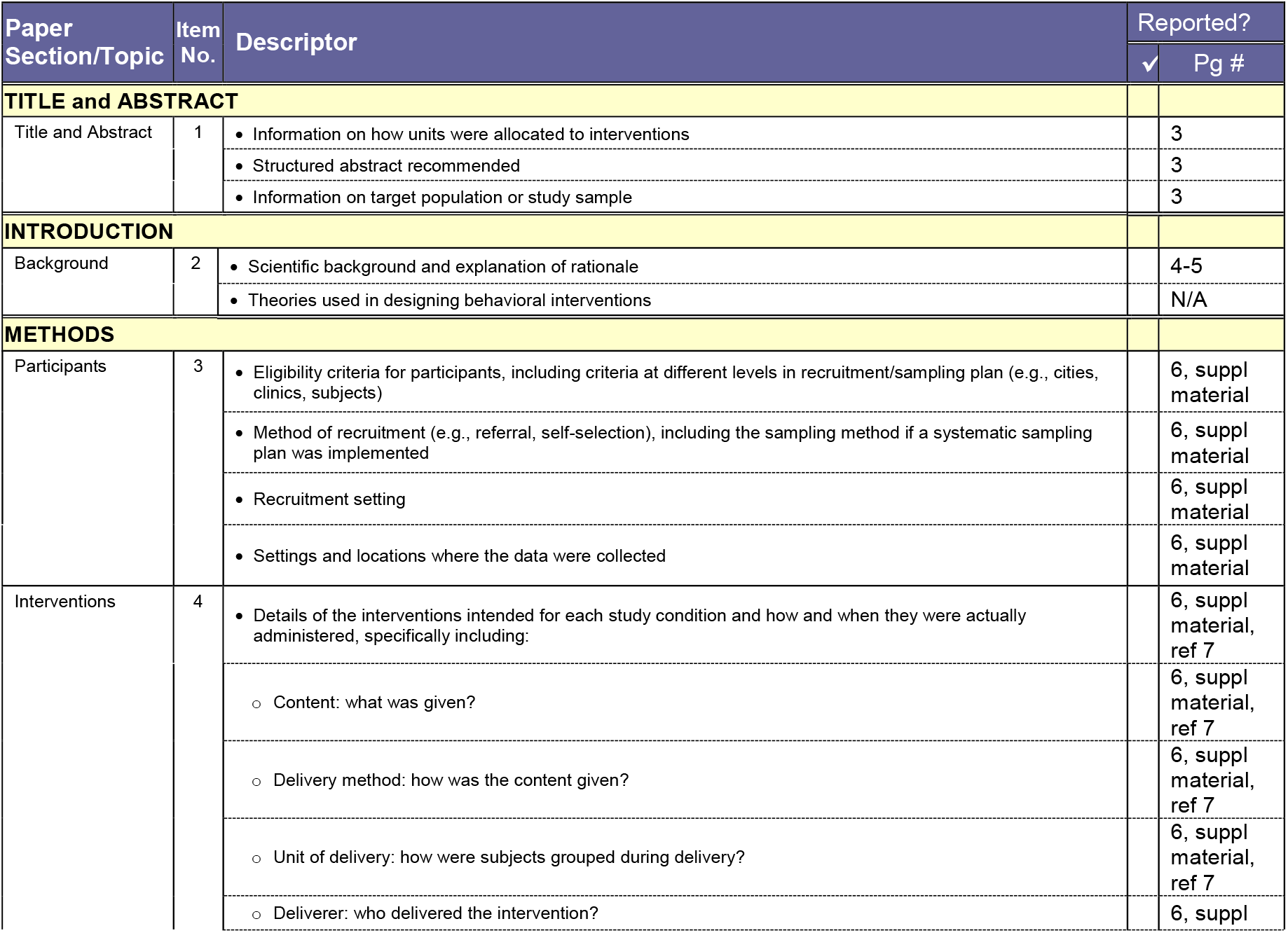
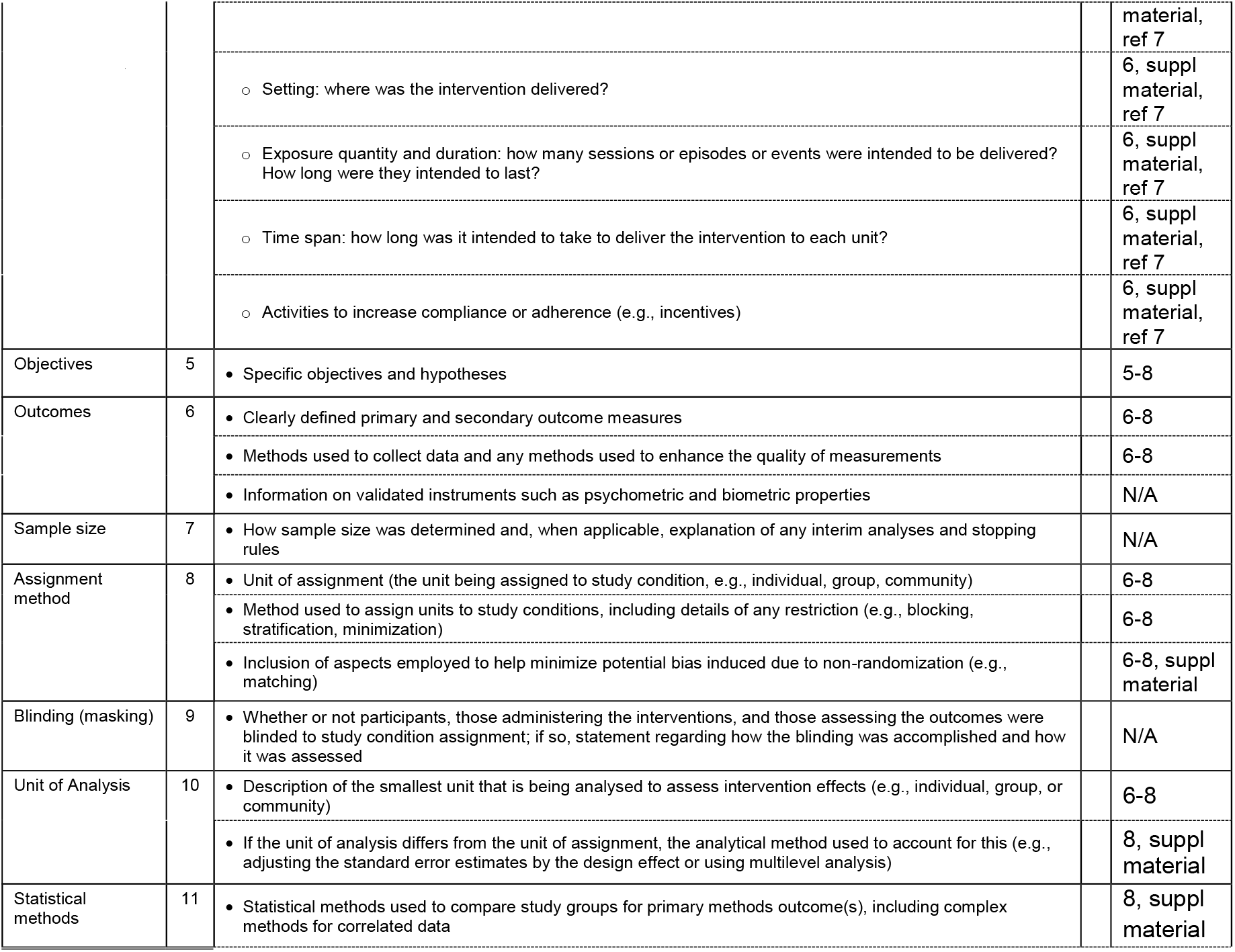
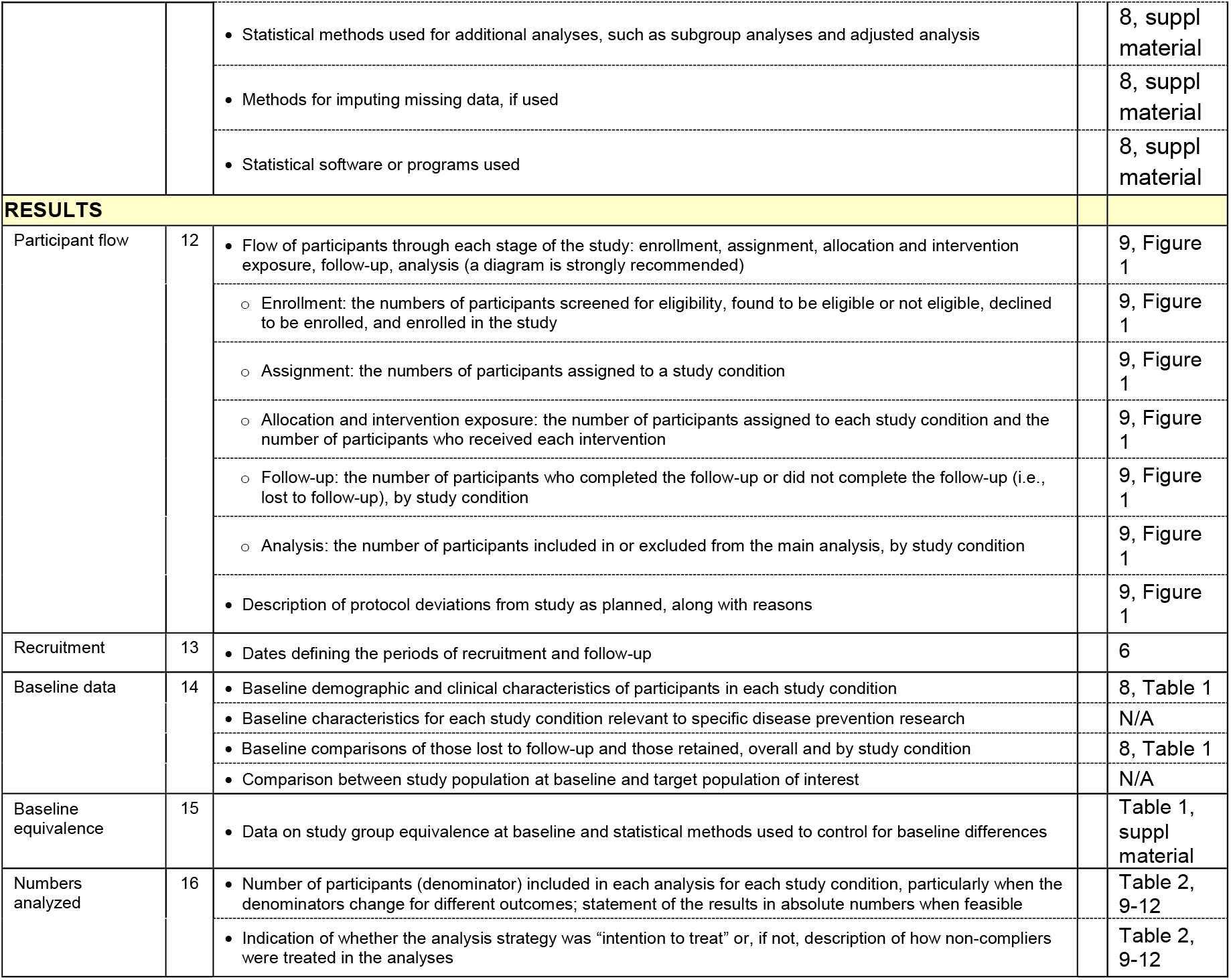
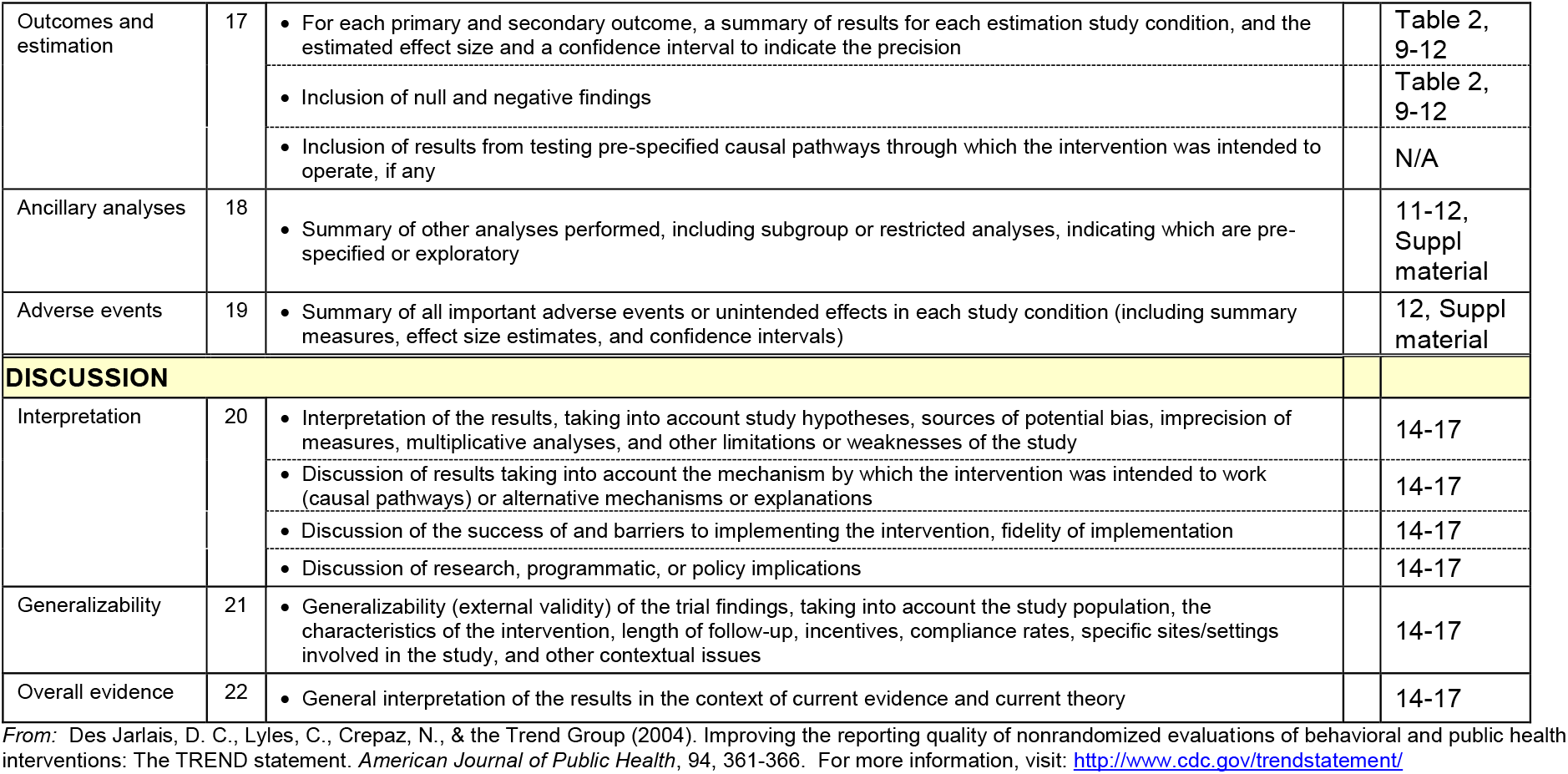
TREND Statement Checklist

| Paper<br>Section/Topic | Item<br>No. | Descriptor | Reported? |  |
| --- | --- | --- | --- | --- |
|  |  |  | ✓ | Pg # |
| TITLE and ABSTRACT |  |  |  |  |
| Title and Abstract | 1 | • Information on how units were allocated to interventions |  | 3 |
|  |  | • Structured abstract recommended |  | 3 |
|  |  | • Information on target population or study sample |  | 3 |
| INTRODUCTION |  |  |  |  |
| Background | 2 | • Scientific background and explanation of rationale |  | 4-5 |
|  |  | • Theories used in designing behavioral interventions |  | N/A |
| METHODS |  |  |  |  |
| Participants | 3 | • Eligibility criteria for participants, including criteria at different levels in recruitment/sampling plan (e.g., cities, clinics, subjects) |  | 6, suppl material |
|  |  | • Method of recruitment (e.g., referral, self-selection), including the sampling method if a systematic sampling plan was implemented |  | 6, suppl material |
|  |  | • Recruitment setting |  | 6, suppl material |
|  |  | • Settings and locations where the data were collected |  | 6, suppl material |
| Interventions | 4 | • Details of the interventions intended for each study condition and how and when they were actually administered, specifically including: |  | 6, suppl material, ref 7 |
|  |  | ○ Content: what was given? |  | 6, suppl material, ref 7 |
|  |  | ○ Delivery method: how was the content given? |  | 6, suppl material, ref 7 |
|  |  | ○ Unit of delivery: how were subjects grouped during delivery? |  | 6, suppl material, ref 7 |
|  |  | ○ Deliverer: who delivered the intervention? |  | 6, suppl |

|  |  |  |  |
| --- | --- | --- | --- |
|  |  |  | material, ref 7 |
|  |  | o Setting: where was the intervention delivered? | 6, suppl material, ref 7 |
|  |  | o Exposure quantity and duration: how many sessions or episodes or events were intended to be delivered? How long were they intended to last? | 6, suppl material, ref 7 |
|  |  | o Time span: how long was it intended to take to deliver the intervention to each unit? | 6, suppl material, ref 7 |
|  |  | o Activities to increase compliance or adherence (e.g., incentives) | 6, suppl material, ref 7 |
| Objectives | 5 | • Specific objectives and hypotheses | 5-8 |
| Outcomes | 6 | • Clearly defined primary and secondary outcome measures | 6-8 |
|  |  | • Methods used to collect data and any methods used to enhance the quality of measurements | 6-8 |
|  |  | • Information on validated instruments such as psychometric and biometric properties | N/A |
| Sample size | 7 | • How sample size was determined and, when applicable, explanation of any interim analyses and stopping rules | N/A |
| Assignment method | 8 | • Unit of assignment (the unit being assigned to study condition, e.g., individual, group, community) | 6-8 |
|  |  | • Method used to assign units to study conditions, including details of any restriction (e.g., blocking, stratification, minimization) | 6-8 |
|  |  | • Inclusion of aspects employed to help minimize potential bias induced due to non-randomization (e.g., matching) | 6-8, suppl material |
| Blinding (masking) | 9 | • Whether or not participants, those administering the interventions, and those assessing the outcomes were blinded to study condition assignment; if so, statement regarding how the blinding was accomplished and how it was assessed | N/A |
| Unit of Analysis | 10 | • Description of the smallest unit that is being analysed to assess intervention effects (e.g., individual, group, or community) | 6-8 |
|  |  | • If the unit of analysis differs from the unit of assignment, the analytical method used to account for this (e.g., adjusting the standard error estimates by the design effect or using multilevel analysis) | 8, suppl material |
| Statistical methods | 11 | • Statistical methods used to compare study groups for primary methods outcome(s), including complex methods for correlated data | 8, suppl material |
|  |  | • Statistical methods used for additional analyses, such as subgroup analyses and adjusted analysis | 8, suppl material |
|  |  | • Methods for imputing missing data, if used | 8, suppl material |
|  |  | • Statistical software or programs used | 8, suppl material |
| RESULTS |  |  |  |
| Participant flow | 12 | • Flow of participants through each stage of the study: enrollment, assignment, allocation and intervention exposure, follow-up, analysis (a diagram is strongly recommended) | 9, Figure 1 |
|  |  | ○ Enrollment: the numbers of participants screened for eligibility, found to be eligible or not eligible, declined to be enrolled, and enrolled in the study | 9, Figure 1 |
|  |  | ○ Assignment: the numbers of participants assigned to a study condition | 9, Figure 1 |
|  |  | ○ Allocation and intervention exposure: the number of participants assigned to each study condition and the number of participants who received each intervention | 9, Figure 1 |
|  |  | ○ Follow-up: the number of participants who completed the follow-up or did not complete the follow-up (i.e., lost to follow-up), by study condition | 9, Figure 1 |
|  |  | ○ Analysis: the number of participants included in or excluded from the main analysis, by study condition | 9, Figure 1 |
|  |  | • Description of protocol deviations from study as planned, along with reasons | 9, Figure 1 |
| Recruitment | 13 | • Dates defining the periods of recruitment and follow-up | 6 |
| Baseline data | 14 | • Baseline demographic and clinical characteristics of participants in each study condition | 8, Table 1 |
|  |  | • Baseline characteristics for each study condition relevant to specific disease prevention research | N/A |
|  |  | • Baseline comparisons of those lost to follow-up and those retained, overall and by study condition | 8, Table 1 |
|  |  | • Comparison between study population at baseline and target population of interest | N/A |
| Baseline equivalence | 15 | • Data on study group equivalence at baseline and statistical methods used to control for baseline differences | Table 1, suppl material |
| Numbers analyzed | 16 | • Number of participants (denominator) included in each analysis for each study condition, particularly when the denominators change for different outcomes; statement of the results in absolute numbers when feasible | Table 2, 9-12 |
|  |  | • Indication of whether the analysis strategy was “intention to treat” or, if not, description of how non-compliers were treated in the analyses | Table 2, 9-12 |
| Outcomes and estimation | 17 | • For each primary and secondary outcome, a summary of results for each estimation study condition, and the estimated effect size and a confidence interval to indicate the precision | Table 2, 9-12 |
|  |  | • Inclusion of null and negative findings | Table 2, 9-12 |
|  |  | • Inclusion of results from testing pre-specified causal pathways through which the intervention was intended to operate, if any | N/A |
| Ancillary analyses | 18 | • Summary of other analyses performed, including subgroup or restricted analyses, indicating which are pre-specified or exploratory | 11-12, Suppl material |
| Adverse events | 19 | • Summary of all important adverse events or unintended effects in each study condition (including summary measures, effect size estimates, and confidence intervals) | 12, Suppl material |
| **DISCUSSION** |  |  |  |
| Interpretation | 20 | • Interpretation of the results, taking into account study hypotheses, sources of potential bias, imprecision of measures, multiplicative analyses, and other limitations or weaknesses of the study | 14-17 |
|  |  | • Discussion of results taking into account the mechanism by which the intervention was intended to work (causal pathways) or alternative mechanisms or explanations | 14-17 |
|  |  | • Discussion of the success of and barriers to implementing the intervention, fidelity of implementation | 14-17 |
|  |  | • Discussion of research, programmatic, or policy implications | 14-17 |
| Generalizability | 21 | • Generalizability (external validity) of the trial findings, taking into account the study population, the characteristics of the intervention, length of follow-up, incentives, compliance rates, specific sites/settings involved in the study, and other contextual issues | 14-17 |
| Overall evidence | 22 | • General interpretation of the results in the context of current evidence and current theory | 14-17 |

From: Des Jarlais, D. C., Lyles, C., Crepaz, N., & the Trend Group (2004). Improving the reporting quality of nonrandomized evaluations of behavioral and public health interventions: The TREND statement. *American Journal of Public Health*, 94, 361-366. For more information, visit: http://www.cdc.gov/trendstatement/

Author contributions

S.J.A, R.N.A, and J.P.M drafted the manuscript. S.J.A, R.N.A, A.L.M, N.H.B, and S.J.H participated in data acquisition and compiling. R.N.A and S.J.A analyzed the data. J.P.M, A.L.M, N.H.B, W.W.C, R.N.A, S.J.A, S.J.H, S.D.P and J.S.V edited the manuscript. All authors approved the final version of the manuscript.

## Abbreviations

CCI: continuous care intervention;
UC: usual care;
T2D: type 2 diabetes;
HbA1c: hemoglobin A1c;
CVD: cardiovascular disease;
VLCD: very low calorie diet;
BMI: body mass index;
BHB: beta-hydroxybutryrate;
BMD: bone mineral density;
CAF: central abdominal fat;
A/G: android:gynoid ratio;
LELM: lower extremities lean mass;
HDL: high density lipoprotein;
LDL: low density lipoprotein;
ALT: alanine aminotransferase;
AST: aspartate aminotransferase;
ALP: alkaline phosphatase;
NAFLD: nonalcoholic fatty liver disease;
NLF: NAFLD liver fat score;
NFS: NAFLD fibrosis score;
TSH: thyroid stimulating hormone;
BUN: blood urea nitrogen;
eGFR: estimated glomerular filtration rate;
hsCRP: high sensitive C-reactive protein;
WBC: white blood cells;
HOMA-IR: Homeostatic Model Assessment of Insulin Resistance;
SGLT-2: sodium-glucose cotransporter-2 inhibitors;
DPP-4: dipeptidyl peptidase-4 inhibitors;
GLP-1: glucagon-like-peptide 1 receptor agonists;
FFM: fat-free mass;
VAT: visceral adipose tissue;
GLM: generalized linear model;
LMM: linear mixed-effect model;
ADA: American Diabetes Association;
CLIA: Clinical Laboratory Improvement Amendments;
IRB: Institutional Review Board;
DXA: dual-energy X-ray absorptiometry

